# ENHANCED CLEAVAGE OF GENOMIC *CCR5* USING CASX2^Max^

**DOI:** 10.1101/2025.07.08.663680

**Authors:** Christine A. Hodge, Niles P. Donegan, David A. Armstrong, Mathew S. Hayden, Alexandra L. Howell

## Abstract

Development of novel CRISPR/Cas systems enhances opportunities for gene editing to treat infectious diseases, cancer, and genetic disorders. We evaluated CasX2 (*Plm*Cas12e), a class II CRISPR system derived from *Planctomycetes*, a non-pathogenic bacterium present in aquatic and terrestrial soils. CasX2 offers several advantages over *Streptococcus pyogenes* Cas9 (*Sp*Cas9) and *Staphylococcus aureus* Cas9 (*Sa*Cas9), including its smaller size, distinct protospacer adjacent motif (PAM) requirements, staggered cleavage cuts that promote homology-directed repair, and no known pre-existing immunity in humans. A recent study reported that a three amino acid substitution in CasX2 significantly enhanced cleavage activity (1). Therefore, we compared cleavage efficiency and double-stranded break repair characteristics between the native CasX2 and the variant, CasX2^Max^, for cleavage of *CCR5*, a gene that encodes the CCR5 receptor important for HIV-1 infection. Two CasX2 single guide RNAs (sgRNAs) were designed that flanked the 32 bases deleted in the natural *CCR5 Δ32* mutation. Nanopore sequencing demonstrated that CasX2 using sgRNAs with spacers of 17 nucleotides (nt), 20 nt or 23 nt in length were ineffective at cleaving genomic *CCR5.* In contrast, CasX2^Max^ using sgRNAs with 20 nt and 23 nt spacer lengths, enabled robust genomic cleavage of *CCR5*. Structural modeling indicated that two of the CasX2^Max^ substitutions enhanced sgRNA-DNA duplex stability, while the third improved DNA strand alignment within the catalytic site. These structural changes likely underlie the increased activity of CasX2^Max^ in cellular gene excision. In sum, CasX2^Max^ consistently outperformed native CasX2 across all assays and represents a superior gene-editing platform for therapeutic applications.

## Introduction

The ability to selectively modify a DNA sequence by excising or replacing one or more nucleotides presents us with a powerful opportunity to treat and potentially cure a wide range of genetically-based diseases. Gene editing using the CRISPR/Cas approach enables precise alterations to specific genomic regions through the combined action of a short RNA sequence, termed a guide RNA (gRNA), and an endonuclease termed Cas (CRISPR-associated). Multiple CRISPR/Cas systems have been adapted for gene editing in eukaryotic cells, each differing in the target recognition requirements of the Cas endonuclease and the design of the corresponding gRNA to recognize target regions in the DNA. In all CRISPR/Cas systems, the Cas enzyme is invariant while the gRNAs contain a spacer sequence that binds in a complementary manner to a short (∼20 nucleotide) region on the target strand (TS), known as the protospacer.

Target regions in the gene of interest are selected for mutation or excision based on the location of a short DNA region termed the protospacer adjacent motif (PAM) that is recognized by the Cas endonuclease. For *Sa*Cas9, the PAM is NNGRRT, and for *Sp*Cas9, the PAM is NGG. Both PAM sequences for Cas9 are located 3’ to the spacer binding site on the non-target strand (NTS). Both of these enzymes generate blunt double-stranded breaks (DSBs), which are typically repaired by non-homologous end-joining (NHEJ). This error-prone repair pathway directly ligates the broken ends without a homologous template, often introducing point mutations, insertions, or deletions at the break site (2).

Metagenomic sequencing and computational efforts have identified a wide variety of other CRISPR/Cas systems, and these investigations have led to the development of two members of the Cas12e family, *Deltaproteobacteria* Cas12e (*Dpb*Cas12e, CasX1) and *Planctomycetes* Cas12e (*Plm*Cas12e, CasX2) for gene editing in eukaryotes (3, 4). The PAM sequence for both Cas12e enzymes is TTCN, located 5’ to the spacer binding region on the NTS. Our previous study in which we used CasX2 and isolated DNA targets, showed enhanced cleavage when the 4^th^ nucleotide of the PAM, located at the -1 position relative to the spacer sequence, was a purine base (5).

Unlike Cas9, the CasX1 and CasX2 endonucleases generate double-stranded DNA breaks with a 5’ overhangs rather than blunt-ends (4). While the precise positioning of these staggered cuts relative to the PAM depends on the specific enzyme (CasX1 or CasX2) and the length of the gRNA spacer, cleavage on the NTS generally occurs 12–19 nucleotides downstream of the PAM, whereas cleavage on the TS is more consistently observed ∼22–25 nucleotides downstream of the PAM (3). For CasX1 cleavage of isolated DNA targets, it was previously found that shorter spacer lengths tend to shift NTS cleavage closer to the PAM, thereby producing longer overhangs (6).

Staggered 5’ overhangs generated by CasX are predicted to favor non-NHEJ DSB repair outcomes, including both homology-directed repair (HDR) and microhomology-mediated end joining (MMEJ), as shown in studies of other Cas12 proteins and engineered Cas9 variants (7–12). HDR is a high-fidelity repair pathway that uses a homologous sequence from the sister chromatid or an exogenous donor template to precisely repair DNA DSBs, but is restricted mainly to the S and G2 phases of the cell cycle (13). MMEJ is an error-prone alternative end-joining pathway that also utilizes short stretches of sequence homology (2–25 base pairs (bp)) flanking the DSB. Studies have shown that 5′ overhangs, like those produced by CasX2, significantly increase the likelihood of MMEJ promoting short-range 5’ to 3’ strand resection of the 5’ overhang and annealing of revealed micro-homologous sequences, and stabilizing the intermediate needed for re-ligation (12, 14). This typically results in deletions defined by the aligned microhomology, often removing the intervening sequence along with one of the repeats. It has been reported that MMEJ activity is low in G0 and G1, and significantly increased in dividing cells during S and G2 (15). However, assessment of the cell cycle dependence of MMEJ has largely been based on blunt DSBs, and the relative contribution of MMEJ throughout the cell cycle for the repair of staggered DSB produced by Cas12 family enzymes, including CasX, is likely increased.

In regard to PAM sequence preferences and staggered cut site patterns, we showed that CasX2 cleavage efficiency of isolated DNA varies with target site selection and spacer length, particularly when comparing gRNA spacer lengths between 17 nt to 23 nt (5). These earlier studies evaluated CasX2 activity using an isolated DNA target derived from the *CCR5* gene. In this report, we extend that analysis to determine whether similar spacer length-dependent differences in cleavage efficiency are observed when targeting *CCR5* sequences in a cellular context. Specifically, we assessed cleavage of two distinct *CCR5* targets, a transfected plasmid expressing a *CCR5* fragment, and the endogenous genomic *CCR5.* We also compared the performance of native CasX2 to a newly developed variant, CasX2^Max^ (1).

CasX2^Max^ was developed by Chen et al. using the MIDAS platform, a strategy for iterative protein engineering to improve the mammalian genome-editing efficiency of a wide range of CRISPR systems (1). MIDAS-derived enhancements were made to augment the interaction of CasX2 with DNA near the PAM on the NTS, as well as with the target DNA near the catalytic site on the TS. CasX2^Max^ incorporates three amino acid substitutions – T26R, K610R, and K808R – that collectively improved DNA cleavage. This group demonstrated that CasX2^Max^ exhibited significantly increased indel formation in mammalian cells compared to native CasX2 – establishing this strategy as a highly potent tool for genome editing (1). Building on their findings, we extended this work by performing direct comparisons of CasX2 and CasX2^Max^ cleavage of both isolated DNA and target DNA in cells, allowing us to examine not only overall cleavage activity, but also excision efficiency using paired gRNAs. This dual validation approach offers complementary insights into the improvements conferred by the CasX2^Max^ mutations.

For our studies, we assessed cleavage of the C-C chemokine receptor type 5 (*CCR5*) gene, that encodes the CCR5 co-receptor used by human immunodeficiency virus-type 1 (HIV-1) to infect target cells. A naturally occurring 32 base-pair deletion in *CCR5* (*CCR5 Δ32*) has been associated with protection against HIV-1 infection (16, 17), and has been implicated in reported cases of HIV-1 resistance and functional cures following allogeneic bone marrow transplantation with donor marrow homozygous for the *CCR5 Δ32* allele (18, 19). As a result, *CCR5* has become an important target for gene editing strategies utilizing CRISPR/Cas systems.

We first assessed the ability of native CasX2 to cleave a plasmid target expressing a region of the *CCR5* gene that was transfected in HEK293-FT cells. Using sgRNAs with varying spacer lengths, we observed spacer length-dependent differences in cleavage efficiency, consistent with our previous results using isolated DNA substrates (5). We then compared the activity of native CasX2 to CasX2^Max^, using isolated DNA as the target, and found that CasX2^Max^ exhibited significantly enhanced cleavage efficiency. To extend these findings to cellular contexts, we evaluated both nucleases for their ability to cleave *CCR5* from either a transfected plasmid or at the endogenous genomic locus (5). While both enzymes cleaved the plasmid target efficiently, only CasX2^Max^ was capable of robust genomic *CCR5* cleavage, while CasX2 had minimal activity at the endogenous *CCR5* locus. To further characterize these differences, we performed Nanopore sequencing to assess the location and frequency of cleavage events across individual sgRNAs of different spacer lengths. Assessment of plasmid targets revealed differences in cleavage patterns between 17 nt, 20 nt, and 23 nt spacer lengths for CasX2 and CasX2^Max^, and some indication of sgRNA-dependent differences in CasX2 and CasX2^Max^ repair patterns. Analysis of cleavage of the genomic target demonstrated minimal activity with the 17 nt spacer for either enzyme, but robust cleavage with gRNAs using 20 nt or 23 nt spacers for CasX2^Max^.

We performed *in silico* structural modeling of the three amino acid modifications to better define the impact of the changes in CasX2^Max^. The T26R and K610R substitutions, located in the oligonucleotide/oligosaccharide-binding fold domain (OBD) near the PAM-proximal separation of the TS and NTS DNA, promote R-loop stabilization by enhancing strand separation and increasing persistence of single-stranded DNA. Our findings show that these substitutions likely enhance protospacer unwinding and sgRNA-DNA hybrid stabilization. In contrast, the K808R substitution is positioned at the interface of the RuvC catalytic domain and the target strand loading (TSL) domain, within a basic electrostatic pocket critical for substrate engagement. This substitution improves contacts with the DNA backbone in both stage I (NTS cleavage) and stage II (TS cleavage) conformations, and is adjacent to a conserved zinc ribbon motif that undergoes redox-dependent structural transitions. These features suggest that K808R stabilizes DNA positioning near the catalytic site and helps preserve local structure under conformational strain, contributing to the enhanced editing activity of CasX2^Max^. Together, our results demonstrate that CasX2^Max^ is significantly more effective than native CasX2 for genome editing in human cells, reinforcing its potential utility for therapeutic applications.

## Materials and Methods

### CasX2 and CasX2^Max^ expression and purification

A pET23a-based construct for expressing CasX2 in *E. coli* called pET-CasX2-twst, was created using Gibson Assembly (E2621S, New England Biolabs (NEB)) to scarlessly replace the DNA sequences between the T7 promoter and T7 terminator with an *E. coli* codon-optimized *casX2* gene containing an N-terminal SV40 nuclear localization signal (NLS), and a C-terminal fusion of a nucleoplasmin NLS, 3xHA epitope tag, and TwinStrep tag. Successful products of this reaction were identified by restriction digest and Sanger sequencing. An equivalent plasmid for expressing CasX2^Max^ (defined as *Plm*CasX2 T26R K610R K808R) was then generated using Q5 Hot Start High-Fidelity DNA Polymerase (20M0493S) to perform site-directed mutagenesis to introduce the three substitutions into CasX2. Successfully mutagenized plasmids containing *casX2*^Max^ were identified by Sanger sequencing.

Chemically competent Rosetta (DE3) *E. coli* (Sigma-Aldrich, St. Louis, MO) were transformed with validated expression vectors for either CasX2 or CasX2^Max^ and used to inoculate starter cultures grown in Terrific Broth (ThermoFisher Scientific) supplemented with ampicillin (100 µg/ml) (Sigma-Aldrich) and chloramphenicol (34 µg/ml) (Sigma-Aldrich) at 37°C, with shaking at 250 RPM in 18 mm glass tubes overnight. Expression cultures were then started by diluting overnight cultures 1:100 v/v ratio in Terrific Broth (Invitrogen), and grown under identical conditions in a 4:1 flask:media v/v ratio until an OD_600_ of 0.5–0.8 was reached (3–4 hours). Cultures were then chilled on ice, and protein expression was induced by the addition of 1 mM isopropyl β-D-1-thiogalactopyranoside (IPTG) (Sigma-Aldrich), after which cells were grown for 16 hours at 16°C. Bacterial cells were then harvested by centrifugation at 4,000 *x g*, 4°C, for 10 min. The resulting cell pellet was weighed and resuspended in lysis buffer (50 mM HEPES-NaOH, 250 mM NaCl, 5 mM MgCl_2_, 1 mM TCEP, 10% glycerol, pH 8) at 5 mL per gram of cell pellet, supplemented with one cOmplete protease inhibitor tablet (Roche) per 50 mL. Resuspended samples were frozen at -80°C until purification.

For protein purification, frozen samples were thawed on ice, after which 50 U of TurboNuclease (Accelagen) per gram of cell pellet was added. Cells were then disrupted by sonication using a Model 550 Sonic Dismembrator (ThermoFisher Scientific). The lysate was clarified by centrifugation at 50,000 *x g*, 4°C, for 30 min in a LE-80K ultracentrifuge (Beckman), and the resulting supernatant was collected. During this time, 1 ml of Strep-Tactin XT 4Flow resin (IBA Life Sciences) per 25 ml of lysate was equilibrated with TwinStrep Wash Buffer (50 mM HEPES-NaOH, 500 mM NaCl, 5 mM MgCl_2_, 1 mM TCEP, 10% glycerol, pH 8). The spun supernatant was adjusted to pH 8.0 to ensure efficient affinity binding, and then mixed with the resin for 30 minutes at 4°C by gentle rocking. After incubation, the mixture was allowed to settle, and the unbound lysate was removed, leaving the resin with bound protein. The resin was washed with 20 resin volumes of TwinStrep Wash Buffer by gravity flow, after which the bound CasX proteins were eluted in two serial applications of five resin volumes of TwinStrep Elution Buffer (50 mM HEPES-NaOH, 500 mM NaCl, 5 mM MgCl2, 100 mM biotin, 1 mM TCEP, 10% glycerol, pH 8), each with 15 minutes of gentle rocking at 4°C. The presence of CasX proteins in eluates was verified via SDS-PAGE using Stain-Free protein gels (Bio-Rad Laboratories). Fractions with CasX were pooled, and dialyzed in 20k MWCO Slide-A-Lyzer dialysis cassettes (ThermoFisher Scientific) at 4°C for 16 hours with two changes of SEC buffer (25 mM sodium phosphate, 300 mM NaCl, 1 mM TCEP, 10% glycerol, pH 7.25), maintaining a minimum dialysis buffer-to-eluted sample v/v ratio of 100:1 for each buffer change. CasX proteins were then concentrated using a 50 kDa MWCO spin concentrator (ThermoFisher Scientific) at 4°C, and solubility and concentrations were verified via SDS-PAGE with Stain-Free protein gels. Samples were then further purified via size exclusion chromatography on a Superdex 200 pg column using an AKTA Go Protein Purification System (Cytiva). CasX-containing fractions were identified by SDS-PAGE, pooled and concentrated as above with 50 kDa MWCO spin concentrators. Purity was assessed by SDS-PAGE with Stain-Free gels, and found to exceed 90% (Supplementary Figure S1). Final preparations were aliquoted, snap-frozen in liquid nitrogen, and stored at -80°C.

### Guide RNA design & synthesis

Cleavage assays of isolated DNA targets utilized sgRNAv2, the native scaffold of CasX2 characterized by Tsuchida et al. (21). We designed CasX2 sgRNAs targeting *CCR5*, designated sg5 and sg10 (5), and used these gRNAs to direct cleavage of the human *CCR5* open reading frame (NC_000003.12: 46370142..46376206), as annotated in the NCBI Reference Sequence Database.

Relative to the start of *CCR5*, the 20 nt spacer version of sg5 targeted nucleotide positions 269 to 288, while the 20 nt spacer version of sg10 targeted nucleotide positions 1018 to 1037. A third gRNA, termed E6, was designed to target green fluorescent protein (*egfp*, GenBank: U55762.1) at positions 216 to 235 relative to the start of the open reading frame (22). Sequences of gRNAs are found in Supplemental Table S1 and sgRNA scaffold sequences are found in Supplemental Table S2.

cDNA templates for each CasX2 sgRNA and subsequent *in vitro* transcription were performed as described in Armstrong et al. (5). The gRNAs were heated to 70°C for 5 minutes, followed by a slow cool to 25°C at a rate of -0.1°C/second. CasX2 ribonucleoprotein (RNP) buffer (25 nM Na_3_PO_4_, 150 mM NaCl, 200mM trehalose, 1mM MgCl_2_, pH 7.5) was then added to each sample, and the mixture heated to 50°C for 5 minutes and then cooled to 25°C at a rate of -0.1°C/second. Folded gRNAs were then quantitated using a Qubit BR RNA Kit (ThermoFisher Scientific), diluted to 10 µM, and stored in aliquots at -80°C.

### Cleavage assays of isolated DNA

Ribonucleoproteins (RNPs) of gRNA plus CasX2 or CasX2^Max^ were generated by combining a 1:1.2 ratio of CasX2 or CasX2^Max^ and gRNA in CasX2 RNP buffer. The mixture was incubated at 37°C for 10 minutes and then placed on ice. RNP complexes were stably stored at -20°C for up to one week. Cleavage reactions were prepared by combining RNPs with target DNA in CasX2 reaction buffer (20 mM Tris HCl pH 7.5, 150 mM NaCl, 10 mM MgCl_2_, 1 mM TCEP, 5% glycerol) in a final volume of 20 µL. The reactions were incubated at 37°C for 1 hour, and stopped by heating to 65°C for five minutes, followed by an incubation with 2 µL of Proteinase K at 50°C for 30 minutes. Cleavage products were visualized by electrophoresis on agarose Tris-Borate-EDTA (TBE) gels stained with GelRed (Sigma-Aldrich), and quantified by densitometric analysis using Image Lab 6.1 (Bio-Rad Laboratories), and statistically analyzed using Prism 10.4.0 (Dotmatics).

### Structural prediction of CasX2^Max^

The structures of CasX2^Max^ bound to a gRNA and its respective DNA target were predicted for confirmational state I (NTS loading), state II (TS loading), and state III (preloading inactive) using the protein structure homology-modeling server SWISS-MODEL (23). The cryo-EM structures of enzymatically inactive d*Plm*CasX complexed with target DNA and sgRNAv1 in various states (21) served as the templates: state I (PDB ID: 7WAY), state II (PDB ID: 7WAZ) and state III (PDB ID: 7WB0). These CasX2 structures are in complex with a target DNA and sgRNAv1, that consists of the native CRISPR RNA (crRNA) and trans-activating CRISPR RNA (tracrRNA) elements of CasX1 (Supplemental Figure S2). This was chosen over the CasX2 sgRNA (sgRNAv2), as a full set of cryo-EM solutions of sgRNAv2 with CasX2 were unavailable (state I: 7WB1). However, sgRNAv1 and sgRNAv2 share 88% identity and provide cross-functionality for each other’s cognate CasX cleavage, supporting the use of this gRNA in our modeling. The sgRNA scaffold sequences are found in Supplemental Table S2.

The five SWISS-MODEL predictions with the highest Global Model Quality Estimate (GMQE) scores for each state were selected for further analysis. These models were aligned to their respective state’s cryo-EM structure using the MatchMaker function in ChimeraX (version 1.9), developed by the Resource for Biocomputing, Visualization, and Informatics at the University of California, San Francisco (24). Structural similarity was quantified by calculating the root-mean-square deviation (RMSD) across the full-length protein (960 atom pairs). The CasX2^Max^ model with the lowest RMSD in each state was then substituted for d*Plm*CasX, while retaining the corresponding sgRNA and dsDNA target from the original equivalent CasX2:sgRNAv1 cryo-EM structure.

### Plasmids to express CasX2, CasX2^Max^ and sgRNAs

Plasmid vectors for CasX2, *Sa*Cas9, and CasX2^Max^ expression with single and dual U6 promoters/sgRNA cassettes were created through standard cloning methods and sequence-verified by Nanopore sequencing using Circuit-seq (25). Plasmid names and sequences are found in Supplemental Table S3. Briefly, the mammalian codon-optimized CasX2 was from pBLO 62.5, a gift from Jennifer Doudna and Benjamin Oakes (Addgene plasmid # 123124) (26). *Sa*Cas9, and portions of the sgRNA cassette, were derived from pX601-AAV-CMV::NLS-SaCas9-NLS-3xHA-bGHpA;U6::BsaI-sgRNA (Addgene plasmid # 61591), a gift from Feng Zhang (27); pDG459 (Addgene plasmid # 100901), a gift from Paul Thomas (28); and pGGAselect (Addgene plasmid # 195714), a gift from Eric Cantor (29). Oligonucleotide sequences for cloning of gRNAs are found in Supplemental Table S4.

### Transfection of HEK293-FT cells with gRNA plus CRISPR/CasX2 or CRISPR/CasX2^Max^

HEK293-FT cells (CVCL_6911, ThermoFisher Scientific) were maintained in Dulbecco’s Modified Eagle’s Media (DMEM) with 10% fetal bovine serum (FBS). Transfection of HEK293-FT cells with plasmids expressing both CasX2 and gRNA were performed using Lipofectamine 3000 (ThermoFisher Scientific) according to the manufacturer’s directions. Twenty-four hours prior to transfection, HEK293-FT cells were seeded at 1.5 x 10^5^ cells/well in 1 mL of complete growth medium in 12 well tissue culture plates. One µg of plasmid DNA was diluted in 50 µl of serum-free Opti-MEM-reduced serum media. Cells were transfected with either 50 ng of pcDNA3-CCR5 target and 950 ng of the indicated enzyme/guide plasmid, or 1,000 ng of the enzyme/guide plasmid. Transfected cells were incubated at 37°C in a CO_2_ incubator for 18–24 hours post-transfection before replacing media with fresh complete growth media. After 72 hours, the genomic DNA was isolated using the Qiagen Blood and Tissue kit (Qiagen) or Monarch genomic DNA purification kit (NEB) according to the manufacturers’ instructions. Genomic DNA was analyzed on Nanodrop 2000 (ThermoFisher Scientific) and Qubit4 Fluorometer using the Qubit double-stranded DNA BR assay kit (ThermoFisher Scientific).

### High-fidelity DNA polymerase chain reaction (PCR) and T7 endonuclease assay

Genomic DNA isolated from CRISPR-treated cells was used as template for a Q5 Polymerase PCR reaction (NEB). Each set of primers were analyzed for optimal PCR conditions (see Supplemental Table S5 for primer sequences and Supplemental Table S6 for PCR conditions). Prior to running the T7 endonuclease assay, 10 µl of each PCR reaction was examined on an agarose/Tris-Borate-EDTA (TBE) gel to confirm correct amplification of the expected DNA product. The remaining 40 µl of the PCR reaction was purified using the Monarch PCR & DNA cleanup kit (NEB) according to the manufacturer’s instructions.

To test for cleavage, 442 ng of clean PCR product was mixed with NEBuffer 2 (50 mM NaCl, 10 mM Tris-HCl, 10 mM MgCl_2_, pH7.9) (NEB) in a final volume of 42 µl (10.5 ng/µl). This was divided into two PCR tubes (21 µl/each), and one replicate received 1 µl (10 units) of T7 endonuclease (M0302, NEB). All replicates were briefly spun, and incubated at 95° C for 5 minutes to denature the DNA, followed by annealing at 95° C – 85° C at a ramp rate of -2° C/second, followed by further cooling from 85° C-25° C at a ramp rate -0.1° C/second. Samples were mixed, spun briefly, and incubated at 37° C for 15 minutes. Samples (+/- T7 endonuclease) were analyzed on a 1.0% agarose/TBE gel and visualized using SyBR safe DNA gel stain (Invitrogen).

### Nanopore sequencing

Briefly, each PCR product was end-repaired with an Ultra II End-prep (E7647S, NEB), and cleaned up with AMPure beads (Beckman Coulter). Each end-repaired and cleaned up PCR product was barcoded with the provided barcodes using Blunt/TA ligase (M0367S, NEB). The barcoded PCR products were then pooled, cleaned up a second time with AMPure beads, and then adapter-ligated using the Native Adapter and NEB Next Quick Ligation Module (E6056S, NEB) and SQK NBD114.96 (Oxford Nanopore Technologies). Sequencing was performed using a MinION Flow Cell (R10.4.1, Oxford Nanopore Technologies) and MinKNOW with parameters: Kit = SQK NBD114.96; min base pair length = 200 bp; Output = Base calling Raw Reads/POD5; Barcoding = FASTQ; and alignment = off. Super-accuracy base calling using Dorado (Oxford Nanopore Technology) was performed, and FastQ files were analyzed for editing efficiency using CRISPResso2 (30) as previously described (31).

## Results

### Targeting *CCR5* with CasX2

To evaluate the ability of CasX2 to disrupt the human *CCR5* gene in cells, we selected two CasX2 gRNAs, sg5 and sg10, that we had previously shown to be effective in cleaving isolated DNA using recombinant CasX2 (5). These target sequences overlap with two *Sa*Cas9 gRNAs described in a recent study (32), allowing for direct comparison with an established Cas9-based approach (Figure 1A). Given that recombinant CasX2 exhibits variable cleavage of isolated DNA depending on spacer length (5), we assessed its activity in cells across multiple spacer lengths. HEK293-FT cells were transfected with plasmids encoding CasX2 and either sg5 or sg10 using spacer lengths of 17, 20 or 23 nt, along with a target plasmid containing the *CCR5* gene. Three days post-transfection, total cellular DNA was extracted, and the target region for sg5 or sg10 was amplified by high-fidelity PCR and subjected to T7 endonuclease assay to detect indels (Figure 1B). Cleavage by sg5 was most effective with a 23 nt spacer, and sg10 showed maximal activity with a 20 nt spacer. Both sg5 and sg10 had minimal activity with a 17nt spacer.

**Figure 1.**
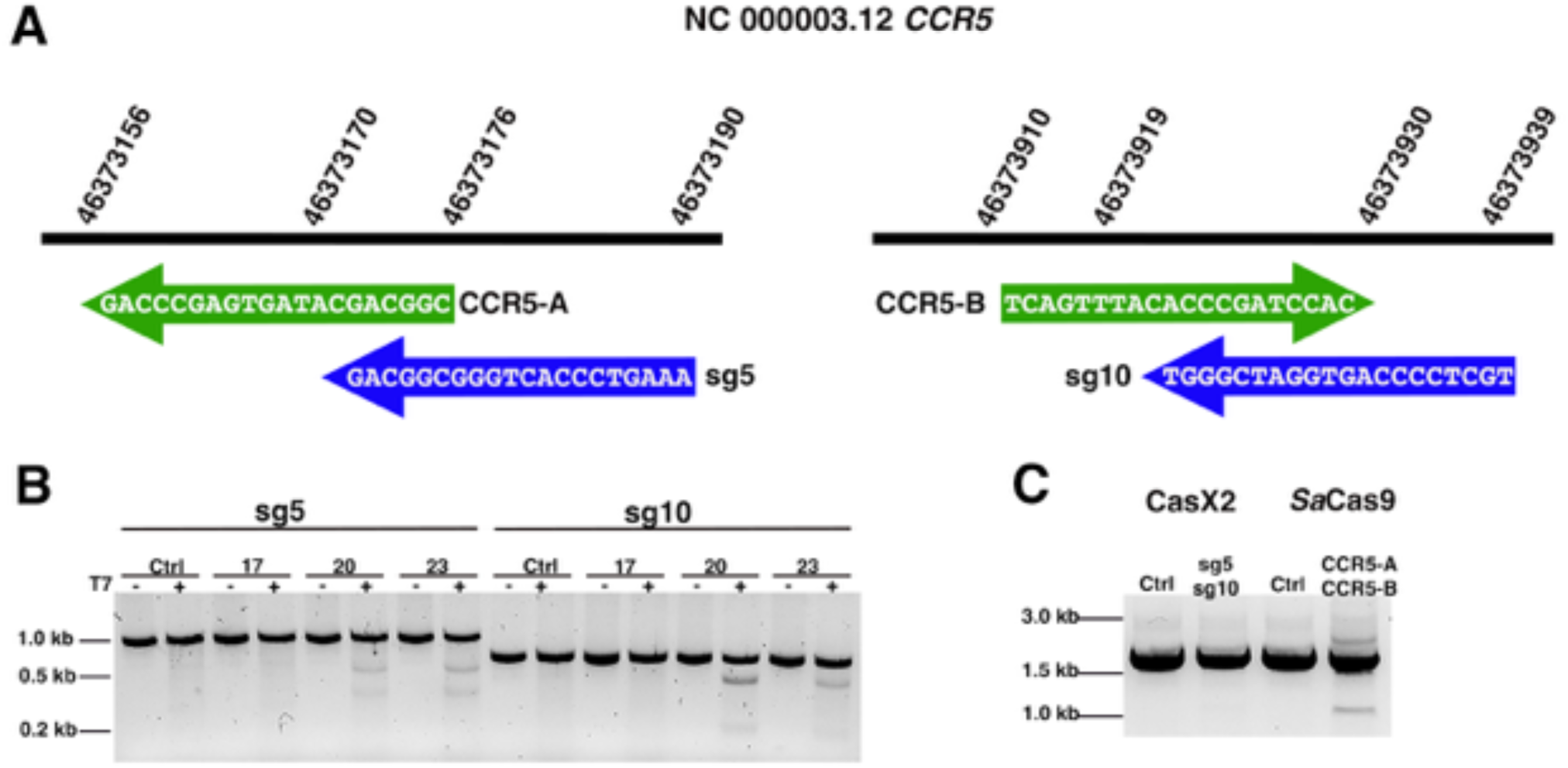
Schematic of CasX2 and *Sa*Cas9 *CCR5* gRNAs and their editing abilities. **A**. Location of tested gRNAs on chromosome 3. *Sa*Cas9 gRNA spacer binding sites CCR5-A and CCR5-B are shown with green arrows, and CasX2 gRNA spacer binding sites sg5 and sg10 sites are indicated with blue arrows. **B.** Gel electrophoresis was used to analyze a T7 endonuclease assay assessing the activity of CasX2 gRNAs sg5 and sg10 of varying guide lengths (17 nt, 20 nt or 23 nt) in HEK293-FT cells. **C.** PCR to detect excision of genomic *CCR5* following transfection of HEK293-FT cells with either CasX2 and sg5, CasX2 and sg10, *Sa*Cas9 and CCR5-A, *Sa*Cas9 and CCR5-B, or respective controls of Cas enzyme but no gRNA.

We next tested the ability of sg5 and sg10 to disrupt the endogenous genomic *CCR5* gene when co-administered as a gRNA pair in HEK293-FT cells. In contrast to *Sa*Cas9 CCR5-A and CCR5-B gRNAs described by Dash, et al. (32), CasX2 yielded no detectable *CCR5* excision as assessed by endpoint PCR (Figure 1C). These findings indicated that CasX2 has poor ability to disrupt *CCR5* compared to *Sa*Cas9, and prompted us to test the cleavage efficiency of various *CCR5* targets with CasX2^Max^.

### CasX2^Max^ exhibits enhanced cleavage of an isolated *CCR5* DNA target compared to CasX2

We established an optimized protocol using TwinStrep-affinity purification to isolate CasX2^Max^ and CasX2, achieving comparable purity and yield for both enzymes (Supplemental Figure S1). In addition, we purified *in vitro*-transcribed *CCR5* gRNAs sg5 and sg10 with 20 nt spacers (as shown in Figure 1) (5), and prepared equimolar quantities of RNPs containing CasX2 or CasX2^Max^. Using these RNPs, we compared cleavage of the isolated *CCR5* DNA target with either CasX2^Max^ or CasX2 (Figure 2A). Densitometric analysis revealed that CasX2^Max^ displayed an average of 2.7-fold increase in cleavage efficiency compared to CasX2 (Figure 2B). Importantly, no non-specific cleavage of the *CCR5* DNA target was observed when the gRNA E6 targeting enhanced GFP (EGFP) was used (Figure 2A).

**Figure 2.**
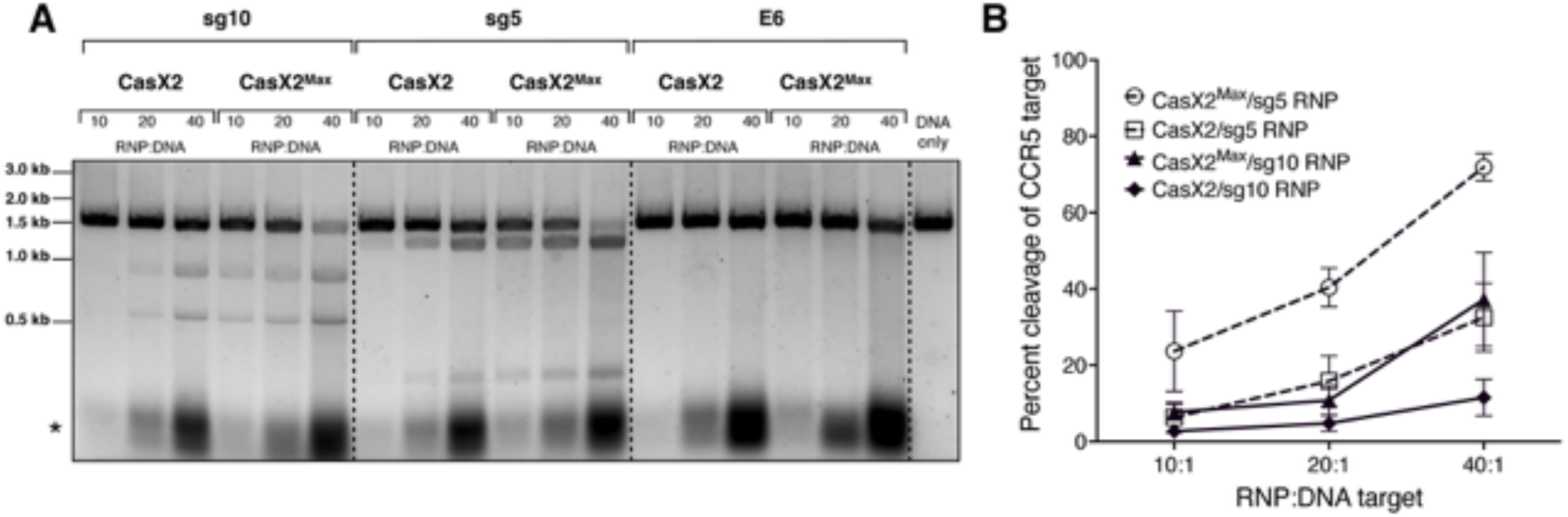
CasX2^Max^ demonstrates increased cleavage activity of isolated DNA targets compared to CasX2. **A.** RNPs were generated by incubating CasX2 or CasX2^Max^ with purified sgRNAs. RNPs were then incubated with target *CCR5* DNA at 37°C for 60 minutes, and DNA cleavage assessed by gel electrophoresis. The “*” denotes the unbound sgRNA. **B.** Densiometric analysis of gel images of three independent cleavage assays were processed using Image Lab, and statistics calculated using Prism.

### Increased targeting of genomic *CCR5* by CasX2^Max^ compared to CasX2

To assess the activity of CasX2^Max^ in human cells, we generated plasmid vectors that expressed human codon-optimized CasX2^Max^ and gRNAs sg5 or sg10 with 17, 20, or 23 nt spacer lengths. HEK293-FT cells were transfected with either plasmids expressing CasX2 or CasX2^Max^ plus sg5 or sg10, and a target plasmid that included the relevant regions of *CCR5*. Editing was assessed by high-fidelity PCR followed by a T7 endonuclease assay and agarose gel electrophoresis (Figure 3A). Consistent with our previous results with isolated DNA and recombinant CasX2, we observed increased editing activity by CasX2^Max^ compared to CasX2 with both sg5 and sg10 at all three spacer lengths. Densitometric analysis demonstrated an approximately 4-fold increase in indel formation with the 20 and 23 nt spacer lengths for both sg5 and sg10 (Figure 3B). CasX2^Max^, in contrast to CasX2, exhibited detectable indel formation for the 17nt spacer length for both sgRNAs, although the activity was still less than the 20 and 23nt spacers (Figure 3A and 3B).

**Figure 3.**
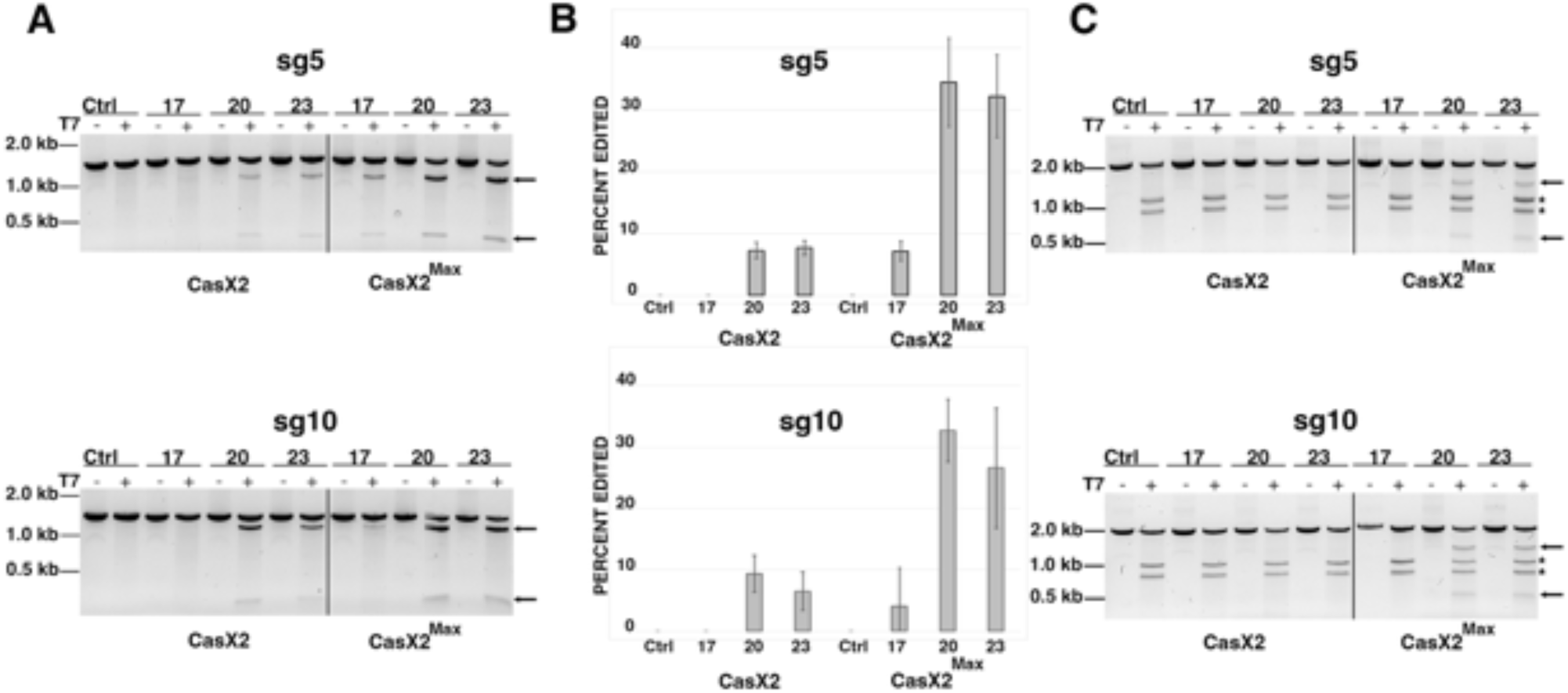
Improved targeting of *CCR5* by CasX2^Max^ compared to CasX2. **A.** T7 endonuclease assay of CasX2 and CasX2^Max^ editing of a *CCR5* plasmid target in HEK293-FT cells using sg5 and sg10 gRNAs with varying spacer lengths (17 nt, 20 nt, and 23 nt). **B.** Quantification of plasmid editing by densitometry. Mean editing is based on n=3 replicates, with error bars indicating standard deviation. **C.** T7 endonuclease assay of CasX2 and CasX2^Max^ editing of genomic *CCR5* in HEK293-FT cells by sg5 and sg10 gRNAs of varying spacer lengths (17 nt, 20 nt, and 23 nt). Arrows indicate expected edited bands. Asterisks “*” indicate a product resulting from T7 endonuclease targeting of the heteroduplexes resulting from the presence of the single *CCR5 Δ32* allele in HEK293-FT cells.

As CasX2 showed little detectable editing of the native genomic *CCR5* locus, we next tested targeting by CasX2^Max^ with either sg5 or sg10 at spacer lengths of 17, 20 and 23 nt (Figure 3C). HEK293 cells are heterozygous for the *CCR5 Δ32* allele, which complicates analysis by T7 assays as the resulting heteroduplex produces T7 cleavage products in all samples (noted with “*” in Figure 3C) (33). We did not observe discernable editing using sgRNAs with the 17 nt spacer length with either CasX2 or CasX2^Max^ as assessed by the T7 endonuclease assay, suggesting that, in contrast to cleavage of isolated DNA (5, 6), gRNAs with spacers 17 nt or shorter are poorly functional in cells. However, unlike CasX2, we observed significant cleavage activity of genomic *CCR5* with CasX2^Max^ using sg5 and sg10 gRNAs at both 20 nt and 23 nt spacer lengths (Figure 3C).

### Detectable differences in CasX2 and CasX2^Max^ indels at some target sites in cells

It was previously shown that the site of CasX1 cleavage on isolated DNA target may partially depend on the length of the gRNA spacer (6). Although spacer length impacts CasX2 function of isolated DNA in a gRNA-specific manner (5), the impact on the nature of the cleavage event is unclear for CasX2. To further characterize the editing events mediated by CasX2 and CasX2^Max^, we performed long-read Nanopore sequencing at sites in *CCR5* targeted by CasX2 and CasX2^Max^. Total DNA from HEK293-FT cells transfected with the *CCR5* plasmid target, along with CasX2 and CasX2^Max^ plus sg5, was used to generate high-fidelity PCR products that were subjected to Nanopore V14 sequencing and super-accuracy base-calling. Reads were assessed for the presence of deletions using CRISPResso2 (30) as previously described for Oxford Nanopore Technology reads (31). These analyses recapitulated the superior performance of CasX2^Max^ compared to CasX2 (Figure 4).

**Figure 4.**
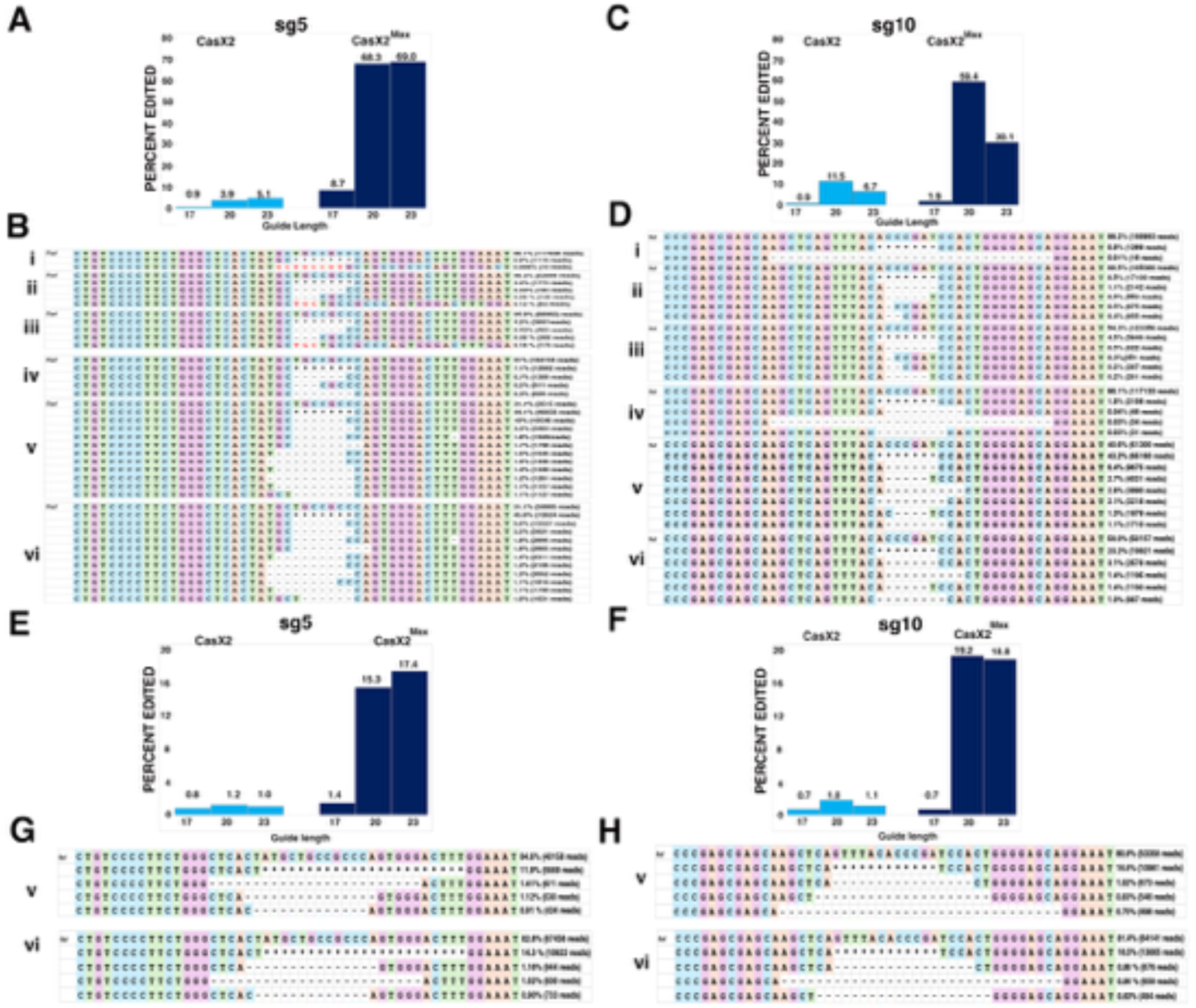
Improved targeting of *CCR5* with CasX2^Max^. **A.** Editing of a transfected *CCR5* target plasmid following co-transfection of HEK293-FT cells with plasmids encoding CasX2 or CasX2^Max^ and sg5 with the indicated spacer length. Editing was assessed by Nanopore sequencing and editing efficiency was assessed using CRISPResso2. **B.** The most frequently detected edits are shown for CasX2 (i-iii) and CasX2^Max^ (iv-vi), and for spacer lengths of 17 nt (i, iv), 20 nt (ii, v), and 23 nt (iii, vi). **C.** Editing efficiency of genomic *CCR5* following transfection of HEK293-FT cells with plasmids encoding CasX2 or CasX2^Max^ and sg10 with the indicated spacer length. Editing was assessed by Nanopore sequencing and editing efficiency was assessed using CRISPResso2. **D.** The most frequently detected edits are shown for CasX2 (i-iii) and CasX2^Max^ (iv-vi), and for spacer lengths of 17 nt (i, iv), 20 nt (ii, v) and 23 nt (iii, vi). **E.** Editing of genomic *CCR5* following transfection of HEK293-FT cells with plasmids encoding CasX2 or CasX2^Max^ and sg5 with the indicated spacer length. Editing was assessed by Nanopore sequencing and editing efficiency by CRISPResso2. **F.** Editing of genomic *CCR5* following transfection of HEK293-FT cells with plasmids encoding CasX2 or CasX2^Max^ and sg10 with the indicated spacer lengths. Editing was assessed by Nanopore sequencing and editing efficiency by CRISPResso2 analysis. **G.** The most frequently detected edits are shown for CasX2^Max^ plus sg5 of spacer lengths of 20 nt (v) and 23 nt (vi). **H.** The most frequently detected edits are shown for CasX2^Max^ plus sg10 of spacer lengths of 20 nt (v) and 23 nt (vi). The “*” indicates summation of reads with edits in the region shown but individually comprising less than 1% of the edited reads. “Ref” indicates the reference sequence.

Consistent with our T7 endonuclease assays (Figure 3A), we detected more targeted indels in cells treated with CasX2^Max^/sg5 compared with CasX2/sg5 at all spacer lengths, as quantified using CRISPResso2 analysis of the Nanopore sequencing data (Figure 4A). We observed differences in the most common indel detected by Nanopore sequencing (Figure 4B), with insertions amongst the most common mutations detected at the sg5 target site with CasX2, but not with CasX2^Max^, across all spacer lengths (Figure 4B, i-iii vs iv-vi). These results suggest that while the amino acid modifications in CasX2^Max^ significantly increases editing frequency, they also likely alter the type of indels generated in cells. For sg10 (Figure 4C), we also detected increased targeting across spacer lengths when comparing CasX2^Max^ to CasX2. For the 17 nt spacer sgRNA, we detected larger 31 bp deletions in both the CasX2 and CasX2^Max^ samples that were not seen with sg10 at the 20 and 23 nt lengths (Figure 4D, i and iv, compared with ii, iii and v, vi). Analyses of cleavage of an abundant plasmid target revealed modest changes in the nature of the indels produced, independent of the target site, spacer length, or CasX2 enzyme used. We next tested targeting of genomic CCR5 in HEK293-FT cells using CasX2 and CasX2^Max^, and sg5 and sg10. As observed by T7 assay (Figure 3C), few indels were detected using CasX2 with any length spacer or CasX2^Max^ with the 17 nt spacer length (Figure 4E and 4F). We observed slightly increased targeting with the 23 nt sg5 and the 20 nt sg10. A similar distribution of deletions was observed for the 20 nt and 23 nt spacers for both sg5 and sg10 with CasX2^Max^ (Figure 4G and 4H). For both sgRNAs and at both 20 nt and 23 nt spacer, we observed a trend towards larger deletions in the genomic CCR5 target (Figure 4G and H) than when targeting the transfected plasmid target (Figure 4B and 4C). The most commonly identified deletions for both sg5 and sg10 when using CasX2^Max^ to target genomic CCR5 are indicative of MMEJ. Potential microhomologies identified for the top deletions include TGGG and CA for sg5 (Figure 4G), and CA, CT, and GAGCA for sg10 (Figure 4H).

### CasX2^Max^ is equivalent to *Sa*Cas9 in inducing disruption of *CCR5* in cells

We initially tested the cleavage efficiency of CasX2^Max^ based on the poor ability of the native CasX2 to excise *CCR5* genomic DNA. Therefore, we next assessed the targeting of genomic *CCR5* following transfection of CasX2 or CasX2^Max^ plus gRNAs in HEK293-FT cells. We first compared indel induction by either CasX2^Max^ with either sg5 and sg10, or *Sa*Cas9 with previously published sgRNAs CCR5-A and CCR5-B (32). Nanopore sequencing and CRISPResso2 analysis demonstrated significant differences in the targeting efficiency between CCR5-A and CCR5-B, and relatively comparable efficiency between CCR5-A and CasX2^Max^ sg5 and sg10 (Figure 5A). Analysis of the targeting event by CRISPResso2 indicates the predominant indels result from MMEJ for CasX2^Max^, and from a mix of NHEJ and MMEJ for *Sa*Cas9 (Figure 5B-5E).

**Figure 5.**
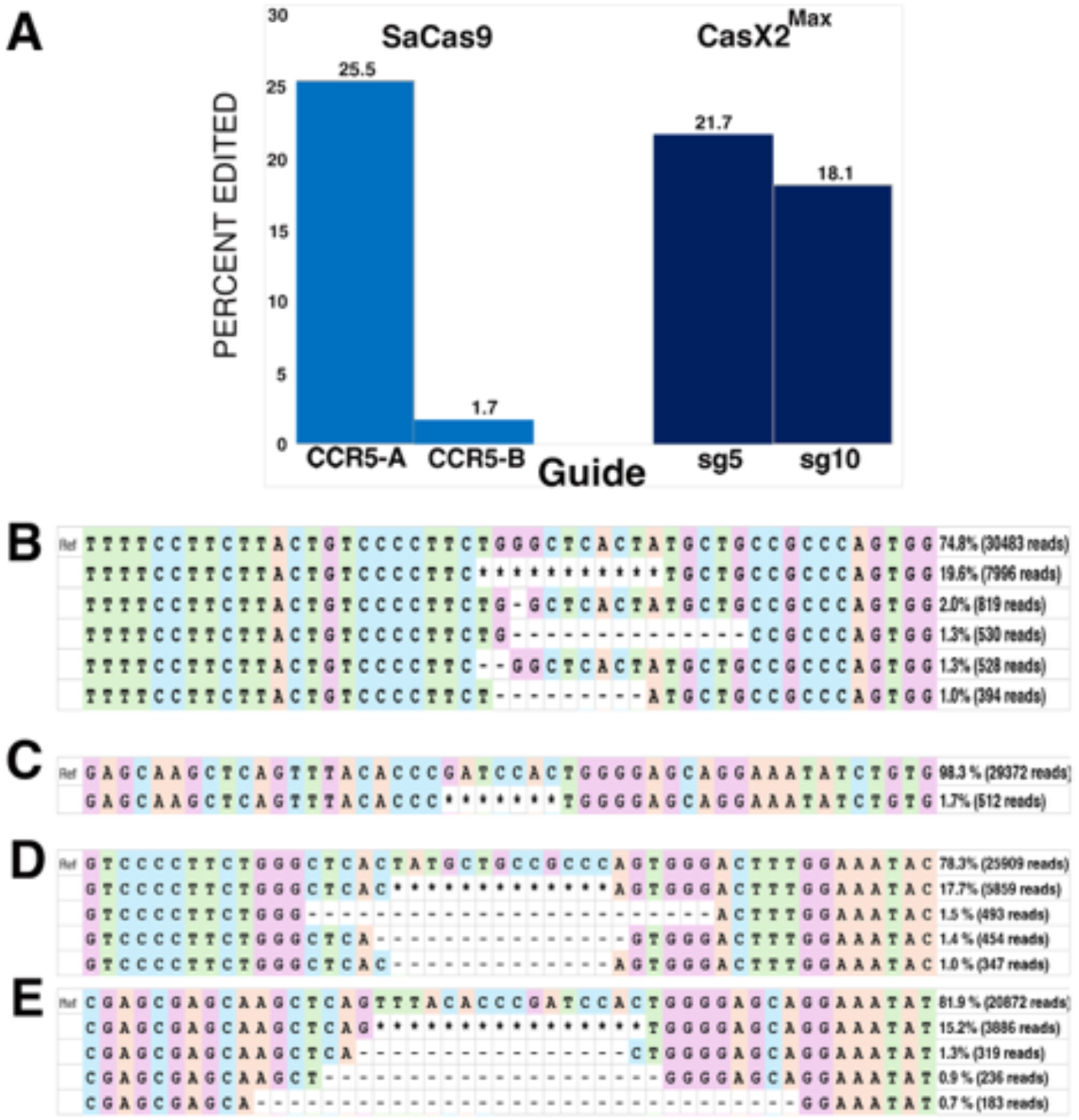
Comparable targeting of genomic *CCR5* with CasX2^Max^ and *Sa*Cas9. **A.** Editing of genomic *CCR5* following transfection of HEK293-FT cells with plasmids encoding *Sa*Cas9 and sgRNAs CCR5-A or CCR5-B, or CasX2^Max^ and sgRNAs sg5 or sg10, was assessed by Nanopore sequencing and CRISPResso2 analysis. **B.** The most frequently detected edits are shown for *Sa*Cas9 CCR5-A at the 20 nt spacer length. **C.** The most frequently detected edits are shown for *Sa*Cas9 CCR5-B at the 20 nt spacer length. **D.** The most frequently detected edits are shown for CasX2^Max^ with sg5 at the 23 nt spacer length. **E.** The most frequently detected edits are shown for CasX2^Max^ with sg10 at the 23 nt spacer length. The “*” indicates summation of reads with edits in the region shown but individually comprising less than 1% of the edited reads.

### CasX2^Max^ mediates disruption of the *CCR5* gene in a human cell line

Based on the enhanced activity of CasX2^Max^ relative to CasX2, we next assessed the ability of the combined use of both sg5 and sg10, along with CasX2^Max^, to excise the intervening region of genomic DNA. We assessed excision of *CCR5* using these paired gRNAs and either CasX2 or CasX2^Max^ by end-point PCR and Nanopore long-read sequencing (Figure 6). In contrast to CasX2, we readily observed the appearance of a PCR product consistent with excision of the genomic DNA between the sg5 and sg10 target sites when using CasX2Max (Figure 6A). Nanopore sequencing of a PCR product across the region of *CCR5* containing both the sg5 and sg10 target sites, confirmed that the HEK293-FT cells are heterozygous for the *CCR5 Δ32* allele (Figure 6B, “*Δ*32”) as previously reported for parental HEK293-T cells (CVCL_0063) (33). Alignment of the Nanopore sequencing reads with the reference *CCR5* genomic DNA sequence using Minimap2 revealed excision of the intervening region of DNA between the sg5 and sg10 target sites when using CasX2^Max^ but not CasX2 (Figure 6B). Thus, CasX2^Max^ can be used to excise a large region of the endogenous *CCR5* gene in human cell lines.

**Figure 6.**
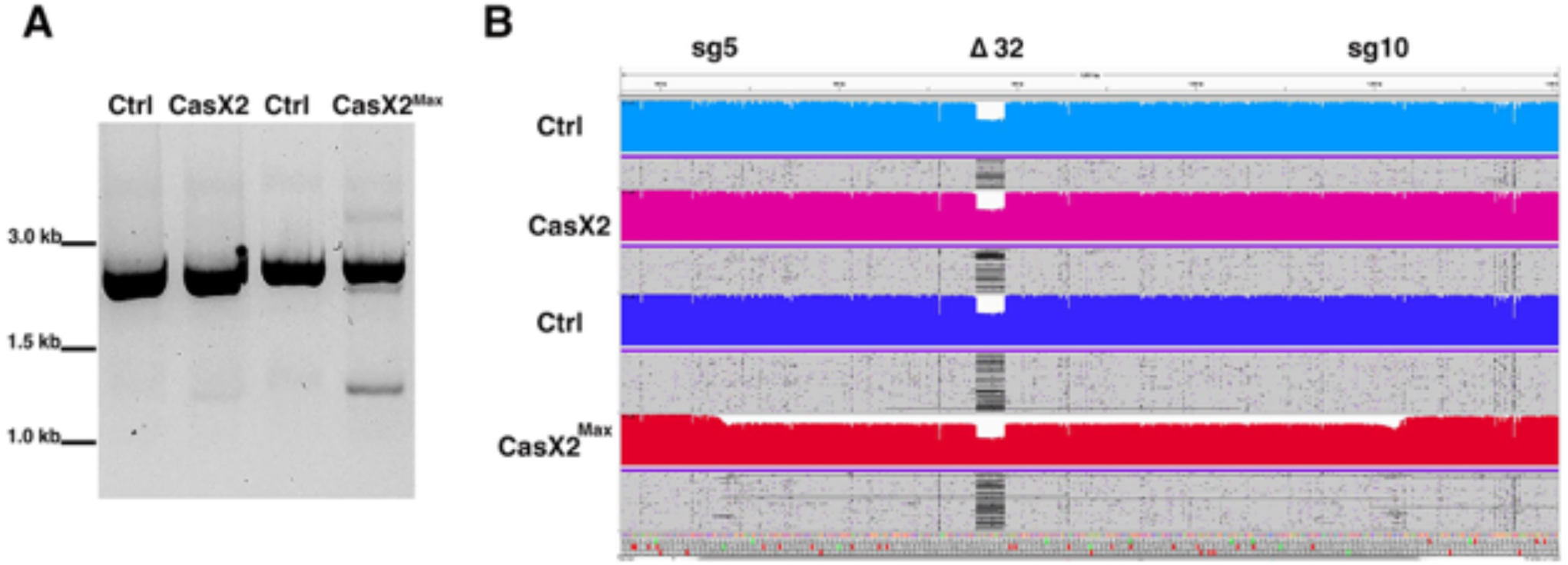
Disruption of *CCR5* using CasX2^Max^ and paired sg5 and sg10 sgRNAs. **A.** PCR amplification of genomic *CCR5* after transfection of HEK293-FT cells with scaffold only-plasmids (Ctrl), or with a combination of both sg5 and sg10 with either CasX2 or CasX2^Max^. **B.** Minimap2 alignments of Nanopore sequencing data obtained from high-fidelity PCR were displayed in Integrative Genomics Viewer (IGV), demonstrating excision of genomic *CCR5* between the sg5 and sg10 binding sites. The target sites for sg5 and sg10 are indicated, as is the location of the *CCR5* D32 allele.

### Structural basis for enhanced PAM-proximal DNA engagement by T26R and K610R in CasX2^Max^

To better understand how each MIDAS-derived amino acid substitution contributes to the enhanced cleavage activity of CasX2^Max^, we generated structural models of the protein in the three conformational states defined by Tsuchida et al. (21) using SWISS-MODEL: NTS cleavage (state I, PDB: 7WAY), TS cleavage (state II, PDB: 7WAZ), and the inactive pre-cleavage conformation (state III, PDB: 7WB0). The structural models for states I and II produced QMEANDisCo Global scores of 0.87 and 0.82, respectively, indicating high-confidence predictions (34). The model for state III received a lower score of 0.70, consistent with the structural disorder in the Helical-II (H2) domain observed in the original cryo-EM structure. The target DNA and spacer (Supplemental Figure S2) from each cryo-EM structure were then superimposed into the appropriate modeled state of CasX2^Max^.

The T26R and K610R MIDAS substitutions, both located on the OB-fold domain, were introduced based on their proximity to the PAM on the NTS, where they were hypothesized to enhance R-loop formation by stabilizing strand separation (1). Similar arginine substitutions near the PAM in other CRISPR enzymes have been shown to strengthen PAM-proximal binding, increase target DNA residence time, and pre-organize the nuclease domains for catalysis (35, 36). In evaluating these modifications in CasX2^Max^, we based our structural analyses on state I, as neither of these residues showed appreciable shifts between states I and II for either CasX2 or CasX2^Max^ (data not shown). In wild-type CasX2, the uncharged hydroxyl group of T26 is directed towards the fork where the NTS and TS separate, and the TS shifts its base pairing to the sgRNA spacer sequence. However, T26 imposes only a minimal steric hinderance in this space, and is too distant to interact electrostatically with either DNA strand (Figure 7A). Instead, T26 contributes structurally to a rigid loop (G24–K25–T26–G27–P28) that orients positively charged K25 towards the NTS backbone. In CasX2^Max^, substitution with arginine places a bulkier, positively charged guanidinium group closer to the fork of the NTS and TS DNA (Figure 7B). This is likely to enhance the existing outward rotation of TS bases seeking to reduce electrostatic interactions with the R26 side chain. This movement will, in turn, favor alignment of these bases to their complementary counterparts on the sgRNA spacer, increasing R-loop stability.

**Figure 7.**
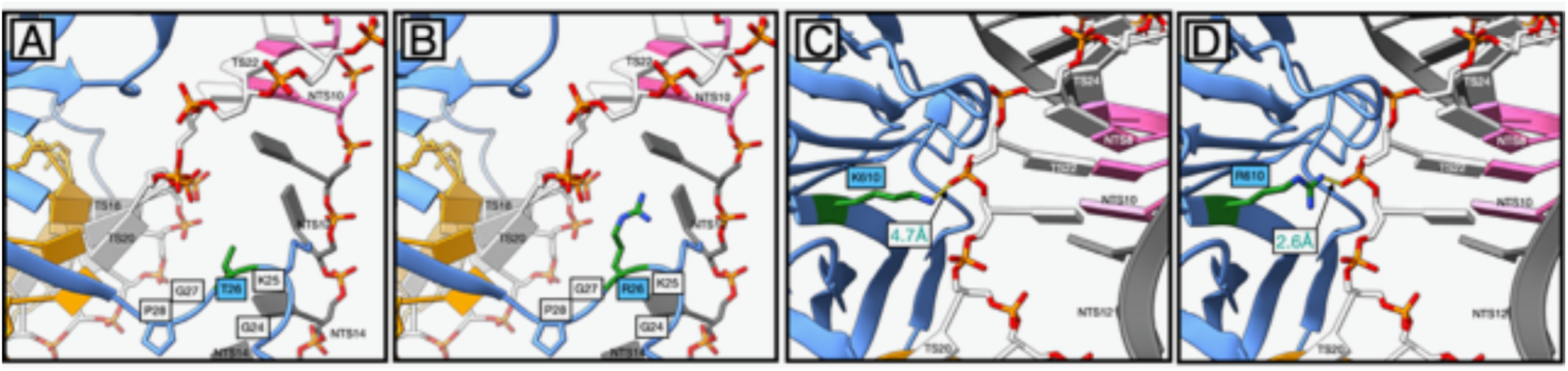
Structural modeling reveals enhanced PAM-proximal DNA interactions mediated by T26R and K610R substitutions in CasX2^Max^. **A**. Positioning of T26 in CasX2. **(B).** Positioning of R26 in CasX2^Max^. Panels **A** and **B** show (in green) the site where double-stranded DNA (dsDNA) separates into single strands. **C.** Positioning of K610 in CasX2. **D.** Positioning of R610 in CasX2^Max^. Panels **C** and **D** show the location near the DNA fork adjacent to the PAM duplex. The target strand (TS, white) and non-target strand (NTS, dark grey) are shown with the NTS PAM nucleotides (TTCA) highlighted in pink. Gold nucleotides represent the spacer region of the sgRNA, complementary to the TS protospacer. Individual nucleotide positions are labeled according to their strand designation (e.g., TS12, NTS20).

The K610R substitution of CasX2^Max^ alters how the enzyme interacts with a portion of the TS backbone immediately opposite the PAM on the NTS (Figure 7C). In wild-type CasX2, the lysine side chain at position 610 is 4.7 Å from the phosphodiester backbone of TS22, a sub-optimal distance for electrostatic interactions (Figure 7C). However, in CasX2^Max^, the K610R substitution allows the guanidinium group of the arginine to be sited closer to this backbone and enables the formation of a hydrogen bond (2.6 Å) with TS22 (Figure 7D). This interaction likely enhances local DNA binding stability at the PAM-proximal region, contributing to improved target sequence engagement, and increased cleavage activity observed with CasX2^Max^.

### K808R enhances DNA strand stabilization across CasX2 catalytic states

The K808R substitution in CasX2^Max^ resides within the RuvC domain, near its interface with the TSL domain, and is situated 10-15 Å the catalytic triad (D659, E756, D922) (3, 21). In state I (NTS loading), K808 in native CasX2 is 3.8 Å from the NTS phosphate backbone between bases 20 and 21, a distance that is sub-optimal for hydrogen bond formation (Figure 8A). K808 also participates in a cluster of basic and polar residues from both the RuvC (R764, R768) and TSL (R856, K898, R904, K955) domains that help align the NTS phosphate backbone for RuvC engagement (Figure 8B) (37). Substitution with arginine in CasX2^Max^ enhances local contacts with the NTS backbone, both by forming an additional hydrogen bond (2.0 Å and 2.2 Å) (Figure 8C) and by contributing to a denser cluster of basic residues surrounding the RuvC cleft (Figure 8D). Together, these interactions improve NTS positioning near the catalytic core.

**Figure 8.**
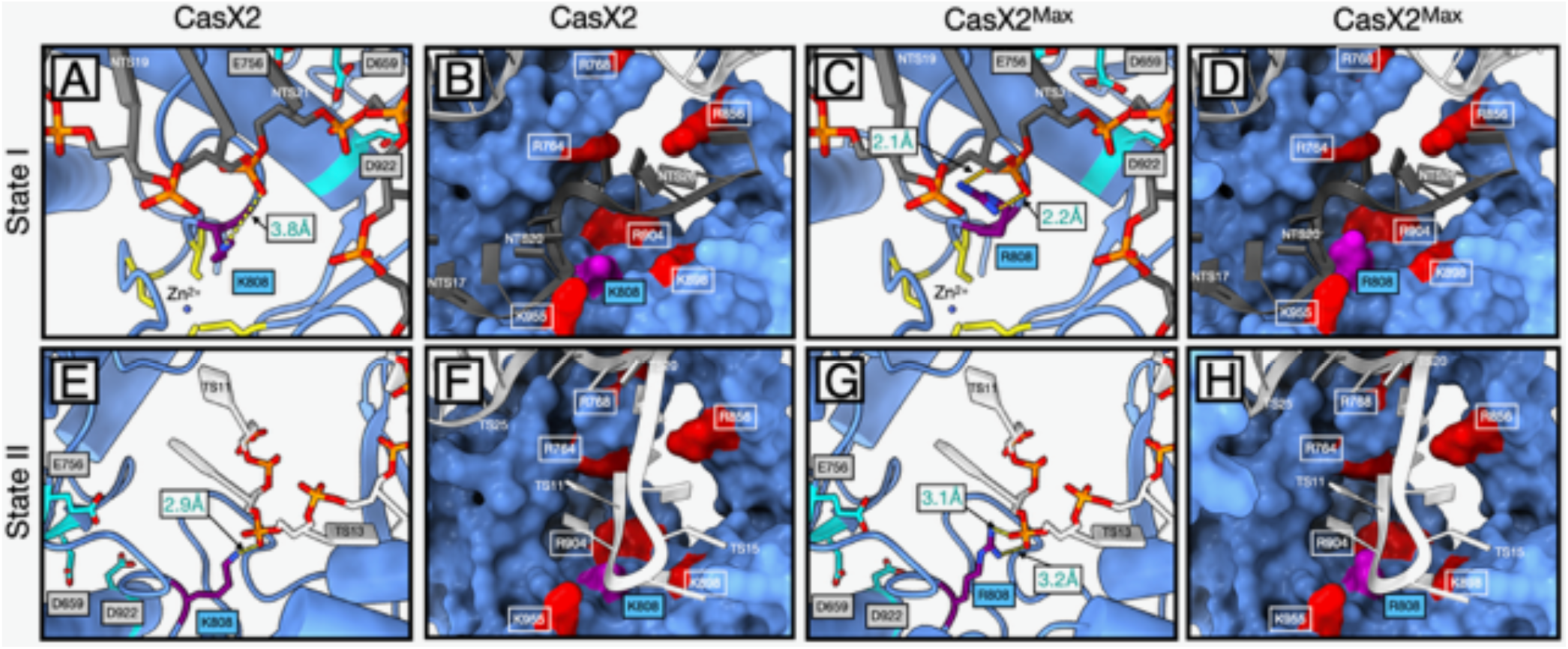
Structural context of K808R in CasX2 and CasX2^Max^ across catalytic states I and II. **A.** In CasX2 state I, K808 is positioned near the RuvC catalytic triad and the zinc ribbon motif. **B.** K808 forms a single 3.8 Å hydrogen bond with the NTS phosphate backbone at NTS20. It resides within a polar pocket composed of nearby basic residues that help stabilize the displaced NTS for catalysis. **C**. In CasX2^Max^ state I, the K808R substitution enables dual hydrogen bonds (2.0 Å and 2.2 Å) with the NTS phosphate at NTS20 increasing backbone contact compared to native CasX2. **D.** The K808R substitution also contributes a larger, more highly charged guanidinium group, subtly reshaping the local electrostatic environment within the same structural pocket. **E.** In CasX2 state II, K808 reorients toward the TS and forms a single 2.9 Å hydrogen bond with the phosphate backbone at TS12. **F.** The K808 residue remains within the same polar environment seen in state I, now positioned to support TS engagement during the cleavage step. **G.** In CasX2^Max^ state II, R808 engages the phosphate backbone at TS12 through dual hydrogen bonds (3.1 Å and 3.2 Å), increasing contact with the substrate strand. **H.** The additional positive charge and extended reach of the arginine side chain further enriches the local electrostatic surface, potentially enhancing retention of the TS near the RuvC active site. Residues and strands are colored as follows: K808 or R808 side chains (purple); basic polar residues (red); zinc-coordinating cysteines of the zinc ribbon motif (yellow); RuvC catalytic residues D659, E756, and D922 (cyan); DNA target strand (TS) (white); and non-target strand (NTS) (dark gray), with non-bridging oxygens (red) and phosphorous atoms (orange) to indicate the phosphodiester backbone.

Following NTS cleavage, CasX2 transitions to state II through a conformational rearrangement that involves downward bending of the sgRNA–DNA duplex and a rotation of the REC lobe (21, 37). This transition repositions the TS for cleavage and alters the orientation of the TSL domain, bringing K808 into proximity with the TS phosphate backbone. Although K808 remains embedded within the same polar surface as in state I, its side chain now contacts the TS rather than the NTS. However, its constrained geometry and mono-dentate hydrogen bonding limit this interaction to a single 2.9 Å hydrogen bond with TS12 (Figure 8E), providing only modest stabilization despite the favorable local charge environment (Figure 8F).

As in state I, the K808R substitution in CasX2^Max^ increases both the number and strength of contacts with the DNA backbone. R808 forms two hydrogen bonds (3.1 Å and 3.2 Å) with the non-bridging oxygens of base T12 (Figure 8G), enhancing local DNA engagement and positioning its guanidinium group within a concentrated cluster of basic residues near the RuvC catalytic cleft (Figure 8H). This configuration likely improves stabilization of the TS within the active site and contributes to the increased editing activity observed in cells.

### Zinc ribbon disruption and K808R compensation in state III

CasX2 state III is a catalytically inactive structural intermediate observed in ∼14% of CasX2– sgRNA complexes. In this conformation, the NTS remains positioned in the RuvC domain as in state I, but a disordered H2 domain disrupts contacts with the sgRNA scaffold stem and pseudoknot, exposing RNA and introducing a ∼20° change in sgRNA bend angle (21). This prevents proper stabilization of the sgRNA–DNA hybrid and misaligns the substrate for cleavage.

Structural comparisons between states I and III revealed localized shifts near residue 808, especially in the C-terminal Bridge Helix (7.2 ± 3.8 Å) and RuvC domain (3.2 ± 1.8 Å)—far greater than the average 0.56 ± 0.37 Å RMSD across the N-terminal region (Figure 9A). Catalytic residues D659, E756, and D922 were displaced by 3.3–5.7 Å (Figures 9A, Supplemental Figure S3). While the TSL domain core (residues 822–898) remained relatively stable (0.76 ± 0.46 Å), flanking segments containing the tetra-Cys zinc ribbon (C810, C813, C910, C912) shifted by 2.8–5.2 Å (Figure 9B). In state I and II, this motif coordinates a zinc atom and stabilizes nearby secondary elements (Figure 9B) (3, 38). However, in state III, C810 and C912 form a disulfide bond, displacing the zinc and collapsing the ribbon (Figure 9C). This redox-induced failure correlates with RMSD shifts >1.5 Å around the tetra-Cys cluster and disrupts local strand-guiding motifs (Figure 9D).

**Figure 9.**
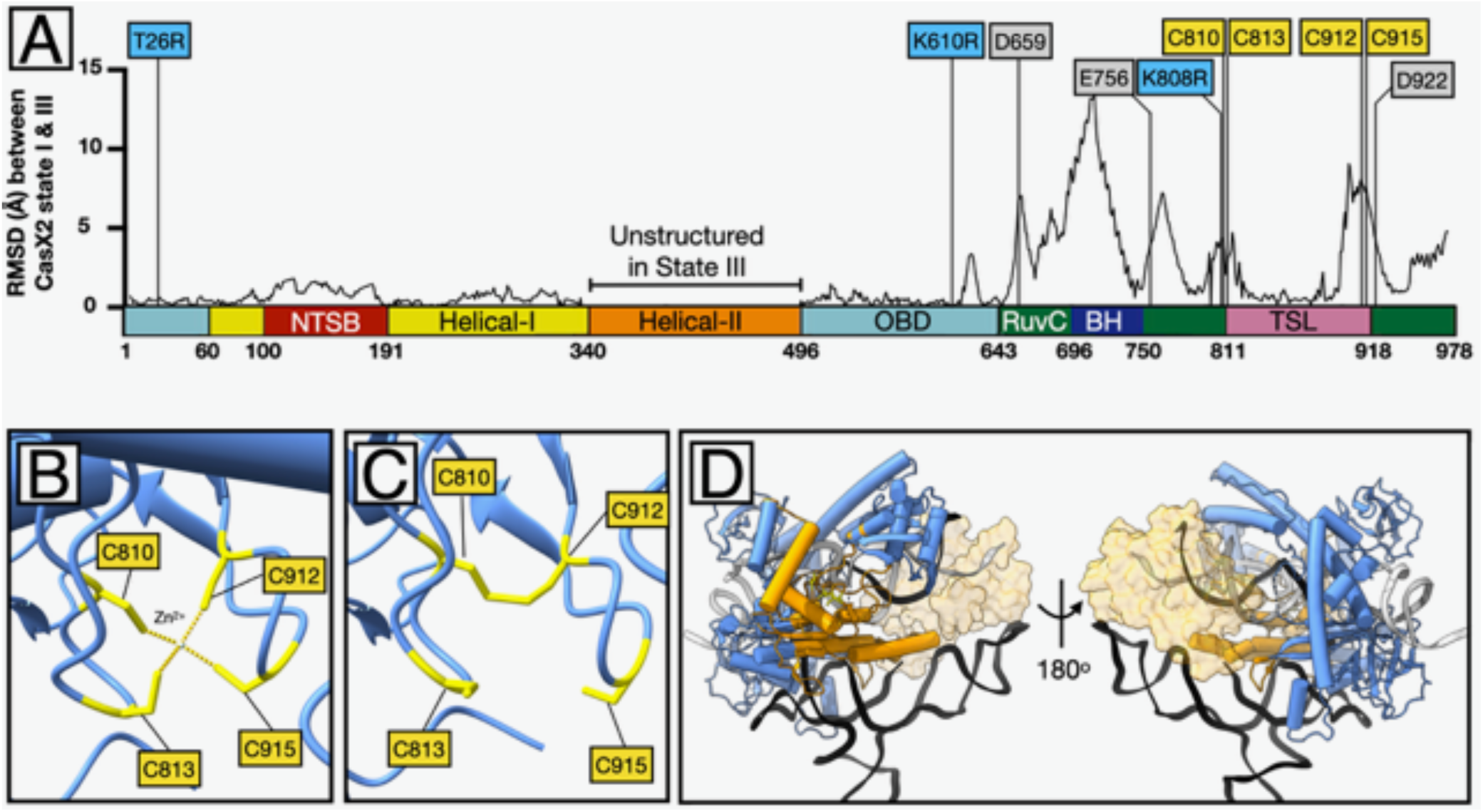
Zinc ribbon collapse in state III and associated structural shifts. **A.** Root mean square deviation (RMSD) values (Å) for α-carbon atoms across the CasX2 sequence, comparing state I and state III. Structural divergence is highest in the bridge helix (BH), RuvC, and TSL domains. Labeled positions indicate CasX2^Max^ substitutions (T26R, K610R, K808R) and selected catalytic and zinc-coordinating residues. Values for the Helical-II domain were excluded due to the absence of resolvable structure in this region in State III. **B.** In CasX2 state I, residues C810, C813, C912, and C915 coordinate a Zn²⁺ ion in a tetrahedral configuration, forming the basis of a stable zinc ribbon that anchors adjacent loops and helices. **C**. In state III, oxidation of C810 and C912 leads to disulfide bond formation, displacing the zinc ion and collapsing the zinc ribbon motif. This rearrangement disrupts the local scaffold, altering the surrounding loop geometry and destabilizing the structural support that normally anchors the adjacent RuvC and TSL domains. **D**. Structural features of CasX2 in state III shown as a ribbon-and-tube backbone model (blue), with bound sgRNA (black ribbon), non-target strand (NTS) (gray), and target strand (TS) (white). The four cysteines comprising the zinc ribbon motif (C810, C813, C912, C915) are shown (yellow). These cysteines form a disulfide bond between C810 and C912 in state III. Residues undergoing >2 Å displacement relative to state I are shown in orange, revealing that the majority of structural rearrangements are concentrated around the oxidized zinc ribbon cysteines. The only major exception is the bridge helix, that connects to the unresolved Helical-II domain. To provide spatial context, the position of the Helical-II domain from state I is shown (transparent tan surface).

K808, two residues upstream of C810, is directly affected by the oxidation-induced disruption of the zinc ribbon motif. In state I, it forms a 3.0 Å electrostatic interaction with the NTS backbone, but shifts 4.3 Å away in state III (Supplemental Figure S4A). The K808R substitution in CasX2Max restores this contact in state III, forming a 2.3 Å bond (Supplemental Figure S4B). This added stabilization likely helps retain the NTS near the catalytic site and lowers the barrier for recovery to the active state I conformation.

## Discussion

CRISPR/Cas12e (CasX) nucleases represent a compact and programmable class of genome editing tools that offer several advantages over the more common Cas9 editors. Several Cas12e enzymes, including CasX1 and CasX2, are smaller than Cas9, facilitating their packaging and delivery *in vivo* using viral vectors (39). Moreover, they derive from the non-pathogenic bacteria, *Deltaproteobacteria* (CasX1), and *Planctomycetes* (CasX2), to which no pre-existing immunity in humans has been reported. In contrast, *Sa*Cas9 and *Sp*Cas9 derive from the common human pathogens *S. aureus* and *S. pyogenes*, respectively, and reports of pre-existing immunity to proteins from these bacteria, including Cas9, have been described, limiting their repeated use as a therapeutic in humans (40, 41). Additional benefits to these Cas12e enzymes include a more restricted PAM sequence requirement, and the generation of a staggered cleavage cut that could favor HDR and MMEJ repair mechanisms in cells.

In this manuscript, we assessed the efficiency of CasX2 cleavage of *CCR5* in cells. We first assessed cleavage of *CCR5* using a plasmid target transfected into HEK293-FT cells. For cleavage analysis, we selected two sgRNAs, sg5 and sg10, that we showed previously were able to cleave an isolated *CCR5* DNA target (5). These sgRNAs targeted two regions in *CCR5* that overlapped two established gRNAs for *Sa*Cas9 reported by Dash et al. (32). Because our previous study showed variable cleavage efficiencies of isolated DNA using sgRNAs with different spacer lengths, we sought to determine whether spacer lengths also impacted *CCR5* cleavage in cells. Similar to our earlier findings with the isolated *CCR5* target (5), sg5 and sg10 displayed variable cleavage efficiencies at spacer lengths of 20 nt and 23 nt in cells. However, we observed minimal cleavage using a spacer of 17 nt. It is likely that spacer sequences for CasX2 that are smaller than 20 nt have reduced efficiency in cells, compared to cleavage of isolated DNA.

Despite the finding that CasX2 readily cleaved the plasmid target expressing *CCR5* in cells, we found that CasX2 showed no appreciable cleavage of genomic *CCR5* relative to *Sa*Cas9 in mammalian cells (1, 21, 42). Whether this is due to a reduced access to the target by the spacer region, it is notable that *Sa*Cas9 showed efficient cleavage of this genomic target, suggesting that other factors may impact CasX2 cleavage of a genomic target. Mechanistic studies have suggested that this inefficiency could arise from the structural properties of the H2 domain, a flexible element unique to CasX2. Tsuchida et al. proposed that H2 domain mobility interferes with stable R-loop formation, allowing the RNP to transition into a pre-cleavage conformation termed state III, which prevents DNA cleavage (21). When evaluating cleavage activity of CasX2 enzymes, it is important to distinguish between results obtained utilizing isolated DNA targets, and those using DNA targets in cells.

The poor cleavage of genomic *CCR5* by the native form of CasX2 prompted us to compare CasX2 to a newly identified modified editor, CasX2^Max^. Developed through the MIDAS engineering protocol, CasX2^Max^ was reported to achieve a >100-fold improvement in editing efficiency at the *CCR5* locus in cells (1). These reported improvements in cellular editing efficiency likely stem from structural alterations that improve R-loop stability and reduce kinetic trapping in non-productive conformations, rather than from changes in intrinsic cleavage rates. Our structural and functional analysis suggests that these mutations in CasX2^Max^, although not located within the H2 domain, indirectly compensate for its destabilizing effects by stabilizing the R-loop and extending the duration during which single-stranded DNA remains aligned with the guide RNA. In particular, the modifications T26R and K610R, strengthen DNA strand separation at the PAM-proximal fork, while the modification K808R reinforces interactions with the NTS in state I and the TS in state II, ensuring alignment within the RuvC catalytic site. These stabilizing effects may reduce the likelihood of kinetic trapping in state III and improve transition efficiency into the cleavage-competent conformations.

Our re-analysis of cryo-EM data from Tsuchida et al. (21) revealed that the catalytically inactive state III was associated with oxidation-induced disruption of the zinc ribbon motif near the RuvC active site, suggesting that CasX2 activity is redox-sensitive. Structural collapse of this motif and adjacent elements impairs NTS engagement and reduces cleavage activity, supporting the idea that oxidizing conditions can shift the enzyme toward an inactive conformation. As *Plm*Cas12e (CasX2) was recently shown to possess robust trans-nuclease activity (39), this redox sensitivity may be particularly relevant for *in vitro* diagnostic applications like DETECTR (43) and SHERLOCK (44), where reduced enzymatic turnover could limit sensitivity and reaction kinetics.

Analyses of the cleavage events and cellular repair mechanisms in cells following either CasX2 or CasX2^Max^ cleavage of genomic *CCR5* showed differences in the types of indels following cleavage of the plasmid *CCR5* target. Our analyses revealed differences in the most common indel as determined by Nanopore sequencing, with insertions of nucleotides as the most common mutations detected following CasX2 cleavage, but not CasX2^Max^ at all spacer lengths tested. While these differences may indicate changes in cleavage type and overhang length, they may also be attributable to changes in the enzyme’s target site residency after cleavage, which may impact DNA repair pathway selection. Future studies should investigate the impact of the alterations in CasX2^Max^ on substate binding, cleavage site, and the interaction of these with DSB repair by host cell machinery. In the case of CasX2^Max^ targeting genomic *CCR5*, we observed 16 - 32 bp deletions that were consistent with MMEJ for both target sites tested. Identical deletions were observed following transfection of sgRNAs with either the 20 or 23 nt spacer lengths, suggesting similar DNA repair mechanisms were triggered. However, CRISPR events that generate different overhangs could result in the same deletion following DNA repair. Therefore, it is not clear whether the spacer length impacts the overhang generated by CasX2 or CasX2^Max^ in the same manner as was previously described for CasX1 *in vitro* (6).

Our results confirm the improved potential of the CasX2 variant, CasX2^Max^, developed by Chen et al. through their MIDAS enzyme engineering approach (1). CasX2 was already an appealing candidate enzyme for CRISPR therapeutics due to the protein’s compact size, use of alternate PAM sequence, and presumed lack of pre-existing immunity in contrast to more widely used enzymes derived from relevant human pathogens. We show that CasX2^Max^ also exhibits efficacy comparable to *Sa*Cas9 in targeting *CCR5* in human cells. We further show, using a single sgRNA, that CasX2^Max^ produced disruptive MMEJ-mediated deletions at target sites that may prove beneficial for gene disruption approaches, including targeting of *CCR5*. Further, CasX2^Max^, with paired sgRNAs, was effective at mediating excision of the intervening genomic region between two protospacers. Taken together, these results support continued development of CasX enzymes in the development of CRISPR therapeutics.

## Author contributions statement

Christine A. Hodge prepared and revised the figures in this manuscript based on her experimental work, and made significant changes during the proof-reading stage.

Niles P. Donegan has made a significant contribution to the work reported. He substantially wrote, revised and critically reviewed the article, and prepared and revised the figures in the manuscript based on his experimental work.

David A. Armstrong has made a significant contribution to the work reported. He provided data supporting the experimental design and approach taken in this manuscript.

Matthew S. Hayden drafted, wrote and substantially revised and critically reviewed the article, and made significant changes at the proof-reading stage. He agrees to take responsibility and to be held accountable for the contents of the article.

Alexandra L. Howell drafted, wrote and substantially revised and critically reviewed the article, and agreed on the journal for submission, and made significant changes at the proof-reading stage. She agrees to take responsibility and to be held accountable for the contents of the article.

---

**Supplemental Figure S1.**
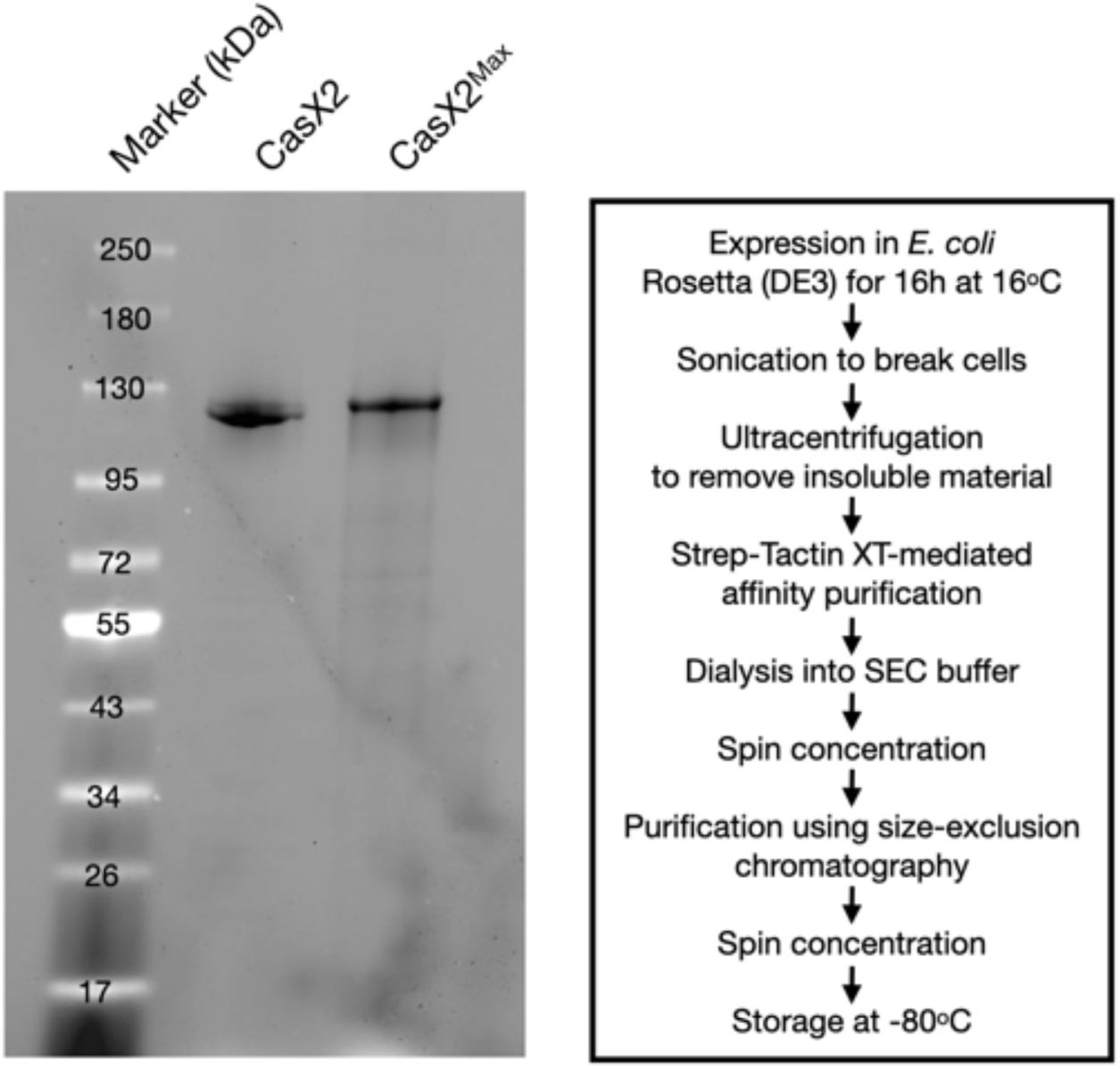
SDS-PAGE analysis of purified CasX2 and CasX2^Max^. **A.** Stain-Free gel showing purified recombinant CasX2 and CasX2^Max^, each bearing an N-terminal SV40 nuclear localization signal (NLS) and C-terminal nucleoplasmin NLS, 3× hemagglutinin (HA) epitope tag, and Twin-Strep tag. Both proteins migrate near their expected molecular weight (∼123 kDa) with no detectable degradation products or significant contaminants. Molecular weight markers (kDa) are shown at left. **B**. Workflow for CasX2 and CasX2^Max^ protein expression and purification. CasX2 and CasX2^Max^ were expressed in *E. coli* Rosetta (DE3) cells for 16 h at 16°C following IPTG induction in Terrific Broth. Cells were lysed by sonication after nuclease treatment, and lysates were clarified by ultracentrifugation at 50,000 *× g*. Soluble protein was purified via Strep-Tactin XT affinity chromatography, dialyzed into SEC buffer, and concentrated using 50 kDa MWCO spin concentrators. Further purification was performed by size-exclusion chromatography on a Superdex 200 column. Final CasX2 preparations were concentrated, snap-frozen in liquid nitrogen, and stored at −80°C.

**Supplemental Figure S2.**
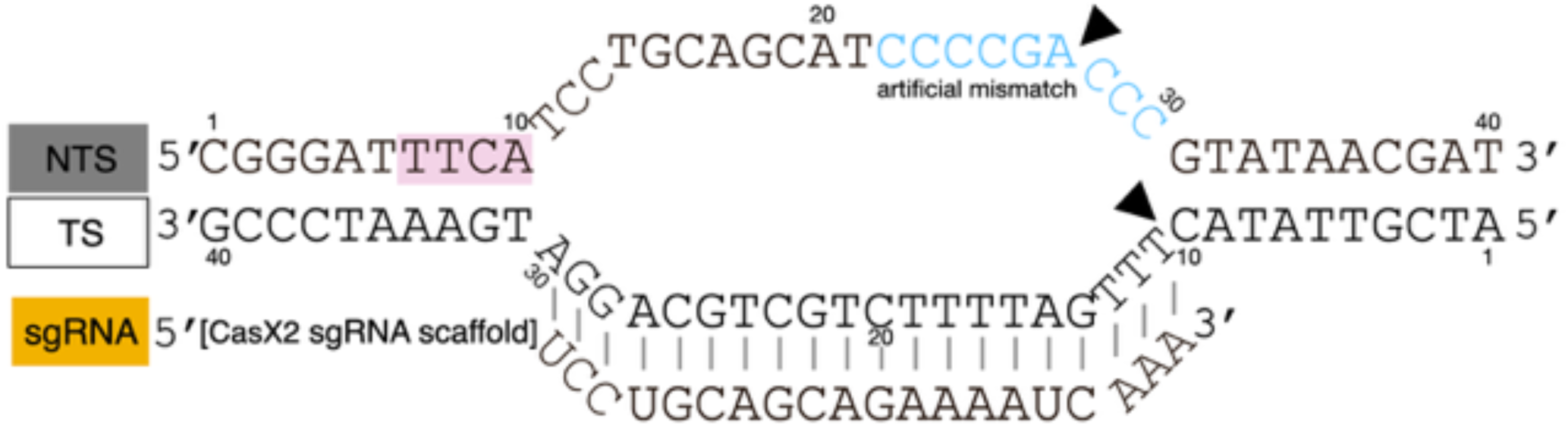
Reference sequence and base numbering used for structural models and interaction mapping. Schematic of the CasX2 target duplex and guide RNA used throughout this study, adapted from the protospacer sequence resolved in cryo-EM structures (e.g., PDB: 7WAY). The non-target strand (NTS) and target strand (TS) are shown with base positions numbered 1–40 for consistency across all structural figures. The 5′ end of the sgRNA spacer hybridizes to the TS and extends toward the sgRNA scaffold (not shown). The PAM sequence (TTCA) is highlighted in pink. Transversion mutations in the NTS (blue) introduce a local mismatch that destabilizes the duplex and promotes R-loop formation for cryo-EM analysis. Triangles (black) indicate the resolved cleavage sites on the NTS and TS observed in structural states I and II. Color blocks to the left denote the standard strand color scheme used throughout the manuscript: NTS (dark gray), TS (white), and sgRNA (gold).

**Supplemental Figure S3.**
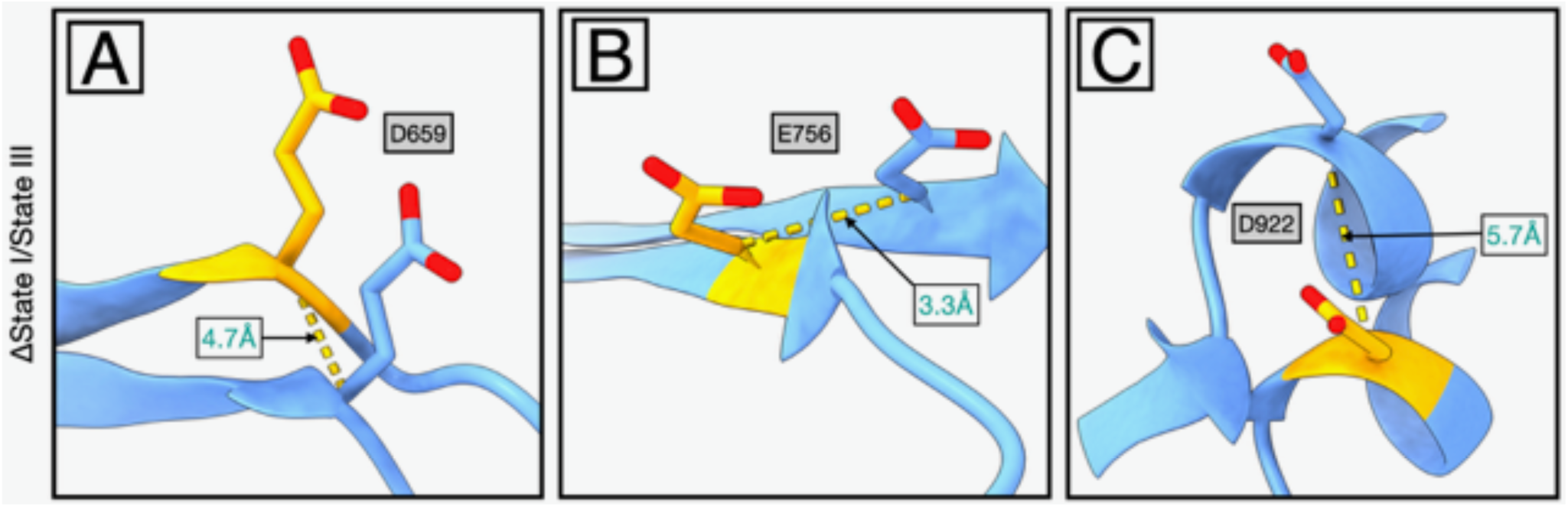
Positional displacement of CasX2 catalytic residues between active and inactive conformations. The positions of the RuvC catalytic triad residues in CasX2 are shown in the active conformation (state I, blue) and the catalytically inactive conformation (state III, yellow). Superimposed backbone models highlight α-carbon shifts for (**A**) D659 (4.7 Å), (**B**) E756 (3.3 Å), and (**C**) D922 (5.7 Å), reflecting localized remodeling of the RuvC domain during the transition to state III.

**Supplemental Figure S4.**
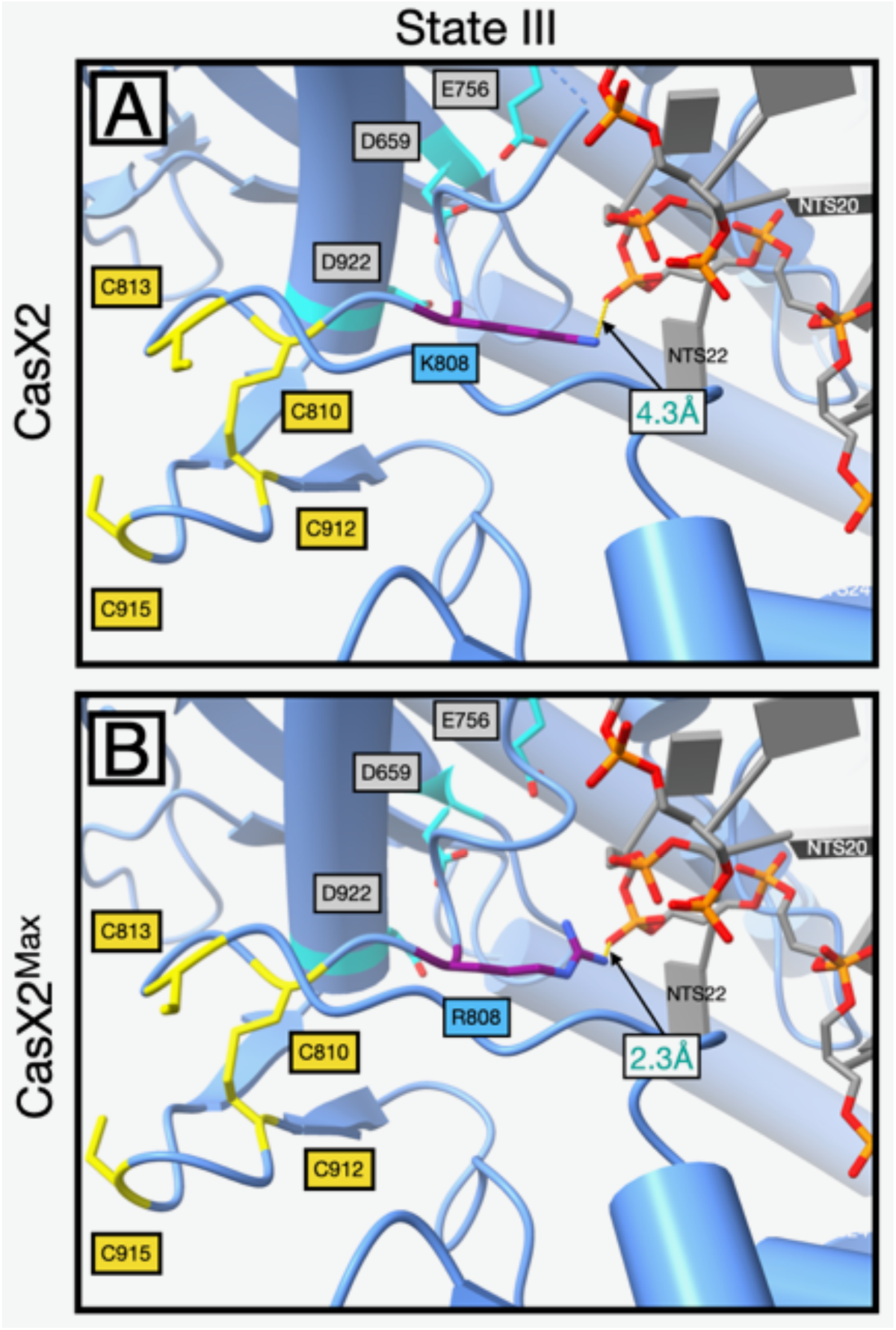
K808R substitution restores NTS phosphate contact in the inactive state III conformation. **A.** In native CasX2, state III rearrangements disrupt the zinc ribbon and reposition K808, increasing its distance to the NTS phosphate at position 22 to 4.3 Å. **B.** In CasX2^Max^, the K808R substitution reduces this distance to 2.3 Å, maintaining contact with the NTS backbone despite the same state III conformation. Residues and strands are colored as follows: K808 or R808 side chains, purple; zinc-coordinating cysteines of the zinc ribbon motif (C810, C813, C912, C915), yellow; RuvC catalytic residues D659, E756, and D922, cyan; DNA non-target strand (NTS), dark gray, with phosphodiester backbone atoms shown as phosphorous (orange) and non-bridging oxygens (red).

**Supplementary Table 1 (Table S1):** List of gRNA spacer sequences used.

**Supplementary Table 2 (Table S2):** List of sgRNA scaffold sequences used.

**Supplementary Table 3 (Table S3):** List of plasmids used with complete plasmid sequence.

**Supplementary Table 4 (Table S4):** List of oligonucleotides used for cloning of gRNAs.

**Supplementary Table 5 (Table S5):** List of PCR primers used in Figures 1, 3, 4, 5 and 6.

**Supplementary Table 6 (Table S6):** List of PCR amplification conditions used in Figures 1, 3, 4, 5 and 6.

## Data Availability Statement

<u>Location of research data associated with this paper</u>: The data from this manuscript is available within this article and the supplemental materials.

<u>Under what conditions the data can be accessed</u>: Data can be accessed from the manuscript’s Figures and Tables. Any underlying data will be made available upon reasonable request.

