## Supplementary material for "ENHANCED CLEAVAGE OF GENOMIC *CCR5* USING CASX2^Max^": Supplmental Table S1

### Supplemental Table S1

#### gRNA Spacer Sequences

| CasX2 sgRNA | Target Description | cDNA of Spacer Sequence | Reference |
|---|---|---|---|
| sg5 (17 nt) | nucleotide positions 272 to 288 in CCR5 | AAAGTCCCACTGGGCGG | Armstrong et al. 2023 |
| sg5 (20 nt) | nucleotide positions 269 to 288 in CCR5 | AAAGTCCCACTGGGCGGCAG | Armstrong et al. 2023 |
| sg5 (23 nt) | nucleotide positions 266 to 288 in CCR5 | AAAGTCCCACTGGGCGGCAGCAT | Armstrong et al. 2023 |
| sg10 (17 nt) | nucleotide positions 1021 to 1037 in CCR5 | TGCTCCCCAGTGGATCG | Armstrong et al. 2023 |
| sg10 (20 nt) | nucleotide positions 1018 to 1037 in CCR5 | TGCTCCCCAGTGGATCGGGT | Armstrong et al. 2023 |
| sg10 (23 nt) | nucleotide positions 1015 to 1037 in CCR5 | TGCTCCCCAGTGGATCGGGTGTA | Armstrong et al. 2023 |
| E6 (20 nt) | nucleotide positions 216 to 235 in GFP | TGTGGTCGGGGTAGCGGCTG | US Patent Application US20220220508A1 |

| SaCas9 sgRNA | | Sequence | Reference |
|---|---|---|---|
| CCR5-A (20 nt) | nucleotide positions 225 to 275 in CCR5 | CGGCAGCATAGTGAGCCCAG | Dash et al. 2023 |
| CCR5-B (20 nt) | nucleotide positions 1009 to1028 in CCR5 | TCAGTTTACACCCGATCCAC | Dash et al. 2023 |

| CasX2$^{Max}$ sgRNA in *in silico* modeling | | Sequence | Reference |
|---|---|---|---|
| CasX2 sgRNA (20 nt) | Fifth protospacer downstream of the DpbCas12e (CasX1) ORF (Accession KU516152.1) | TCCTGCAGCAGAAAATCAAA | Tsuchida et al 2022 |
