## Supplementary material for "ENHANCED CLEAVAGE OF GENOMIC *CCR5* USING CASX2^Max^": Supplmental Table S2

### Supplemental Table S2

#### sgRNA scaffold sequences

| **ID** | **Description** | **Description** | **Reference** |
|---|---|---|---|
| CasX2 sgRNA | Native CasX2 sgRNA (sgRNAv2) | GUACUGGCGCUUUUAUCUCAUUACUUUGAGAGCCAUCACCAGCGACUAUGUCGUAUGGGUAAAGCGCUUAUUUAUCGGAGAGAAAUCCGAUAAAUAAGAAGCAUCAAAG | Tsuchida et al 2022 |
| CasX1 sgRNA | Native CasX1 sgRNA (sgRNAv1) | GGCGCGUUUAUUCCAUUACUUUGGAGCCAGUCCCAGCGACUAUGUCGUAUGGACGAAGCGCUUAUUUAUCGGAGAGAAACCGAUAAGUAAAACGCAUCAAAGUCCUGCAGCAGAAAAUCAAA | Burstein et al 2017 |
| SaCas9 sgRNA | Native SaCas9 sgRNA | GUUUUAGUACUCUGGAAACAGAAUCUACUAAAACAAGGCAAAAUGCCGUGUUUAUCUCGUCAACUUGUUGGCGAGA | Ran et al 2015 |
