## Supplementary material for "ENHANCED CLEAVAGE OF GENOMIC *CCR5* USING CASX2^Max^": Supplmental Table S3

### Supplemental Table S3

### Plasmid Sequences

**>CCR5_TTCC_target**

gacggatcgggagatctcccgatcccctatggtgcactctcagtacaatctgctctgatgccgcatagttaagccag
tatctgctccctgcttgtgtgttggaggtcgctgagtagtgcgcgagcaaaatttaagctacaacaaggcaaggctt
gaccgacaattgcatgaagaatctgcttagggttaggcgttttgcgctgcttcgcgatgtacgggccagatatacgc
gttgacattgattattgactagttattaatagtaatcaattacggggtcattagttcatagcccatatatggagttc
cgcgttacataacttacggtaaatggcccgcctggctgaccgcccaacgacccccgcccattgacgtcaataatgac
gtatgttcccatagtaacgccaatagggactttccattgacgtcaatgggtggagtatttacggtaaactgcccact
tggcagtacatcaagtgtatcatatgccaagtacgccccctattgacgtcaatgacggtaaatggcccgcctggcat
tatgcccagtacatgaccttatgggactttcctacttggcagtacatctacgtattagtcatcgctattaccatggt
gatgcggttttggcagtacatcaatgggcgtggatagcggtttgactcacggggatttccaagtctccaccccattg
acgtcaatgggagtttgttttggcaccaaaatcaacgggactttccaaaatgtcgtaacaactccgccccattgacg
caaatgggcggtaggcgtgtacggtgggaggtctatataagcagagctctctggctaactagagaacccactgctta
ctggcttatcgaaattaatacgactcactatagggagacccaagctggctagcgccaccatggattatcaagtgtca
agtccaatctatgacatcaattatatacatcggagccctgccaaaaaatcaatgGgaagcaaatcgcagcccgcct
cctgcctccgctctactcactggtgttcatctttggttttgtgggcaacatgctggtcatcctcatcctgataaact
gcaaaaggcGaagagcatgactgacatctacctgctcaacctggccatctctgacctgtttttccttcCtactgtcc
ccttcCgggctcactatgctgccgcccagtgggactttggaaatacaatgtgtcaactcttgacagggctctatttt
ataggcttcttctctggaatcttcttcatcatcctcctgacaatcgataggtacctggctgtcgtccatgctgtgtt
tgctttaaaagccaggacggtcacctttggggtggtgacaagtgtgatcacttgggtggtggctgtgtttgcgtctc
tcccaggaatcatctttaccagatctcaaaaagaaggtcttcattacacctgcagctctcattttccatacagtcag
tatcaattctggaagaatttccagacattaaagatagtcatcttggggctggtcctgccgctgcttgtcatggtcat
ctgctactcgggaatcctaaaaactctgcttcggtgtcgaaatgagaagaagaggcacagggctgtgaggcttatct
tcaccatcatgattgtttattttctcttcCgggctccctacaacattgtccttctcctgaacaccttccaggaattc
tttggccGgaataattgcagtagctctaacaggttggaccaagctatgcaggtgacagagactcttgggatgacgca
ctgctgcatcaaccccatcatctatgcctttgtcggggagaagttcagaaactacctcttagtcttcttccaaaagc
acattgccaaacgcttctgcaaatgctgttcCattttccagcaagaggctcccgagcgagcaagctcagtttacacc
cgatccactggggagcaggaaatatctgtgggcttgtgataactcgagtctagagggcccgtttaaacccgctgatc
agcctcgactgtgccttctagttgccagccatctgttgtttgcccctcccccgtgccttccttgaccctggaaggtg
ccactcccactgtcctttcctaataaaatgaggaaattgcatcgcattgtctgagtaggtgtcattctattctgggg
ggtggggtggggcaggacagcaagggggaggattgggaagacaatagcaggcatgctggggatgcggtgggctctat
ggcttctgaggcggaaagaaccagctggggctctagggggtatccccacgcgccctgtagcggcgcattaagcgcgg
cgggtgtggtggttacgcgcagcgtgaccgctacacttgccagcgccctagcgcccgctcctttcgctttcttccct
tcctttctcgccacgttcgccggctttccccgtcaagctctaaatcgggggctccctttagggttccgatttagtgc
tttacggcacctcgaccccaaaaaacttgattagggtgatggttcacgtagtgggccatcgccctgatagacggttt
ttcgccctttgacgttggagtccacgttctttaatagtggactcttgttccaaactggaacaacactcaaccctatc
tcggtctattcttttgatttataagggattttgccgatttcggcctattggttaaaaaatgagctgatttaacaaaa
atttaacgcgaattaattctgtggaatgtgtgtcagttagggtgtggaaagtccccaggctccccagcaggcagaag
tatgcaaagcatgcatctcaattagtcagcaaccaggtgtggaaagtccccaggctccccagcaggcagaagtatgc
aaagcatgcatctcaattagtcagcaaccatagtcccgcccctaactccgcccatcccgcccctaactccgcccagt
tccgcccattctccgccccatggctgactaattttttttatttatgcagaggccgaggccgcctctgcctctgagct
attccagaagtagtgaggaggcttttttggaggcctaggcttttgcaaaaagctcccgggagcttgtatatccattt
tcggatctgatcaagagacaggatgaggatcgtttcgcatgattgaacaagatggattgcacgcaggttctccggcc
gcttgggtggagaggctattcggctatgactgggcacaacagacaatcggctgctctgatgccgccgtgttccggct
gtcagcgcaggggcgcccggttctttttgtcaagaccgacctgtccggtgccctgaatgaactgcaggacgaggcag
cgcggctatcgtggctggccacgacgggcgttccttgcgcagctgtgctcgacgttgtcactgaagcgggaaggac

tggctgctattgggcgaagtgccggggcaggatctcctgtcatctcaccttgctcctgccgagaaagtatccatcat
ggctgatgcaatgcggcggctgcatacgcttgatccggctacctgcccattcgaccaccaagcgaaacatcgcatcg
agcgagcacgtactcggatggaagccggtcttgtcgatcaggatgatctggacgaagagcatcaggggctcgcgcca
gccgaactgttcgccaggctcaaggcgcgcatgcccgacggcgaggatctcgtcgtgacccatggcgatgcctgctt
gccgaatatcatggtggaaaatggccgcttttctggattcatcgactgtggccggctgggtgtggcggaccgctatc
aggacatagcgttggctacccgtgatattgctgaagagcttggcggcgaatgggctgaccgcttcctcgtgctttac
ggtatcgccgctcccgattcgcagcgcatcgccttctatcgccttcttgacgagttcttctgagcgggactctgggg
ttcgaaatgaccgaccaagcgacgcccaacctgccatcacgagatttcgattccaccgccgccttctatgaaaggtt
gggcttcggaatcgttttccgggacgccggctggatgatcctccagcgcggggatctcatgctggagttcttcgccc
accccaacttgtttattgcagcttataatggttacaaataaagcaatagcatcacaaatttcacaaataaagcattt
ttttcactgcattctagttgtggtttgtccaaactcatcaatgtatcttatcatgtctgtataccgtcgacctctag
ctagagcttggcgtaatcatggtcatagctgtttcctgtgtgaaattgttatccgctcacaattccacacaacatac
gagccggaagcataaagtgtaaagcctggggtgcctaatgagtgagctaactcacattaattgcgttgcgctcactg
cccgctttccagtcgggaaacctgtcgtgccagctgcattaatgaatcggccaacgcgcggggagaggcggtttgcg
tattgggcgctcttccgcttcctcgctcactgactcgctgcgctcggtcgttcggctgcggcgagcggtatcagctc
actcaaaggcggtaatacggttatccacagaatcaggggataacgcaggaaagaacatgtgagcaaaaggccagcaa
aaggccaggaaccgtaaaaaggccgcgttgctggcgtttttccataggctccgcccccctgacgagcatcacaaaaa
tcgacgctcaagtcagaggtggcgaaacccgacaggactataaagataccaggcgtttccccctggaagctccctcg
tgcgctctcctgttccgaccctgccgcttaccggatacctgtccgcctttctcccttcgggaagcgtggcgctttct
catagctcacgctgtaggtatctcagttcggtgtaggtcgttcgctccaagctgggctgtgtgcacgaaccccccgt
tcagcccgaccgctgcgccttatccggtaactatcgtcttgagtccaacccggtaagacacgacttatcgccactgg
cagcagccactggtaacaggattagcagagcgaggtatgtaggcggtgctacagagttcttgaagtggtggcctaac
tacggctacactagaagaacagtatttggtatctgcgctctgctgaagccagttaccttcggaaaaagagttggtag
ctcttgatccggcaaacaaaccaccgctggtagcggtttttttgtttgcaagcagcagattacgcgcagaaaaaaag
gatctcaagaagatcctttgatcttttctacggggtctgacgctcagtggaacgaaaactcacgttaagggattttg
gtcatgagattatcaaaaaggatcttcacctagatccttttaaattaaaaatgaagttttaaatcaatctaaagtat
atatgagtaaacttggtctgacagttaccaatgcttaatcagtgaggcacctatctcagcgatctgtctatttcgtt
catccatagttgcctgactccccgtcgtgtagataactacgatacgggagggcttaccatctggccccagtgctgca
atgataccgcgagacccacgctcaccggctccagatttatcagcaataaaccagccagccggaagggccgagcgcag
aagtggtcctgcaactttatccgcctccatccagtctattaattgttgccgggaagctagagtaagtagttcgccag
ttaatagtttgcgcaacgttgttgccattgctacaggcatcgtggtgtcacgctcgtcgtttggtatggcttcattc
agctccggttcccaacgatcaaggcgagttacatgatcccccatgttgtgcaaaaaagcggttagctccttcggtcc
tccgatcgttgtcagaagtaagttggccgcagtgttatcactcatggttatggcagcactgcataattctcttactg
tcatgccatccgtaagatgcttttctgtgactggtgagtactcaaccaagtcattctgagaatagtgtatgcggcga
ccgagttgctcttgcccggcgtcaatacgggataataccgcgccacatagcagaactttaaaagtgctcatcattgg
aaaacgttcttcggggcgaaaactctcaaggatcttaccgctgttgagatccagttcgatgtaacccactcgtgcac
ccaactgatcttcagcatcttttactttcaccagcgtttctgggtgagcaaaaacaggaaggcaaaatgccgcaaaa
aagggaataagggcgacacggaaatgttgaatactcatactcttcctttttcaatattattgaagcatttatcaggg
ttattgtctcatgagcggatacatatttgaatgtatttagaaaaataaacaaataggggttccgcgcacatttcccc
gaaaagtgccacctgacgtc

**>CCR5_TTCC_target**

TAGAAAAGATCAAAGGATCTTCTTGAGATCCTTTTTTTCTGCGCGTAATCTGCTGCTTGCAAACAAAAAAACCACCG
CTACCAGCGGTGGTTTGTTTGCCGGATCAAGAGCTACCAACTCTTTTTCCGAAGGTAACTGGCTTCAGCAGAGCGCA
GATACCAAATACTGTTCTTCTAGTGTAGCCGTAGTTAGGCCACCACTTCAAGAACTCTGTAGCACCGCCTACATACC
TCGCTCTGCTAATCCTGTTACCAGTGGCTGCTGCCAGTGGCGATAAGTCGTGTCTTACCGGGTTGGACTCAAGACGA
TAGTTACCGGATAAGGCGCAGCGGTCGGGCTGAACGGGGGGTTCGTGCACACAGCCCAGCTTGGAGCGAACGACCTA

CACCGAACTGAGATACCTACAGCGTGAGCTATGAGAAAGCGCCACGCTTCCCGAAGGGAGAAAGGCGGACAGGTATC
CGGTAAGCGGCAGGGTCGGAACAGGAGAGCGCACGAGGGAGCTTCCAGGGGGAAACGCCTGGTATCTTTATAGTCCT
GTCGGGTTTCGCCACCTCTGACTTGAGCGTCGATTTTTGTGATGCTCGTCAGGGGGGCGGAGCCTATGGAAAAACGC
CAGCAACGCGGCCTTTTTACGGTTCCTGGCCTTTTGCTGGCCTTTTGCTCACATGTCCTGCAGGCAGCTGCGCGCTC
GCTCGCTCACTGAGGCCGCCCGGGCGTCGGGCGACCTTTGGTCGCCCGGCCTCAGTGAGCGAGCGAGCGCGCAGAGA
GGGAGTGGCCAACTCCATCACTAGGGGTTCCTGCGGCCTCTAGACTCGAGGCGTTGACATTGATTATTGACTAGTTA
TTAATAGTAATCAATTACGGGGTCATTAGTTCATAGCCCATATATGGAGTTCCGCGTTACATAACTTACGGTAAATG
GCCCGCCTGGCTGACCGCCCAACGACCCCCGCCCATTGACGTCAATAATGACGTATGTTCCCATAGTAACGCCAATA
GGGACTTTCCATTGACGTCAATGGGTGGAGTATTTACGGTAAACTGCCCACTTGGCAGTACATCAAGTGTATCATAT
GCCAAGTACGCCCCCTATTGACGTCAATGACGGTAAATGGCCCGCCTGGCATTATGCCCAGTACATGACCTTATGGG
ACTTTCCTACTTGGCAGTACATCTACGTATTAGTCATCGCTATTACCATGGTGATGCGGTTTTGGCAGTACATCAAT
GGGCGTGGATAGCGGTTTGACTCACGGGGATTTCCAAGTCTCCACCCCATTGACGTCAATGGGAGTTTGTTTTGGCA
CCAAAATCAACGGGACTTTCCAAAATGTCGTAACAACTCCGCCCCATTGACGCAAATGGGCGGTAGGCGTGTACGGT
GGGAGGTCTATATAAGCAGAGCTCTCTGGCTAACTACCGGTGCCACCATGTCCGGATCCCCTGCTGCCAAGAGGGTC
AAGTTGGACATGCAAGAGATCAAGAGAATCAACAAGATCAGAAGGAGACTGGTCAAGGAcagcaacacaaagaaggc
cggcaagacaggccccatgaaaaccctgctcgtcagagtgatgacccctgacctgagagagcggctggaaaacctga
gaaagaagcccgagaacatccctcagcctatcagcaacaccagcagggccaacctgaacaagctgctgaccgactac
accgagatgaagaaagccatcctgcacgtgtactgggaagagttccagaaagaccccgtgggcctgatgagcagagt
tgctcagcccgctcctaagaacatcgaccagagaaagctgatccccgtgaaggacggcaacgagagactgacctcta
gcggctttgcctgcagccagtgttgccagcctctgtacgtgtacaagctggaacaagtgaacgacaagggcaagccc
cacaccaactacttcggcagatgcaacgtgtccgagcacgagaggctgatcctgctgtctcctcacaagcccgaggc
caacgatgagctggtcacatacagcctgggcaagttcggacagagagccctggacttctacagcatccacgtgacca
gggagagcaatcaccctgtgaagcccctggaacagatcggcggcaatagctgtgcctctggacctgtgggaaaagcc
ctgagcgacgcctgtatgggagccgtggcatccttcctgaccaagtaccaggacatcatcctggaacaccagaaagt
gatcaagaagaacgagaaaagactggccaacctcaaggatatcgccagcgctaacggcctggcctttcctaagatca
ccctgcctccacagcctcacaccaaagagggcatcgaggcctacaacaacgtggtggcccagatcgtgatttgggtc
aacctgaatctgtggcagaagctgaagatcggcagggacgaagccaagccactgcagagactgaagggcttccctag
cttccctctggtggaaagacaggccaatgaagtggattggtgggacatggtctgcaacgtgaagaagctgatcaacg
agaagaaagaggatggcaaggttttctggcagaacctggccggctacaagagacaagaagccctgctgccttacctg
agcagcgaagaggaccggaagaagggcaagaagttcgccagataccagttcggcgacctgctgctgcacctggaaaa
gaagcacggcgaggactggggcaaagtgtacgatgaggcctgggagagaatcgacaagaaggtggaaggcctgagca
agcacattaagctggaagaggaaagaaggagcgaggacgcccaatctaaagccgctctgaccgattggctgagagcc
aaggccagctttgtgatcgagggcctgaaagaggccgacaaggacgagttctgcagatgcgagctgaagctgcagaa
gtggtacggcgatctgagaggcaagcccttcgccattgaggccgagaacagcatcctggacatcagcggcttcagca
agcagtacaactgcgccttcatttggcagaaagacggcgtcaagaaactgaacctgtacctgatcatcaattacttc
aaaggcggcaagctgcggttcaagaagatcaaacccgaggccttcgaggctaacagattctacaccgtgatcaacaa
aaagtccggcgagatcgtgcccatggaagtgaacttcaacttcgacgaccccaacctgattatcctgcctctggcct
tcggcaagagacagggcagagagttcatctggaacgatctgctgagcctggaaaccggctctctgaagctggccaat
ggcagagtgatcgagaaaaccctgtacaacaggagaaccagacaggacgagcctgctctgtttgtggccctgacctt
cgagagaagagaggtgctggacagcagcaacatcaagcccatgaacctgatcggcatcgaccggggcgagaatatcc
ctgctgtgatcgccctgacagaccctgaaggatgcccactgagcagattcaaggactccctgggcaaccctacacac
atcctgagaatcggcgagagctacaaagagaagcagaggacaatccaggccgccaaagaggtggaacagagaagagc
cggcggatactctaggaagtacgccagcaaggccaagaatctggccgacgacatggtccgaaacaccgccagagatc
tgctgtactacgccgtgacacaggacgccatgctgatcttcgagaatctgagcagaggcttcggccggcagggcaag
agaacctttatggccgagaggcagtacaccagaatggaagattggctcacagctaaactggcctacgagggactgcc
cagcaagacctacctgtccaaaacactggcccagtatacctccaagacctgcagcaattgcggcttcaccatcacca
gcgccgactacgacagagtgctggaaaagctcaagaaaaccgccaccggctggatgaccaccatcaacggcaaagag
ctgaaggttgagggccagatcacctactacaacaggtacaagaggcagaacgtcgtgaaggatctgagcgtggaact
ggacagactgagcgaagagagcgtgaacaacgacatcagcagctggacaaagggcagatcaggcgaggctctgagcc

tgctgaagaagaggtttagccacagacctgtgcaagagaagttcgtgtgcctgaactgcggcttcgagacacacgcc
gatgaacaggctgccctgaacattgccagaagctggctgttcctgagaagccaagagtacaagaagtaccagaccaa
caagaccaccggcaacaccgacaagagggcctttgtggaaacctggcagagcttctacagaaaaaagctgaaagaag
tctggaagcccgccgtgAATGCATTGCCAAAGAAGAAGCGGAAGGTCGGCAGTTACCCATACGATGTTCCAGATTAC
GCTTACCCATACGATGTTCCAGATTACGCTTACCCATACGATGTTCCAGATTACGCTTAAGAATTCCTAGAGCTCGC
TGATCAGCCTCGACTGTGCCTTCTAGTTGCCAGCCATCTGTTGTTTGCCCCTCCCCCGTGCCTTCCTTGACCCTGGA
AGGTGCCACTCCCACTGTCCTTTCCTAATAAAATGAGGAAATTGCATCGCATTGTCTGAGTAGGTGTCATTCTATTC
TGGGGGGTGGGGTGGGGCAGGACAGCAAGGGGGAGGATTGGGAAGAGAATAGCAGGCATGCTGGGGAGGTACCCTTT
TGCTGGCCTTTTGCTCACATGTGAGGGCCTATTTCCCATGATTCCTTCATATTTGCATATACGATACAAGGCTGTTA
GAGAGATAATTGGAATTAATTTGACTGTAAACACAAAGATATTAGTACAAAATACGTGACGTAGAAAGTAATAATTT
CTTGGGTAGTTTGCAGTTTTAAAATTATGTTTTAAAATGGACTATCATATGCTTACCGTAACTTGAAAGTATTTCGA
TTTCTTGGCTTTATATATCTTGTGGAAAGGACGAAACACCGTACTGGCGCTTTTATCTCATTACTTTGAGAGCCATC
ACCAGCGACTATGTCGTATGGGTAAAGCGCTTATTTATCGGAGAGAAATCCGATAAATAAGAAGCATCAAAGGGTCT
TCTCGAAGACATGCCGTTTGCGGCCGCAGGAACCCCTAGTGATGGAGTTGGCCACTCCCTCTCTGCGCGCTCGCTCG
CTCACTGAGGCCGGGCGACCAAAGGTCGCCCGACGCCCGGGCTTTGCCCGGGCGGCCTCAGTGAGCGAGCGAGCGCG
CAGCTGCCTGCAGGGGCGCCTGATGCGGTATTTTCTCCTTACGCATCTGTGCGGTATTTCACACCGCATACGTCAAA
GCAACCATAGTACGCGCCCTGTAGCGGCGCATTAAGCGCGGCGGGTGTGGTGGTTACGCGCAGCGTGACCGCTACAC
TTGCCAGCGCCTTAGCGCCCGCTCCTTTCGCTTTCTTCCCTTCCTTTCTCGCCACGTTCGCCGGCTTTCCCCGTCAA
GCTCTAAATCGGGGGCTCCCTTTAGGGTTCCGATTTAGTGCTTTACGGCACCTCGACCCCAAAAAACTTGATTTGGG
TGATGGTTCACGTAGTGGGCCATCGCCCTGATAGACGGTTTTTCGCCCTTTGACGTTGGAGTCCACGTTCTTTAATA
GTGGACTCTTGTTCCAAACTGGAACAACACTCAACTCTATCTCGGGCTATTCTTTTGATTTATAAGGGATTTTGCCG
ATTTCGGTCTATTGGTTAAAAAATGAGCTGATTTAACAAAAATTTAACGCGAATTTTAACAAAATATTAACGTTTAC
AATTTTATGGTGCACTCTCAGTACAATCTGCTCTGATGCCGCATAGTTAAGCCAGCCCCGACACCCGCCAACACCCG
CTGACGCGCCCTGACGGGCTTGTCTGCTCCCGGCATCCGCTTACAGACAAGCTGTGACCGTCTCCGGGAGCTGCATG
TGTCAGAGGTTTTCACCGTCATCACCGAAACGCGCGAGACGAAAGGGCCTCGTGATACGCCTATTTTTATAGGTTAA
TGTCATGATAATAATGGTTTCTTAGACGTCAGGTGGCACTTTTCGGGGAAATGTGCGCGGAACCCCTATTTGTTTAT
TTTTCTAAATACATTCAAATATGTATCCGCTCATGAGACAATAACCCTGATAAATGCTTCAATAATATTGAAAAAGG
AAGAGTATGAGTATTCAACATTTCCGTGTCGCCCTTATTCCCTTTTTTGCGGCATTTTGCCTTCCTGTTTTTGCTCA
CCCAGAAACGCTGGTGAAAGTAAAAGATGCTGAAGATCAGTTGGGTGCACGAGTGGGTTACATCGAACTGGATCTCA
ACAGCGGTAAGATCCTTGAGAGTTTTCGCCCCGAAGAACGTTTTCCAATGATGAGCACTTTTAAAGTTCTGCTATGT
GGCGCGGTATTATCCCGTATTGACGCCGGGCAAGAGCAACTCGGTCGCCGCATACACTATTCTCAGAATGACTTGGT
TGAGTACTCACCAGTCACAGAAAAGCATCTTACGGATGGCATGACAGTAAGAGAATTATGCAGTGCTGCCATAACCA
TGAGTGATAACACTGCGGCCAACTTACTTCTGACAACGATCGGAGGACCGAAGGAGCTAACCGCTTTTTTGCACAAC
ATGGGGGATCATGTAACTCGCCTTGATCGTTGGGAACCGGAGCTGAATGAAGCCATACCAAACGACGAGCGTGACAC
CACGATGCCTGTAGCAATGGCAACAACGTTGCGCAAACTATTAACTGGCGAACTACTTACTCTAGCTTCCCGGCAAC
AATTAATAGACTGGATGGAGGCGGATAAAGTTGCAGGACCACTTCTGCGCTCGGCCCTTCCGGCTGGCTGGTTTATT
GCTGATAAATCTGGAGCCGGTGAGCGTGGAAGCCGCGGTATCATTGCAGCACTGGGGCCAGATGGTAAGCCCTCCCG
TATCGTAGTTATCTACACGACGGGGAGTCAGGCAACTATGGATGAACGAAATAGACAGATCGCTGAGATAGGTGCCT
CACTGATTAAGCATTGGTAACTGTCAGACCAAGTTTACTCATATATACTTTAGATTGATTTAAAACTTCATTTTTAA
TTTAAAAGGATCTAGGTGAAGATCCTTTTTGATAATCTCATGACCAAAATCCCTTAACGTGAGTTTTCGTTCCACTG
AGCGTCAGACCCCG

**>CasX2_dual scaffold**

TAGAAAAGATCAAAGGATCTTCTTGAGATCCTTTTTTTCTGCGCGTAATCTGCTGCTTGCAAACAAAAAAACCACCG
CTACCAGCGGTGGTTTGTTTGCCGGATCAAGAGCTACCAACTCTTTTTCCGAAGGTAACTGGCTTCAGCAGAGCGCA
GATACCAAATACTGTTCTTCTAGTGTAGCCGTAGTTAGGCCACCACTTCAAGAACTCTGTAGCACCGCCTACATACC
TCGCTCTGCTAATCCTGTTACCAGTGGCTGCTGCCAGTGGCGATAAGTCGTGTCTTACCGGGTTGGACTCAAGACGA

TAGTTACCGGATAAGGCGCAGCGGTCGGGCTGAACGGGGGGTTCGTGCACACAGCCCAGCTTGGAGCGAACGACCTA
CACCGAACTGAGATACCTACAGCGTGAGCTATGAGAAAGCGCCACGCTTCCCGAAGGGAGAAAGGCGGACAGGTATC
CGGTAAGCGGCAGGGTCGGAACAGGAGAGCGCACGAGGGAGCTTCCAGGGGGAAACGCCTGGTATCTTTATAGTCCT
GTCGGGTTTCGCCACCTCTGACTTGAGCGTCGATTTTTGTGATGCTCGTCAGGGGGGCGGAGCCTATGGAAAAACGC
CAGCAACGCGGCCTTTTTACGGTTCCTGGCCTTTTGCTGGCCTTTTGCTCACATGTCCTGCAGGCAGCTGCGCGCTC
GCTCGCTCACTGAGGCCGCCCGGGCGTCGGGCGACCTTTGGTCGCCCGGCCTCAGTGAGCGAGCGAGCGCGCAGAGA
GGGAGTGGCCAACTCCATCACTAGGGGTTCCTGCGGCCTCTAGACTCGAGGCGTTGACATTGATTATTGACTAGTTA
TTAATAGTAATCAATTACGGGGTCATTAGTTCATAGCCCATATATGGAGTTCCGCGTTACATAACTTACGGTAAATG
GCCCGCCTGGCTGACCGCCCAACGACCCCCGCCCATTGACGTCAATAATGACGTATGTTCCCATAGTAACGCCAATA
GGGACTTTCCATTGACGTCAATGGGTGGAGTATTTACGGTAAACTGCCCACTTGGCAGTACATCAAGTGTATCATAT
GCCAAGTACGCCCCCTATTGACGTCAATGACGGTAAATGGCCCGCCTGGCATTATGCCCAGTACATGACCTTATGGG
ACTTTCCTACTTGGCAGTACATCTACGTATTAGTCATCGCTATTACCATGGTGATGCGGTTTTGGCAGTACATCAAT
GGGCGTGGATAGCGGTTTGACTCACGGGGATTTCCAAGTCTCCACCCCATTGACGTCAATGGGAGTTTGTTTTGGCA
CCAAAATCAACGGGACTTTCCAAAATGTCGTAACAACTCCGCCCCATTGACGCAAATGGGCGGTAGGCGTGTACGGT
GGGAGGTCTATATAAGCAGAGCTCTCTGGCTAACTACCGGTGCCACCATGTCCGGATCCCCTGCTGCCAAGAGGGTC
AAGTTGGACATGCAAGAGATCAAGAGAATCAACAAGATCAGAAGGAGACTGGTCAAGGAcagcaacacaaagaaggc
cggcaagacaggccccatgaaaaccctgctcgtcagagtgatgacccctgacctgagagagcggctggaaaacctga
gaaagaagcccgagaacatccctcagcctatcagcaacaccagcagggccaacctgaacaagctgctgaccgactac
accgagatgaagaaagccatcctgcacgtgtactgggaagagttccagaaagaccccgtgggcctgatgagcagagt
tgctcagcccgctcctaagaacatcgaccagagaaagctgatccccgtgaaggacggcaacgagagactgacctcta
gcggctttgcctgcagccagtgttgccagcctctgtacgtgtacaagctggaacaagtgaacgacaagggcaagccc
cacaccaactacttcggcagatgcaacgtgtccgagcacgagaggctgatcctgctgtctcctcacaagcccgaggc
caacgatgagctggtcacatacagcctgggcaagttcggacagagagccctggacttctacagcatccacgtgacca
gggagagcaatcaccctgtgaagcccctggaacagatcggcggcaatagctgtgcctctggacctgtgggaaaagcc
ctgagcgacgcctgtatgggagccgtggcatccttcctgaccaagtaccaggacatcatcctggaacaccagaaagt
gatcaagaagaacgagaaaagactggccaacctcaaggatatcgccagcgctaacggcctggcctttcctaagatca
ccctgcctccacagcctcacaccaaagagggcatcgaggcctacaacaacgtggtggcccagatcgtgatttgggtc
aacctgaatctgtggcagaagctgaagatcggcagggacgaagccaagccactgcagagactgaagggcttccctag
cttccctctggtggaaagacaggccaatgaagtggattggtgggacatggtctgcaacgtgaagaagctgatcaacg
agaagaaagaggatggcaaggttttctggcagaacctggccggctacaagagacaagaagccctgctgccttacctg
agcagcgaagaggaccggaagaagggcaagaagttcgccagataccagttcggcgacctgctgctgcacctggaaaa
gaagcacggcgaggactggggcaaagtgtacgatgaggcctgggagagaatcgacaagaaggtggaaggcctgagca
agcacattaagctggaagaggaaagaaggagcgaggacgcccaatctaaagccgctctgaccgattggctgagagcc
aaggccagctttgtgatcgagggcctgaaagaggccgacaaggacgagttctgcagatgcgagctgaagctgcagaa
gtggtacggcgatctgagaggcaagcccttcgccattgaggccgagaacagcatcctggacatcagcggcttcagca
agcagtacaactgcgccttcatttggcagaaagacggcgtcaagaaactgaacctgtacctgatcatcaattacttc
aaaggcggcaagctgcggttcaagaagatcaaacccgaggccttcgaggctaacagattctacaccgtgatcaacaa
aaagtccggcgagatcgtgcccatggaagtgaacttcaacttcgacgaccccaacctgattatcctgcctctggcct
tcggcaagagacagggcagagagttcatctggaacgatctgctgagcctggaaaccggctctctgaagctggccaat
ggcagagtgatcgagaaaaccctgtacaacaggagaaccagacaggacgagcctgctctgtttgtggccctgacctt
cgagagaagagaggtgctggacagcagcaacatcaagcccatgaacctgatcggcatcgaccggggcgagaatatcc
ctgctgtgatcgccctgacagaccctgaaggatgcccactgagcagattcaaggactccctgggcaaccctacacac
atcctgagaatcggcgagagctacaaagagaagcagaggacaatccaggccgccaaagaggtggaacagagaagagc
cggcggatactctaggaagtacgccagcaaggccaagaatctggccgacgacatggtccgaaacaccgccagagatc
tgctgtactacgccgtgacacaggacgccatgctgatcttcgagaatctgagcagaggcttcggccggcagggcaag
agaacctttatggccgagaggcagtacaccagaatggaagattggctcacagctaaactggcctacgagggactgcc
cagcaagacctacctgtccaaaacactggcccagtatacctccaagacctgcagcaattgcggcttcaccatcacca
gcgccgactacgacagagtgctggaaaagctcaagaaaaccgccaccggctggatgaccaccatcaacggcaaagag
ctgaaggttgagggccagatcacctactacaacaggtacaagaggcagaacgtcgtgaaggatctgagcgtggaact

ggacagactgagcgaagagagcgtgaacaacgacatcagcagctggacaaagggcagatcaggcgaggctctgagcc
tgctgaagaagaggtttagccacagacctgtgcaagagaagttcgtgtgcctgaactgcggcttcgagacacacgcc
gatgaacaggctgccctgaacattgccagaagctggctgttcctgagaagccaagagtacaagaagtaccagaccaa
caagaccaccggcaacaccgacaagagggcctttgtggaaacctggcagagcttctacagaaaaaagctgaaagaag
tctggaagcccgccgtgAATGCATTGCCAAAGAAGAAGCGGAAGGTCGGCAGTTACCCATACGATGTTCCAGATTAC
GCTTACCCATACGATGTTCCAGATTACGCTTACCCATACGATGTTCCAGATTACGCTTAAGAATTCCTAGAGCTCGC
TGATCAGCCTCGACTGTGCCTTCTAGTTGCCAGCCATCTGTTGTTTGCCCCTCCCCCGTGCCTTCCTTGACCCTGGA
AGGTGCCACTCCCACTGTCCTTTCCTAATAAAATGAGGAAATTGCATCGCATTGTCTGAGTAGGTGTCATTCTATTC
TGGGGGGTGGGGTGGGGCAGGACAGCAAGGGGGAGGATTGGGAAGAGAATAGCAGGCATGCTGGGGAGGTACCCTTT
TGCTGGCCTTTTGCTCACATGTGAGGGCCTATTTCCCATGATTCCTTCATATTTGCATATACGATACAAGGCTGTTA
GAGAGATAATTGGAATTAATTTGACTGTAAACACAAAGATATTAGTACAAAATACGTGACGTAGAAAGTAATAATTT
CTTGGGTAGTTTGCAGTTTTAAAATTATGTTTTAAAATGGACTATCATATGCTTACCGTAACTTGAAAGTATTTCGA
TTTCTTGGCTTTATATATCTTGTGGAAAGGACGAAACACCGTACTGGCGCTTTTATCTCATTACTTTGAGAGCCATC
ACCAGCGACTATGTCGTATGGGTAAAGCGCTTATTTATCGGAGAGAAATCCGATAAATAAGAAGCATCAAGTCTTCG
AGAAGACCTTTTTTGGTACGGCCGCCTAGTCTGGCCGCGAGGGCCTATTTCCCATGATTCCTTCATATTTGCATATA
CGATACAAGGCTGTTAGAGAGATAATTGGAATTAATTTGACTGTAAACACAAAGATATTAGTACAAAATACGTGACG
TAGAAAGTAATAATTTCTTGGGTAGTTTGCAGTTTTAAAATTATGTTTTAAAATGGACTATCATATGCTTACCGTAA
CTTGAAAGTATTTCGATTTCTTGGCTTTATATATCTTGTGGAAAGGACGAAACACCGTACTGGCGCTTTTATCTCAT
TACTTTGAGAGCCATCACCAGCGACTATGTCGTATGGGTAAAGCGCTTATTTATCGGAGAGAAATCCGATAAATAAG
AAGCATCAAAGGGTCTTCTCGAAGACATGCCGTTTGCGGCCGCAGGAACCCCTAGTGATGGAGTTGGCCACTCCCTC
TCTGCGCGCTCGCTCGCTCACTGAGGCCGGGCGACCAAAGGTCGCCCGACGCCCGGGCTTTGCCCGGGCGGCCTCAG
TGAGCGAGCGAGCGCGCAGCTGCCTGCAGGGGCGCCTGATGCGGTATTTTCTCCTTACGCATCTGTGCGGTATTTCA
CACCGCATACGTCAAAGCAACCATAGTACGCGCCCTGTAGCGGCGCATTAAGCGCGGCGGGTGTGGTGGTTACGCGC
AGCGTGACCGCTACACTTGCCAGCGCCTTAGCGCCCGCTCCTTTCGCTTTCTTCCCTTCCTTTCTCGCCACGTTCGC
CGGCTTTCCCCGTCAAGCTCTAAATCGGGGGCTCCCTTTAGGGTTCCGATTTAGTGCTTTACGGCACCTCGACCCCA
AAAAACTTGATTTGGGTGATGGTTCACGTAGTGGGCCATCGCCCTGATAGACGGTTTTTCGCCCTTTGACGTTGGAG
TCCACGTTCTTTAATAGTGGACTCTTGTTCCAAACTGGAACAACACTCAACTCTATCTCGGGCTATTCTTTTGATTT
ATAAGGGATTTTGCCGATTTCGGTCTATTGGTTAAAAAATGAGCTGATTTAACAAAAATTTAACGCGAATTTTAACA
AAATATTAACGTTTACAATTTTATGGTGCACTCTCAGTACAATCTGCTCTGATGCCGCATAGTTAAGCCAGCCCCGA
CACCCGCCAACACCCGCTGACGCGCCCTGACGGGCTTGTCTGCTCCCGGCATCCGCTTACAGACAAGCTGTGACCGT
CTCCGGGAGCTGCATGTGTCAGAGGTTTTCACCGTCATCACCGAAACGCGCGAGACGAAAGGGCCTCGTGATACGCC
TATTTTTATAGGTTAATGTCATGATAATAATGGTTTCTTAGACGTCAGGTGGCACTTTTCGGGGAAATGTGCGCGGA
ACCCCTATTTGTTTATTTTTCTAAATACATTCAAATATGTATCCGCTCATGAGACAATAACCCTGATAAATGCTTCA
ATAATATTGAAAAAGGAAGAGTATGAGTATTCAACATTTCCGTGTCGCCCTTATTCCCTTTTTTGCGGCATTTTGCC
TTCCTGTTTTTGCTCACCCAGAAACGCTGGTGAAAGTAAAAGATGCTGAAGATCAGTTGGGTGCACGAGTGGGTTAC
ATCGAACTGGATCTCAACAGCGGTAAGATCCTTGAGAGTTTTCGCCCCGAAGAACGTTTTCCAATGATGAGCACTTT
TAAAGTTCTGCTATGTGGCGCGGTATTATCCCGTATTGACGCCGGGCAAGAGCAACTCGGTCGCCGCATACACTATT
CTCAGAATGACTTGGTTGAGTACTCACCAGTCACAGAAAAGCATCTTACGGATGGCATGACAGTAAGAGAATTATGC
AGTGCTGCCATAACCATGAGTGATAACACTGCGGCCAACTTACTTCTGACAACGATCGGAGGACCGAAGGAGCTAAC
CGCTTTTTTGCACAACATGGGGGATCATGTAACTCGCCTTGATCGTTGGGAACCGGAGCTGAATGAAGCCATACCAA
ACGACGAGCGTGACACCACGATGCCTGTAGCAATGGCAACAACGTTGCGCAAACTATTAACTGGCGAACTACTTACT
CTAGCTTCCCGGCAACAATTAATAGACTGGATGGAGGCGGATAAAGTTGCAGGACCACTTCTGCGCTCGGCCCTTCC
GGCTGGCTGGTTTATTGCTGATAAATCTGGAGCCGGTGAGCGTGGAAGCCGCGGTATCATTGCAGCACTGGGGCCAG
ATGGTAAGCCCTCCCGTATCGTAGTTATCTACACGACGGGGAGTCAGGCAACTATGGATGAACGAAATAGACAGATC
GCTGAGATAGGTGCCTCACTGATTAAGCATTGGTAACTGTCAGACCAAGTTTACTCATATATACTTTAGATTGATTT
AAAACTTCATTTTTAATTTAAAAGGATCTAGGTGAAGATCCTTTTTGATAATCTCATGACCAAAATCCCTTAACGTG
AGTTTTCGTTCCACTGAGCGTCAGACCCCG

**>CasX2Max_CCR5_sg5**

TAGAAAAGATCAAAGGATCTTCTTGAGATCCTTTTTTTCTGCGCGTAATCTGCTGCTTGCAAACAAAAAAACCACCG
CTACCAGCGGTGGTTTGTTTGCCGGATCAAGAGCTACCAACTCTTTTTCCGAAGGTAACTGGCTTCAGCAGAGCGCA
GATACCAAATACTGTTCTTCTAGTGTAGCCGTAGTTAGGCCACCACTTCAAGAACTCTGTAGCACCGCCTACATACC
TCGCTCTGCTAATCCTGTTACCAGTGGCTGCTGCCAGTGGCGATAAGTCGTGTCTTACCGGGTTGGACTCAAGACGA
TAGTTACCGGATAAGGCGCAGCGGTCGGGCTGAACGGGGGGTTCGTGCACACAGCCCAGCTTGGAGCGAACGACCTA
CACCGAACTGAGATACCTACAGCGTGAGCTATGAGAAAGCGCCACGCTTCCCGAAGGGAGAAAGGCGGACAGGTATC
CGGTAAGCGGCAGGGTCGGAACAGGAGAGCGCACGAGGGAGCTTCCAGGGGGAAACGCCTGGTATCTTTATAGTCCT
GTCGGGTTTCGCCACCTCTGACTTGAGCGTCGATTTTTGTGATGCTCGTCAGGGGGGCGGAGCCTATGGAAAAACGC
CAGCAACGCGGCCTTTTTACGGTTCCTGGCCTTTTGCTGGCCTTTTGCTCACATGTCCTGCAGGCAGCTGCGCGCTC
GCTCGCTCACTGAGGCCGCCCGGGCGTCGGGCGACCTTTGGTCGCCCGGCCTCAGTGAGCGAGCGAGCGCGCAGAGA
GGGAGTGGCCAACTCCATCACTAGGGGTTCCTGCGGCCTCTAGACTCGAGGCGTTGACATTGATTATTGACTAGTTA
TTAATAGTAATCAATTACGGGGTCATTAGTTCATAGCCCATATATGGAGTTCCGCGTTACATAACTTACGGTAAATG
GCCCGCCTGGCTGACCGCCCAACGACCCCCGCCCATTGACGTCAATAATGACGTATGTTCCCATAGTAACGCCAATA
GGGACTTTCCATTGACGTCAATGGGTGGAGTATTTACGGTAAACTGCCCACTTGGCAGTACATCAAGTGTATCATAT
GCCAAGTACGCCCCCTATTGACGTCAATGACGGTAAATGGCCCGCCTGGCATTATGCCCAGTACATGACCTTATGGG
ACTTTCCTACTTGGCAGTACATCTACGTATTAGTCATCGCTATTACCATGGTGATGCGGTTTTGGCAGTACATCAAT
GGGCGTGGATAGCGGTTTGACTCACGGGGATTTCCAAGTCTCCACCCCATTGACGTCAATGGGAGTTTGTTTTGGCA
CCAAAATCAACGGGACTTTCCAAAATGTCGTAACAACTCCGCCCCATTGACGCAAATGGGCGGTAGGCGTGTACGGT
GGGAGGTCTATATAAGCAGAGCTCTCTGGCTAACTACCGGTTCTAGAGCGCTGCCACCATGTCCGGATCCCCTGCTG
CCAAGAGGGTCAAGTTGGACATGCAAGAGATCAAGAGAATCAACAAGATCAGAAGGAGACTGGTCAAGGACAGCAAC
ACAAAGAAGGCCGGCAAGCGCGGCCCCATGAAAACCCTGCTCGTCAGAGTGATGACCCCTGACCTGAGAGAGCGGCT
GGAAAACCTGAGAAAGAAGCCCGAGAACATCCCTCAGCCTATCAGCAACACCAGCAGGGCCAACCTGAACAAGCTGC
TGACCGACTACACCGAGATGAAGAAAGCCATCCTGCACGTGTACTGGGAAGAGTTCCAGAAAGACCCCGTGGGCCTG
ATGAGCAGAGTTGCTCAGCCCGCTCCTAAGAACATCGACCAGAGAAAGCTGATCCCCGTGAAGGACGGCAACGAGAG
ACTGACCTCTAGCGGCTTTGCCTGCAGCCAGTGTTGCCAGCCTCTGTACGTGTACAAGCTGGAACAAGTGAACGACA
AGGGCAAGCCCCACACCAACTACTTCGGCAGATGCAACGTGTCCGAGCACGAGAGGCTGATCCTGCTGTCTCCTCAC
AAGCCCGAGGCCAACGATGAGCTGGTCACATACAGCCTGGGCAAGTTCGGACAGAGAGCCCTGGACTTCTACAGCAT
CCACGTGACCAGGGAGAGCAATCACCCTGTGAAGCCCCTGGAACAGATCGGCGGCAATAGCTGTGCCTCTGGACCTG
TGGGAAAAGCCCTGAGCGACGCCTGTATGGGAGCCGTGGCATCCTTCCTGACCAAGTACCAGGACATCATCCTGGAA
CACCAGAAAGTGATCAAGAAGAACGAGAAAAGACTGGCCAACCTCAAGGATATCGCCAGCGCTAACGGCCTGGCCTT
TCCTAAGATCACCCTGCCTCCACAGCCTCACACCAAAGAGGGCATCGAGGCCTACAACAACGTGGTGGCCCAGATCG
TGATTTGGGTCAACCTGAATCTGTGGCAGAAGCTGAAGATCGGCAGGGACGAAGCCAAGCCACTGCAGAGACTGAAG
GGCTTCCCTAGCTTCCCTCTGGTGGAAAGACAGGCCAATGAAGTGGATTGGTGGGACATGGTCTGCAACGTGAAGAA
GCTGATCAACGAGAAGAAAGAGGATGGCAAGGTTTTCTGGCAGAACCTGGCCGGCTACAAGAGACAAGAAGCCCTGC
TGCCTTACCTGAGCAGCGAAGAGGACCGGAAGAAGGGCAAGAAGTTCGCCAGATACCAGTTCGGCGACCTGCTGCTG
CACCTGGAAAAGAAGCACGGCGAGGACTGGGGCAAAGTGTACGATGAGGCCTGGGAGAGAATCGACAAGAAGGTGGA
AGGCCTGAGCAAGCACATTAAGCTGGAAGAGGAAAGAAGGAGCGAGGACGCCCAATCTAAAGCCGCTCTGACCGATT
GGCTGAGAGCCAAGGCCAGCTTTGTGATCGAGGGCCTGAAAGAGGCCGACAAGGACGAGTTCTGCAGATGCGAGCTG
AAGCTGCAGAAGTGGTACGGCGATCTGAGAGGCAAGCCCTTCGCCATTGAGGCCGAGAACAGCATCCTGGACATCAG
CGGCTTCAGCAAGCAGTACAACTGCGCCTTCATTTGGCAGAAAGACGGCGTCAAGAAACTGAACCTGTACCTGATCA
TCAATTACTTCAAAGGCGGCAAGCTGCGGTTCAAGAAGATCAAACCCGAGGCCTTCGAGGCTAACAGATTCTACACC
GTGATCAACAAAAAGTCCGGCGAGATCGTGCCCATGGAAGTGAACTTCAACTTCGACGACCCCAACCTGATTATCCT
GCCTCTGGCCTTCGGCAAGAGACAGGGCAGAGAGTTCATCTGGAACGATCTGCTGAGCCTGGAAACCGGCTCTCTGC
GCCTGGCCAATGGCAGAGTGATCGAGAAAACCCTGTACAACAGGAGAACCAGACAGGACGAGCCTGCTCTGTTTGTG
GCCCTGACCTTCGAGAGAAGAGAGGTGCTGGACAGCAGCAACATCAAGCCCATGAACCTGATCGGCATCGACCGGGG
CGAGAATATCCCTGCTGTGATCGCCCTGACAGACCCTGAAGGATGCCCACTGAGCAGATTCAAGGACTCCCTGGGCA

ACCCTACACACATCCTGAGAATCGGCGAGAGCTACAAAGAGAAGCAGAGGACAATCCAGGCCGCCAAAGAGGTGGAA
CAGAGAAGAGCCGGCGGATACTCTAGGAAGTACGCCAGCAAGGCCAAGAATCTGGCCGACGACATGGTCCGAAACAC
CGCCAGAGATCTGCTGTACTACGCCGTGACACAGGACGCCATGCTGATCTTCGAGAATCTGAGCAGAGGCTTCGGCC
GGCAGGGCAAGAGAACCTTTATGGCCGAGAGGCAGTACACCAGAATGGAAGATTGGCTCACAGCTAAACTGGCCTAC
GAGGGACTGCCCAGCAAGACCTACCTGTCCAAAACACTGGCCCAGTATACCTCCCGCACCTGCAGCaattgcggctt
caccatcaccagcgccgactacgacagagtgctggaaaagctcaagaaaaccgccaccggctggatgaccaccatca
acggcaaagagctgaaggttgagggccagatcacctactacaacaggtacaagaggcagaacgtcgtgaaggatctg
agcgtggaactggacagactgagcgaagagagcgtgaacaacgacatcagcagctggacaaagggcagatcaggcga
ggctctgagcctgctgaagaagaggtttagccacagacctgtgcaagagaagttcgtgtgcctgaactgcggcttcg
agacacacgccgatgaacaggctgccctgaacattgccagaagctggctgttcctgagaagccaagagtacaagaag
taccagaccaacaagaccaccggcaacaccgacaagagggcctttgtggaaacctggcagagcttctacagaaaaaa
gctgaaagaagtctggaagcccgccgtgAATGCATTGCCAAAGAAGAAGCGGAAGGTCGGCAGTTACCCATACGATG
TTCCAGATTACGCTTACCCATACGATGTTCCAGATTACGCTTACCCATACGATGTTCCAGATTACGCTTAAGAATTC
CTAGAGCTCGCTGATCAGCCTCGACTGTGCCTTCTAGTTGCCAGCCATCTGTTGTTTGCCCCTCCCCCGTGCCTTCC
TTGACCCTGGAAGGTGCCACTCCCACTGTCCTTTCCTAATAAAATGAGGAAATTGCATCGCATTGTCTGAGTAGGTG
TCATTCTATTCTGGGGGGTGGGGTGGGGCAGGACAGCAAGGGGGAGGATTGGGAAGAGAATAGCAGGCATGCTGGGG
AGGTACCCTTTTGCTGGCCTTTTGCTCACATGTGAGGGCCTATTTCCCATGATTCCTTCATATTTGCATATACGATA
CAAGGCTGTTAGAGAGATAATTGGAATTAATTTGACTGTAAACACAAAGATATTAGTACAAAATACGTGACGTAGAA
AGTAATAATTTCTTGGGTAGTTTGCAGTTTTAAAATTATGTTTTAAAATGGACTATCATATGCTTACCGTAACTTGA
AAGTATTTCGATTTCTTGGCTTTATATATCTTGTGGAAAGGACGAAACACCGTACTGGCGCTTTTATCTCATTACTT
TGAGAGCCATCACCAGCGACTATGTCGTATGGGTAAAGCGCTTATTTATCGGAGAGAAATCCGATAAATAAGAAGCA
TCAAAGAAAGTCCCACTGGGCGGCAGCATTTTTTGCCGTTTGCGGCCGCAGGAACCCCTAGTGATGGAGTTGGCCAC
TCCCTCTCTGCGCGCTCGCTCGCTCACTGAGGCCGGGCGACCAAAGGTCGCCCGACGCCCGGGCTTTGCCCGGGCGG
CCTCAGTGAGCGAGCGAGCGCGCAGCTGCCTGCAGGGGCGCCTGATGCGGTATTTTCTCCTTACGCATCTGTGCGGT
ATTTCACACCGCATACGTCAAAGCAACCATAGTACGCGCCCTGTAGCGGCGCATTAAGCGCGGCGGGTGTGGTGGTT
ACGCGCAGCGTGACCGCTACACTTGCCAGCGCCTTAGCGCCCGCTCCTTTCGCTTTCTTCCCTTCCTTTCTCGCCAC
GTTCGCCGGCTTTCCCCGTCAAGCTCTAAATCGGGGGCTCCCTTTAGGGTTCCGATTTAGTGCTTTACGGCACCTCG
ACCCCAAAAAACTTGATTTGGGTGATGGTTCACGTAGTGGGCCATCGCCCTGATAGACGGTTTTTCGCCCTTTGACG
TTGGAGTCCACGTTCTTTAATAGTGGACTCTTGTTCCAAACTGGAACAACACTCAACTCTATCTCGGGCTATTCTTT
TGATTTATAAGGGATTTTGCCGATTTCGGTCTATTGGTTAAAAAATGAGCTGATTTAACAAAAATTTAACGCGAATT
TTAACAAAATATTAACGTTTACAATTTTATGGTGCACTCTCAGTACAATCTGCTCTGATGCCGCATAGTTAAGCCAG
CCCCGACACCCGCCAACACCCGCTGACGCGCCCTGACGGGCTTGTCTGCTCCCGGCATCCGCTTACAGACAAGCTGT
GACCGTCTCCGGGAGCTGCATGTGTCAGAGGTTTTCACCGTCATCACCGAAACGCGCGAGACGAAAGGGCCTCGTGA
TACGCCTATTTTTATAGGTTAATGTCATGATAATAATGGTTTCTTAGACGTCAGGTGGCACTTTTCGGGGAAATGTG
CGCGGAACCCCTATTTGTTTATTTTTCTAAATACATTCAAATATGTATCCGCTCATGAGACAATAACCCTGATAAAT
GCTTCAATAATATTGAAAAAGGAAGAGTATGAGTATTCAACATTTCCGTGTCGCCCTTATTCCCTTTTTTGCGGCAT
TTTGCCTTCCTGTTTTTGCTCACCCAGAAACGCTGGTGAAAGTAAAAGATGCTGAAGATCAGTTGGGTGCACGAGTG
GGTTACATCGAACTGGATCTCAACAGCGGTAAGATCCTTGAGAGTTTTCGCCCCGAAGAACGTTTTCCAATGATGAG
CACTTTTAAAGTTCTGCTATGTGGCGCGGTATTATCCCGTATTGACGCCGGGCAAGAGCAACTCGGTCGCCGCATAC
ACTATTCTCAGAATGACTTGGTTGAGTACTCACCAGTCACAGAAAAGCATCTTACGGATGGCATGACAGTAAGAGAA
TTATGCAGTGCTGCCATAACCATGAGTGATAACACTGCGGCCAACTTACTTCTGACAACGATCGGAGGACCGAAGGA
GCTAACCGCTTTTTTGCACAACATGGGGGATCATGTAACTCGCCTTGATCGTTGGGAACCGGAGCTGAATGAAGCCA
TACCAAACGACGAGCGTGACACCACGATGCCTGTAGCAATGGCAACAACGTTGCGCAAACTATTAACTGGCGAACTA
CTTACTCTAGCTTCCCGGCAACAATTAATAGACTGGATGGAGGCGGATAAAGTTGCAGGACCACTTCTGCGCTCGGC
CCTTCCGGCTGGCTGGTTTATTGCTGATAAATCTGGAGCCGGTGAGCGTGGAAGCCGCGGTATCATTGCAGCACTGG
GGCCAGATGGTAAGCCCTCCCGTATCGTAGTTATCTACACGACGGGGAGTCAGGCAACTATGGATGAACGAAATAGA
CAGATCGCTGAGATAGGTGCCTCACTGATTAAGCATTGGTAACTGTCAGACCAAGTTTACTCATATATACTTTAGAT
TGATTTAAAACTTCATTTTTAATTTAAAAGGATCTAGGTGAAGATCCTTTTTGATAATCTCATGACCAAAATCCCTT
AACGTGAGTTTTCGTTCCACTGAGCGTCAGACCCCG

**>CasX2Max_CCR5_sg10**

TAGAAAAGATCAAAGGATCTTCTTGAGATCCTTTTTTTCTGCGCGTAATCTGCTGCTTGCAAACAAAAAAACCACCG
CTACCAGCGGTGGTTTGTTTGCCGGATCAAGAGCTACCAACTCTTTTTCCGAAGGTAACTGGCTTCAGCAGAGCGCA
GATACCAAATACTGTTCTTCTAGTGTAGCCGTAGTTAGGCCACCACTTCAAGAACTCTGTAGCACCGCCTACATACC
TCGCTCTGCTAATCCTGTTACCAGTGGCTGCTGCCAGTGGCGATAAGTCGTGTCTTACCGGGTTGGACTCAAGACGA
TAGTTACCGGATAAGGCGCAGCGGTCGGGCTGAACGGGGGGTTCGTGCACACAGCCCAGCTTGGAGCGAACGACCTA
CACCGAACTGAGATACCTACAGCGTGAGCTATGAGAAAGCGCCACGCTTCCCGAAGGGAGAAAGGCGGACAGGTATC
CGGTAAGCGGCAGGGTCGGAACAGGAGAGCGCACGAGGGAGCTTCCAGGGGGAAACGCCTGGTATCTTTATAGTCCT
GTCGGGTTTCGCCACCTCTGACTTGAGCGTCGATTTTTGTGATGCTCGTCAGGGGGGCGGAGCCTATGGAAAAACGC
CAGCAACGCGGCCTTTTTACGGTTCCTGGCCTTTTGCTGGCCTTTTGCTCACATGTCCTGCAGGCAGCTGCGCGCTC
GCTCGCTCACTGAGGCCGCCCGGGCGTCGGGCGACCTTTGGTCGCCCGGCCTCAGTGAGCGAGCGAGCGCGCAGAGA
GGGAGTGGCCAACTCCATCACTAGGGGTTCCTGCGGCCTCTAGACTCGAGGCGTTGACATTGATTATTGACTAGTTA
TTAATAGTAATCAATTACGGGGTCATTAGTTCATAGCCCATATATGGAGTTCCGCGTTACATAACTTACGGTAAATG
GCCCGCCTGGCTGACCGCCCAACGACCCCCGCCCATTGACGTCAATAATGACGTATGTTCCCATAGTAACGCCAATA
GGGACTTTCCATTGACGTCAATGGGTGGAGTATTTACGGTAAACTGCCCACTTGGCAGTACATCAAGTGTATCATAT
GCCAAGTACGCCCCCTATTGACGTCAATGACGGTAAATGGCCCGCCTGGCATTATGCCCAGTACATGACCTTATGGG
ACTTTCCTACTTGGCAGTACATCTACGTATTAGTCATCGCTATTACCATGGTGATGCGGTTTTGGCAGTACATCAAT
GGGCGTGGATAGCGGTTTGACTCACGGGGATTTCCAAGTCTCCACCCCATTGACGTCAATGGGAGTTTGTTTTGGCA
CCAAAATCAACGGGACTTTCCAAAATGTCGTAACAACTCCGCCCCATTGACGCAAATGGGCGGTAGGCGTGTACGGT
GGGAGGTCTATATAAGCAGAGCTCTCTGGCTAACTACCGGTTCTAGAGCGCTGCCACCATGTCCGGATCCCCTGCTG
CCAAGAGGGTCAAGTTGGACATGCAAGAGATCAAGAGAATCAACAAGATCAGAAGGAGACTGGTCAAGGACAGCAAC
ACAAAGAAGGCCGGCAAGCGCGGCCCCATGAAAACCCTGCTCGTCAGAGTGATGACCCCTGACCTGAGAGAGCGGCT
GGAAAACCTGAGAAAGAAGCCCGAGAACATCCCTCAGCCTATCAGCAACACCAGCAGGGCCAACCTGAACAAGCTGC
TGACCGACTACACCGAGATGAAGAAAGCCATCCTGCACGTGTACTGGGAAGAGTTCCAGAAAGACCCCGTGGGCCTG
ATGAGCAGAGTTGCTCAGCCCGCTCCTAAGAACATCGACCAGAGAAAGCTGATCCCCGTGAAGGACGGCAACGAGAG
ACTGACCTCTAGCGGCTTTGCCTGCAGCCAGTGTTGCCAGCCTCTGTACGTGTACAAGCTGGAACAAGTGAACGACA
AGGGCAAGCCCCACACCAACTACTTCGGCAGATGCAACGTGTCCGAGCACGAGAGGCTGATCCTGCTGTCTCCTCAC
AAGCCCGAGGCCAACGATGAGCTGGTCACATACAGCCTGGGCAAGTTCGGACAGAGAGCCCTGGACTTCTACAGCAT
CCACGTGACCAGGGAGAGCAATCACCCTGTGAAGCCCCTGGAACAGATCGGCGGCAATAGCTGTGCCTCTGGACCTG
TGGGAAAAGCCCTGAGCGACGCCTGTATGGGAGCCGTGGCATCCTTCCTGACCAAGTACCAGGACATCATCCTGGAA
CACCAGAAAGTGATCAAGAAGAACGAGAAAAGACTGGCCAACCTCAAGGATATCGCCAGCGCTAACGGCCTGGCCTT
TCCTAAGATCACCCTGCCTCCACAGCCTCACACCAAAGAGGGCATCGAGGCCTACAACAACGTGGTGGCCCAGATCG
TGATTTGGGTCAACCTGAATCTGTGGCAGAAGCTGAAGATCGGCAGGGACGAAGCCAAGCCACTGCAGAGACTGAAG
GGCTTCCCTAGCTTCCCTCTGGTGGAAAGACAGGCCAATGAAGTGGATTGGTGGGACATGGTCTGCAACGTGAAGAA
GCTGATCAACGAGAAGAAAGAGGATGGCAAGGTTTTCTGGCAGAACCTGGCCGGCTACAAGAGACAAGAAGCCCTGC
TGCCTTACCTGAGCAGCGAAGAGGACCGGAAGAAGGGCAAGAAGTTCGCCAGATACCAGTTCGGCGACCTGCTGCTG
CACCTGGAAAAGAAGCACGGCGAGGACTGGGGCAAAGTGTACGATGAGGCCTGGGAGAGAATCGACAAGAAGGTGGA
AGGCCTGAGCAAGCACATTAAGCTGGAAGAGGAAAGAAGGAGCGAGGACGCCCAATCTAAAGCCGCTCTGACCGATT
GGCTGAGAGCCAAGGCCAGCTTTGTGATCGAGGGCCTGAAAGAGGCCGACAAGGACGAGTTCTGCAGATGCGAGCTG
AAGCTGCAGAAGTGGTACGGCGATCTGAGAGGCAAGCCCTTCGCCATTGAGGCCGAGAACAGCATCCTGGACATCAG
CGGCTTCAGCAAGCAGTACAACTGCGCCTTCATTTGGCAGAAAGACGGCGTCAAGAAACTGAACCTGTACCTGATCA
TCAATTACTTCAAAGGCGGCAAGCTGCGGTTCAAGAAGATCAAACCCGAGGCCTTCGAGGCTAACAGATTCTACACC
GTGATCAACAAAAAGTCCGGCGAGATCGTGCCCATGGAAGTGAACTTCAACTTCGACGACCCCAACCTGATTATCCT
GCCTCTGGCCTTCGGCAAGAGACAGGGCAGAGAGTTCATCTGGAACGATCTGCTGAGCCTGGAAACCGGCTCTCTGC
GCCTGGCCAATGGCAGAGTGATCGAGAAAACCCTGTACAACAGGAGAACCAGACAGGACGAGCCTGCTCTGTTTGTG
GCCCTGACCTTCGAGAGAAGAGAGGTGCTGGACAGCAGCAACATCAAGCCCATGAACCTGATCGGCATCGACCGGGG
CGAGAATATCCCTGCTGTGATCGCCCTGACAGACCCTGAAGGATGCCCACTGAGCAGATTCAAGGACTCCCTGGGCA

ACCCTACACACATCCTGAGAATCGGCGAGAGCTACAAAGAGAAGCAGAGGACAATCCAGGCCGCCAAAGAGGTGGAA
CAGAGAAGAGCCGGCGGATACTCTAGGAAGTACGCCAGCAAGGCCAAGAATCTGGCCGACGACATGGTCCGAAACAC
CGCCAGAGATCTGCTGTACTACGCCGTGACACAGGACGCCATGCTGATCTTCGAGAATCTGAGCAGAGGCTTCGGCC
GGCAGGGCAAGAGAACCTTTATGGCCGAGAGGCAGTACACCAGAATGGAAGATTGGCTCACAGCTAAACTGGCCTAC
GAGGGACTGCCCAGCAAGACCTACCTGTCCAAAACACTGGCCCAGTATACCTCCCGCACCTGCAGCaattgcggctt
caccatcaccagcgccgactacgacagagtgctggaaaagctcaagaaaaccgccaccggctggatgaccaccatca
acggcaaagagctgaaggttgagggccagatcacctactacaacaggtacaagaggcagaacgtcgtgaaggatctg
agcgtggaactggacagactgagcgaagagagcgtgaacaacgacatcagcagctggacaaagggcagatcaggcga
ggctctgagcctgctgaagaagaggtttagccacagacctgtgcaagagaagttcgtgtgcctgaactgcggcttcg
agacacacgccgatgaacaggctgccctgaacattgccagaagctggctgttcctgagaagccaagagtacaagaag
taccagaccaacaagaccaccggcaacaccgacaagagggcctttgtggaaacctggcagagcttctacagaaaaaa
gctgaaagaagtctggaagcccgccgtgAATGCATTGCCAAAGAAGAAGCGGAAGGTCGGCAGTTACCCATACGATG
TTCCAGATTACGCTTACCCATACGATGTTCCAGATTACGCTTACCCATACGATGTTCCAGATTACGCTTAAGAATTC
CTAGAGCTCGCTGATCAGCCTCGACTGTGCCTTCTAGTTGCCAGCCATCTGTTGTTTGCCCCTCCCCCGTGCCTTCC
TTGACCCTGGAAGGTGCCACTCCCACTGTCCTTTCCTAATAAAATGAGGAAATTGCATCGCATTGTCTGAGTAGGTG
TCATTCTATTCTGGGGGGTGGGGTGGGGCAGGACAGCAAGGGGGAGGATTGGGAAGAGAATAGCAGGCATGCTGGGG
AGGTACCCTTTTGCTGGCCTTTTGCTCACATGTGAGGGCCTATTTCCCATGATTCCTTCATATTTGCATATACGATA
CAAGGCTGTTAGAGAGATAATTGGAATTAATTTGACTGTAAACACAAAGATATTAGTACAAAATACGTGACGTAGAA
AGTAATAATTTCTTGGGTAGTTTGCAGTTTTAAAATTATGTTTTAAAATGGACTATCATATGCTTACCGTAACTTGA
AAGTATTTCGATTTCTTGGCTTTATATATCTTGTGGAAAGGACGAAACACCGTACTGGCGCTTTTATCTCATTACTT
TGAGAGCCATCACCAGCGACTATGTCGTATGGGTAAAGCGCTTATTTATCGGAGAGAAATCCGATAAATAAGAAGCA
TCAAAGTGCTCCCCAGTGGATCGGGTTTTTTGCCGTTTGCGGCCGCAGGAACCCCTAGTGATGGAGTTGGCCACTCC
CTCTCTGCGCGCTCGCTCGCTCACTGAGGCCGGGCGACCAAAGGTCGCCCGACGCCCGGGCTTTGCCCGGGCGGCCT
CAGTGAGCGAGCGAGCGCGCAGCTGCCTGCAGGGGCGCCTGATGCGGTATTTTCTCCTTACGCATCTGTGCGGTATT
TCACACCGCATACGTCAAAGCAACCATAGTACGCGCCCTGTAGCGGCGCATTAAGCGCGGCGGGTGTGGTGGTTACG
CGCAGCGTGACCGCTACACTTGCCAGCGCCTTAGCGCCCGCTCCTTTCGCTTTCTTCCCTTCCTTTCTCGCCACGTT
CGCCGGCTTTCCCCGTCAAGCTCTAAATCGGGGGCTCCCTTTAGGGTTCCGATTTAGTGCTTTACGGCACCTCGACC
CCAAAAAACTTGATTTGGGTGATGGTTCACGTAGTGGGCCATCGCCCTGATAGACGGTTTTTCGCCCTTTGACGTTG
GAGTCCACGTTCTTTAATAGTGGACTCTTGTTCCAAACTGGAACAACACTCAACTCTATCTCGGGCTATTCTTTTGA
TTTATAAGGGATTTTGCCGATTTCGGTCTATTGGTTAAAAAATGAGCTGATTTAACAAAAATTTAACGCGAATTTTA
ACAAAATATTAACGTTTACAATTTTATGGTGCACTCTCAGTACAATCTGCTCTGATGCCGCATAGTTAAGCCAGCCC
CGACACCCGCCAACACCCGCTGACGCGCCCTGACGGGCTTGTCTGCTCCCGGCATCCGCTTACAGACAAGCTGTGAC
CGTCTCCGGGAGCTGCATGTGTCAGAGGTTTTCACCGTCATCACCGAAACGCGCGAGACGAAAGGGCCTCGTGATAC
GCCTATTTTTATAGGTTAATGTCATGATAATAATGGTTTCTTAGACGTCAGGTGGCACTTTTCGGGGAAATGTGCGC
GGAACCCCTATTTGTTTATTTTTCTAAATACATTCAAATATGTATCCGCTCATGAGACAATAACCCTGATAAATGCT
TCAATAATATTGAAAAAGGAAGAGTATGAGTATTCAACATTTCCGTGTCGCCCTTATTCCCTTTTTTGCGGCATTTT
GCCTTCCTGTTTTTGCTCACCCAGAAACGCTGGTGAAAGTAAAAGATGCTGAAGATCAGTTGGGTGCACGAGTGGGT
TACATCGAACTGGATCTCAACAGCGGTAAGATCCTTGAGAGTTTTCGCCCCGAAGAACGTTTTCCAATGATGAGCAC
TTTTAAAGTTCTGCTATGTGGCGCGGTATTATCCCGTATTGACGCCGGGCAAGAGCAACTCGGTCGCCGCATACACT
ATTCTCAGAATGACTTGGTTGAGTACTCACCAGTCACAGAAAAGCATCTTACGGATGGCATGACAGTAAGAGAATTA
TGCAGTGCTGCCATAACCATGAGTGATAACACTGCGGCCAACTTACTTCTGACAACGATCGGAGGACCGAAGGAGCT
AACCGCTTTTTTGCACAACATGGGGGATCATGTAACTCGCCTTGATCGTTGGGAACCGGAGCTGAATGAAGCCATAC
CAAACGACGAGCGTGACACCACGATGCCTGTAGCAATGGCAACAACGTTGCGCAAACTATTAACTGGCGAACTACTT
ACTCTAGCTTCCCGGCAACAATTAATAGACTGGATGGAGGCGGATAAAGTTGCAGGACCACTTCTGCGCTCGGCCCT
TCCGGCTGGCTGGTTTATTGCTGATAAATCTGGAGCCGGTGAGCGTGGAAGCCGCGGTATCATTGCAGCACTGGGGC
CAGATGGTAAGCCCTCCCGTATCGTAGTTATCTACACGACGGGGAGTCAGGCAACTATGGATGAACGAAATAGACAG
ATCGCTGAGATAGGTGCCTCACTGATTAAGCATTGGTAACTGTCAGACCAAGTTTACTCATATATACTTTAGATTGA
TTTAAAACTTCATTTTTAATTTAAAAGGATCTAGGTGAAGATCCTTTTTGATAATCTCATGACCAAAATCCCTTAAC
GTGAGTTTTCGTTCCACTGAGCGTCAGACCCCG

**>CasX2Max_single scaffold**

TAGAAAAGATCAAAGGATCTTCTTGAGATCCTTTTTTTCTGCGCGTAATCTGCTGCTTGCAAACAAAAAAACCACCG
CTACCAGCGGTGGTTTGTTTGCCGGATCAAGAGCTACCAACTCTTTTTCCGAAGGTAACTGGCTTCAGCAGAGCGCA
GATACCAAATACTGTTCTTCTAGTGTAGCCGTAGTTAGGCCACCACTTCAAGAACTCTGTAGCACCGCCTACATACC
TCGCTCTGCTAATCCTGTTACCAGTGGCTGCTGCCAGTGGCGATAAGTCGTGTCTTACCGGGTTGGACTCAAGACGA
TAGTTACCGGATAAGGCGCAGCGGTCGGGCTGAACGGGGGGTTCGTGCACACAGCCCAGCTTGGAGCGAACGACCTA
CACCGAACTGAGATACCTACAGCGTGAGCTATGAGAAAGCGCCACGCTTCCCGAAGGGAGAAAGGCGGACAGGTATC
CGGTAAGCGGCAGGGTCGGAACAGGAGAGCGCACGAGGGAGCTTCCAGGGGGAAACGCCTGGTATCTTTATAGTCCT
GTCGGGTTTCGCCACCTCTGACTTGAGCGTCGATTTTTGTGATGCTCGTCAGGGGGGCGGAGCCTATGGAAAAACGC
CAGCAACGCGGCCTTTTTACGGTTCCTGGCCTTTTGCTGGCCTTTTGCTCACATGTCCTGCAGGCAGCTGCGCGCTC
GCTCGCTCACTGAGGCCGCCCGGGCGTCGGGCGACCTTTGGTCGCCCGGCCTCAGTGAGCGAGCGAGCGCGCAGAGA
GGGAGTGGCCAACTCCATCACTAGGGGTTCCTGCGGCCTCTAGACTCGAGGCGTTGACATTGATTATTGACTAGTTA
TTAATAGTAATCAATTACGGGGTCATTAGTTCATAGCCCATATATGGAGTTCCGCGTTACATAACTTACGGTAAATG
GCCCGCCTGGCTGACCGCCCAACGACCCCCGCCCATTGACGTCAATAATGACGTATGTTCCCATAGTAACGCCAATA
GGGACTTTCCATTGACGTCAATGGGTGGAGTATTTACGGTAAACTGCCCACTTGGCAGTACATCAAGTGTATCATAT
GCCAAGTACGCCCCCTATTGACGTCAATGACGGTAAATGGCCCGCCTGGCATTATGCCCAGTACATGACCTTATGGG
ACTTTCCTACTTGGCAGTACATCTACGTATTAGTCATCGCTATTACCATGGTGATGCGGTTTTGGCAGTACATCAAT
GGGCGTGGATAGCGGTTTGACTCACGGGGATTTCCAAGTCTCCACCCCATTGACGTCAATGGGAGTTTGTTTTGGCA
CCAAAATCAACGGGACTTTCCAAAATGTCGTAACAACTCCGCCCCATTGACGCAAATGGGCGGTAGGCGTGTACGGT
GGGAGGTCTATATAAGCAGAGCTCTCTGGCTAACTACCGGTTCTAGAGCGCTGCCACCATGTCCGGATCCCCTGCTG
CCAAGAGGGTCAAGTTGGACATGCAAGAGATCAAGAGAATCAACAAGATCAGAAGGAGACTGGTCAAGGACAGCAAC
ACAAAGAAGGCCGGCAAGCGCGGCCCCATGAAAACCCTGCTCGTCAGAGTGATGACCCCTGACCTGAGAGAGCGGCT
GGAAAACCTGAGAAAGAAGCCCGAGAACATCCCTCAGCCTATCAGCAACACCAGCAGGGCCAACCTGAACAAGCTGC
TGACCGACTACACCGAGATGAAGAAAGCCATCCTGCACGTGTACTGGGAAGAGTTCCAGAAAGACCCCGTGGGCCTG
ATGAGCAGAGTTGCTCAGCCCGCTCCTAAGAACATCGACCAGAGAAAGCTGATCCCCGTGAAGGACGGCAACGAGAG
ACTGACCTCTAGCGGCTTTGCCTGCAGCCAGTGTTGCCAGCCTCTGTACGTGTACAAGCTGGAACAAGTGAACGACA
AGGGCAAGCCCCACACCAACTACTTCGGCAGATGCAACGTGTCCGAGCACGAGAGGCTGATCCTGCTGTCTCCTCAC
AAGCCCGAGGCCAACGATGAGCTGGTCACATACAGCCTGGGCAAGTTCGGACAGAGAGCCCTGGACTTCTACAGCAT
CCACGTGACCAGGGAGAGCAATCACCCTGTGAAGCCCCTGGAACAGATCGGCGGCAATAGCTGTGCCTCTGGACCTG
TGGGAAAAGCCCTGAGCGACGCCTGTATGGGAGCCGTGGCATCCTTCCTGACCAAGTACCAGGACATCATCCTGGAA
CACCAGAAAGTGATCAAGAAGAACGAGAAAAGACTGGCCAACCTCAAGGATATCGCCAGCGCTAACGGCCTGGCCTT
TCCTAAGATCACCCTGCCTCCACAGCCTCACACCAAAGAGGGCATCGAGGCCTACAACAACGTGGTGGCCCAGATCG
TGATTTGGGTCAACCTGAATCTGTGGCAGAAGCTGAAGATCGGCAGGGACGAAGCCAAGCCACTGCAGAGACTGAAG
GGCTTCCCTAGCTTCCCTCTGGTGGAAAGACAGGCCAATGAAGTGGATTGGTGGGACATGGTCTGCAACGTGAAGAA
GCTGATCAACGAGAAGAAAGAGGATGGCAAGGTTTTCTGGCAGAACCTGGCCGGCTACAAGAGACAAGAAGCCCTGC
TGCCTTACCTGAGCAGCGAAGAGGACCGGAAGAAGGGCAAGAAGTTCGCCAGATACCAGTTCGGCGACCTGCTGCTG
CACCTGGAAAAGAAGCACGGCGAGGACTGGGGCAAAGTGTACGATGAGGCCTGGGAGAGAATCGACAAGAAGGTGGA
AGGCCTGAGCAAGCACATTAAGCTGGAAGAGGAAAGAAGGAGCGAGGACGCCCAATCTAAAGCCGCTCTGACCGATT
GGCTGAGAGCCAAGGCCAGCTTTGTGATCGAGGGCCTGAAAGAGGCCGACAAGGACGAGTTCTGCAGATGCGAGCTG
AAGCTGCAGAAGTGGTACGGCGATCTGAGAGGCAAGCCCTTCGCCATTGAGGCCGAGAACAGCATCCTGGACATCAG
CGGCTTCAGCAAGCAGTACAACTGCGCCTTCATTTGGCAGAAAGACGGCGTCAAGAAACTGAACCTGTACCTGATCA
TCAATTACTTCAAAGGCGGCAAGCTGCGGTTCAAGAAGATCAAACCCGAGGCCTTCGAGGCTAACAGATTCTACACC
GTGATCAACAAAAAGTCCGGCGAGATCGTGCCCATGGAAGTGAACTTCAACTTCGACGACCCCAACCTGATTATCCT
GCCTCTGGCCTTCGGCAAGAGACAGGGCAGAGAGTTCATCTGGAACGATCTGCTGAGCCTGGAAACCGGCTCTCTGC
GCCTGGCCAATGGCAGAGTGATCGAGAAAACCCTGTACAACAGGAGAACCAGACAGGACGAGCCTGCTCTGTTTGTG
GCCCTGACCTTCGAGAGAAGAGAGGTGCTGGACAGCAGCAACATCAAGCCCATGAACCTGATCGGCATCGACCGGGG
CGAGAATATCCCTGCTGTGATCGCCCTGACAGACCCTGAAGGATGCCCACTGAGCAGATTCAAGGACTCCCTGGGCA

ACCCTACACACATCCTGAGAATCGGCGAGAGCTACAAAGAGAAGCAGAGGACAATCCAGGCCGCCAAAGAGGTGGAA
CAGAGAAGAGCCGGCGGATACTCTAGGAAGTACGCCAGCAAGGCCAAGAATCTGGCCGACGACATGGTCCGAAACAC
CGCCAGAGATCTGCTGTACTACGCCGTGACACAGGACGCCATGCTGATCTTCGAGAATCTGAGCAGAGGCTTCGGCC
GGCAGGGCAAGAGAACCTTTATGGCCGAGAGGCAGTACACCAGAATGGAAGATTGGCTCACAGCTAAACTGGCCTAC
GAGGGACTGCCCAGCAAGACCTACCTGTCCAAAACACTGGCCCAGTATACCTCCCGCACCTGCAGCaattgcggctt
caccatcaccagcgccgactacgacagagtgctggaaaagctcaagaaaaccgccaccggctggatgaccaccatca
acggcaaagagctgaaggttgagggccagatcacctactacaacaggtacaagaggcagaacgtcgtgaaggatctg
agcgtggaactggacagactgagcgaagagagcgtgaacaacgacatcagcagctggacaaagggcagatcaggcga
ggctctgagcctgctgaagaagaggtttagccacagacctgtgcaagagaagttcgtgtgcctgaactgcggcttcg
agacacacgccgatgaacaggctgccctgaacattgccagaagctggctgttcctgagaagccaagagtacaagaag
taccagaccaacaagaccaccggcaacaccgacaagagagggcctttgtggaaacctggcagagcttctacagaaaaaa
gctgaaagaagtctggaagcccgccgtgAATGCATTGCCAAAGAAGAAGCGGAAGGTCGGCAGTTACCCATACGATG
TTCCAGATTACGCTTACCCATACGATGTTCCAGATTACGCTTACCCATACGATGTTCCAGATTACGCTTAAGAATTC
CTAGAGCTCGCTGATCAGCCTCGACTGTGCCTTCTAGTTGCCAGCCATCTGTTGTTTGCCCCTCCCCCGTGCCTTCC
TTGACCCTGGAAGGTGCCACTCCCACTGTCCTTTCCTAATAAAATGAGGAAATTGCATCGCATTGTCTGAGTAGGTG
TCATTCTATTCTGGGGGGTGGGGTGGGGCAGGACAGCAAGGGGGAGGATTGGGAAGAGAATAGCAGGCATGCTGGGG
AGGTACCCTTTTGCTGGCCTTTTGCTCACATGTGAGGGCCTATTTCCCATGATTCCTTCATATTTGCATATACGATA
CAAGGCTGTTAGAGAGATAATTGGAATTAATTTGACTGTAAACACAAAGATATTAGTACAAAATACGTGACGTAGAA
AGTAATAATTTCTTGGGTAGTTTGCAGTTTTAAAATTATGTTTTAAAATGGACTATCATATGCTTACCGTAACTTGA
AAGTATTTCGATTTCTTGGCTTTATATATCTTGTGGAAAGGACGAAACACCGTACTGGCGCTTTTATCTCATTACTT
TGAGAGCCATCACCAGCGACTATGTCGTATGGGTAAAGCGCTTATTTATCGGAGAGAAATCCGATAAATAAGAAGCA
TCAAAGGGTCTTCTCGAAGACATGCCGTTTGCGGCCGCAGGAACCCCTAGTGATGGAGTTGGCCACTCCCTCTCTGC
GCGCTCGCTCGCTCACTGAGGCCGGGCGACCAAAGGTCGCCCGACGCCCGGGCTTTGCCCGGGCGGCCTCAGTGAGC
GAGCGAGCGCGCAGCTGCCTGCAGGGGCGCCTGATGCGGTATTTTCTCCTTACGCATCTGTGCGGTATTTCACACCG
CATACGTCAAAGCAACCATAGTACGCGCCCTGTAGCGGCGCATTAAGCGCGGCGGGTGTGGTGGTTACGCGCAGCGT
GACCGCTACACTTGCCAGCGCCTTAGCGCCCGCTCCTTTCGCTTTCTTCCCTTCCTTTCTCGCCACGTTCGCCGGCT
TTCCCCGTCAAGCTCTAAATCGGGGGCTCCCTTTAGGGTTCCGATTTAGTGCTTTACGGCACCTCGACCCCAAAAAA
CTTGATTTGGGTGATGGTTCACGTAGTGGGCCATCGCCCTGATAGACGGTTTTTCGCCCTTTGACGTTGGAGTCCAC
GTTCTTTAATAGTGGACTCTTGTTCCAAACTGGAACAACACTCAACTCTATCTCGGGCTATTCTTTTGATTTATAAG
GGATTTTGCCGATTTCGGTCTATTGGTTAAAAAATGAGCTGATTTAACAAAAATTTAACGCGAATTTTAACAAAATA
TTAACGTTTACAATTTTATGGTGCACTCTCAGTACAATCTGCTCTGATGCCGCATAGTTAAGCCAGCCCCGACACCC
GCCAACACCCGCTGACGCGCCCTGACGGGCTTGTCTGCTCCCGGCATCCGCTTACAGACAAGCTGTGACCGTCTCCG
GGAGCTGCATGTGTCAGAGGTTTTCACCGTCATCACCGAAACGCGCGAGACGAAAGGGCCTCGTGATACGCCTATTT
TTATAGGTTAATGTCATGATAATAATGGTTTCTTAGACGTCAGGTGGCACTTTTCGGGGAAATGTGCGCGGAACCCC
TATTTGTTTATTTTTCTAAATACATTCAAATATGTATCCGCTCATGAGACAATAACCCTGATAAATGCTTCAATAAT
ATTGAAAAAGGAAGAGTATGAGTATTCAACATTTCCGTGTCGCCCTTATTCCCTTTTTTGCGGCATTTTGCCTTCCT
GTTTTTGCTCACCCAGAAACGCTGGTGAAAGTAAAAGATGCTGAAGATCAGTTGGGTGCACGAGTGGGTTACATCGA
ACTGGATCTCAACAGCGGTAAGATCCTTGAGAGTTTTCGCCCCGAAGAACGTTTTCCAATGATGAGCACTTTTAAAG
TTCTGCTATGTGGCGCGGTATTATCCCGTATTGACGCCGGGCAAGAGCAACTCGGTCGCCGCATACACTATTCTCAG
AATGACTTGGTTGAGTACTCACCAGTCACAGAAAAGCATCTTACGGATGGCATGACAGTAAGAGAATTATGCAGTGC
TGCCATAACCATGAGTGATAACACTGCGGCCAACTTACTTCTGACAACGATCGGAGGACCGAAGGAGCTAACCGCTT
TTTTGCACAACATGGGGGATCATGTAACTCGCCTTGATCGTTGGGAACCGGAGCTGAATGAAGCCATACCAAACGAC
GAGCGTGACACCACGATGCCTGTAGCAATGGCAACAACGTTGCGCAAACTATTAACTGGCGAACTACTTACTCTAGC
TTCCCGGCAACAATTAATAGACTGGATGGAGGCGGATAAAGTTGCAGGACCACTTCTGCGCTCGGCCCTTCCGGCTG
GCTGGTTTATTGCTGATAAATCTGGAGCCGGTGAGCGTGGAAGCCGCGGTATCATTGCAGCACTGGGGCCAGATGGT
AAGCCCTCCCGTATCGTAGTTATCTACACGACGGGGAGTCAGGCAACTATGGATGAACGAAATAGACAGATCGCTGA
GATAGGTGCCTCACTGATTAAGCATTGGTAACTGTCAGACCAAGTTTACTCATATATACTTTAGATTGATTTAAAAC
TTCATTTTTAATTTAAAAGGATCTAGGTGAAGATCCTTTTTGATAATCTCATGACCAAAATCCCTTAACGTGAGTTT
TCGTTCCACTGAGCGTCAGACCCCG

**>CasX2Max_CCR5_sg5_sg10**

TTGAGATCCTTTTTTTCTGCGCGTAATCTGCTGCTTGCAAACAAAAAAACCACCGCTACCAGCGGTGGTTTGTTTGC
CGGATCAAGAGCTACCAACTCTTTTTCCGAAGGTAACTGGCTTCAGCAGAGCGCAGATACCAAATACTGTTCTTCTA
GTGTAGCCGTAGTTAGGCCACCACTTCAAGAACTCTGTAGCACCGCCTACATACCTCGCTCTGCTAATCCTGTTACC
AGTGGCTGCTGCCAGTGGCGATAAGTCGTGTCTTACCGGGTTGGACTCAAGACGATAGTTACCGGATAAGGCGCAGC
GGTCGGGCTGAACGGGGGGTTCGTGCACACAGCCCAGCTTGGAGCGAACGACCTACACCGAACTGAGATACCTACAG
CGTGAGCTATGAGAAAGCGCCACGCTTCCCGAAGGGAGAAAGGCGGACAGGTATCCGGTAAGCGGCAGGGTCGGAAC
AGGAGAGCGCACGAGGGAGCTTCCAGGGGGAAACGCCTGGTATCTTTATAGTCCTGTCGGGTTTCGCCACCTCTGAC
TTGAGCGTCGATTTTTGTGATGCTCGTCAGGGGGGCGGAGCCTATGGAAAAACGCCAGCAACGCGGCCTTTTTACGG
TTCCTGGCCTTTTGCTGGCCTTTTGCTCACATGTCCTGCAGGCAGCTGCGCGCTCGCTCGCTCACTGAGGCCGCCCG
GGCGTCGGGCGACCTTTGGTCGCCCGGCCTCAGTGAGCGAGCGAGCGCGCAGAGAGGGAGTGGCCAACTCCATCACT
AGGGGTTCCTGCGGCCTCTAGACTCGAGGCGTTGACATTGATTATTGACTAGTTATTAATAGTAATCAATTACGGGG
TCATTAGTTCATAGCCCATATATGGAGTTCCGCGTTACATAACTTACGGTAAATGGCCCGCCTGGCTGACCGCCCAA
CGACCCCCGCCCATTGACGTCAATAATGACGTATGTTCCCATAGTAACGCCAATAGGGACTTTCCATTGACGTCAAT
GGGTGGAGTATTTACGGTAAACTGCCCACTTGGCAGTACATCAAGTGTATCATATGCCAAGTACGCCCCCTATTGAC
GTCAATGACGGTAAATGGCCCGCCTGGCATTATGCCCAGTACATGACCTTATGGGACTTTCCTACTTGGCAGTACAT
CTACGTATTAGTCATCGCTATTACCATGGTGATGCGGTTTTGGCAGTACATCAATGGGCGTGGATAGCGGTTTGACT
CACGGGGATTTCCAAGTCTCCACCCCATTGACGTCAATGGGAGTTTGTTTTGGCACCAAAATCAACGGGACTTTCCA
AAATGTCGTAACAACTCCGCCCCATTGACGCAAATGGGCGGTAGGCGTGTACGGTGGGAGGTCTATATAAGCAGAGC
TCTCTGGCTAACTACCGGTTCTAGAGCGCTGCCACCATGTCCGGATCCCCTGCTGCCAAGAGGGTCAAGTTGGACAT
GCAAGAGATCAAGAGAATCAACAAGATCAGAAGGAGACTGGTCAAGGACAGCAACACAAAGAAGGCCGGCAAGCGCG
GCCCCATGAAAACCCTGCTCGTCAGAGTGATGACCCCTGACCTGAGAGAGCGGCTGGAAAACCTGAGAAAGAAGCCC
GAGAACATCCCTCAGCCTATCAGCAACACCAGCAGGGCCAACCTGAACAAGCTGCTGACCGACTACACCGAGATGAA
GAAAGCCATCCTGCACGTGTACTGGGAAGAGTTCCAGAAAGACCCCGTGGGCCTGATGAGCAGAGTTGCTCAGCCCG
CTCCTAAGAACATCGACCAGAGAAAGCTGATCCCCGTGAAGGACGGCAACGAGAGACTGACCTCTAGCGGCTTTGCC
TGCAGCCAGTGTTGCCAGCCTCTGTACGTGTACAAGCTGGAACAAGTGAACGACAAGGGCAAGCCCCACACCAACTA
CTTCGGCAGATGCAACGTGTCCGAGCACGAGAGGCTGATCCTGCTGTCTCCTCACAAGCCCGAGGCCAACGATGAGC
TGGTCACATACAGCCTGGGCAAGTTCGGACAGAGAGCCCTGGACTTCTACAGCATCCACGTGACCAGGGAGAGCAAT
CACCCTGTGAAGCCCCTGGAACAGATCGGCGGCAATAGCTGTGCCTCTGGACCTGTGGGAAAAGCCCTGAGCGACGC
CTGTATGGGAGCCGTGGCATCCTTCCTGACCAAGTACCAGGACATCATCCTGGAACACCAGAAAGTGATCAAGAAGA
ACGAGAAAAGACTGGCCAACCTCAAGGATATCGCCAGCGCTAACGGCCTGGCCTTTCCTAAGATCACCCTGCCTCCA
CAGCCTCACACCAAAGAGGGCATCGAGGCCTACAACAACGTGGTGGCCCAGATCGTGATTTGGGTCAACCTGAATCT
GTGGCAGAAGCTGAAGATCGGCAGGGACGAAGCCAAGCCACTGCAGAGACTGAAGGGCTTCCCTAGCTTCCCTCTGG
TGGAAAGACAGGCCAATGAAGTGGATTGGTGGGACATGGTCTGCAACGTGAAGAAGCTGATCAACGAGAAGAAAGAG
GATGGCAAGGTTTTCTGGCAGAACCTGGCCGGCTACAAGAGACAAGAAGCCCTGCTGCCTTACCTGAGCAGCGAAGA
GGACCGGAAGAAGGGCAAGAAGTTCGCCAGATACCAGTTCGGCGACCTGCTGCTGCACCTGGAAAAGAAGCACGGCG
AGGACTGGGGCAAAGTGTACGATGAGGCCTGGGAGAGAATCGACAAGAAGGTGGAAGGCCTGAGCAAGCACATTAAG
CTGGAAGAGGAAAGAAGGAGCGAGGACGCCCAATCTAAAGCCGCTCTGACCGATTGGCTGAGAGCCAAGGCCAGCTT
TGTGATCGAGGGCCTGAAAGAGGCCGACAAGGACGAGTTCTGCAGATGCGAGCTGAAGCTGCAGAAGTGGTACGGCG
ATCTGAGAGGCAAGCCCTTCGCCATTGAGGCCGAGAACAGCATCCTGGACATCAGCGGCTTCAGCAAGCAGTACAAC
TGCGCCTTCATTTGGCAGAAAGACGGCGTCAAGAAACTGAACCTGTACCTGATCATCAATTACTTCAAAGGCGGCAA
GCTGCGGTTCAAGAAGATCAAACCCGAGGCCTTCGAGGCTAACAGATTCTACACCGTGATCAACAAAAAGTCCGGCG
AGATCGTGCCCATGGAAGTGAACTTCAACTTCGACGACCCCAACCTGATTATCCTGCCTCTGGCCTTCGGCAAGAGA
CAGGGCAGAGAGTTCATCTGGAACGATCTGCTGAGCCTGGAAACCGGCTCTCTGCGCCTGGCCAATGGCAGAGTGAT
CGAGAAAACCCTGTACAACAGGAGAACCAGACAGGACGAGCCTGCTCTGTTTGTGGCCCTGACCTTCGAGAGAAGAG
AGGTGCTGGACAGCAGCAACATCAAGCCCATGAACCTGATCGGCATCGACCGGGGCGAGAATATCCCTGCTGTGATC
GCCCTGACAGACCCTGAAGGATGCCCACTGAGCAGATTCAAGGACTCCCTGGGCAACCCTACACACATCCTGAGAAT

CGGCGAGAGCTACAAAGAGAAGCAGAGGACAATCCAGGCCGCCAAAGAGGTGGAACAGAGAAGAGCCGGCGGATACT
CTAGGAAGTACGCCAGCAAGGCCAAGAATCTGGCCGACGACATGGTCCGAAACACCGCCAGAGATCTGCTGTACTAC
GCCGTGACACAGGACGCCATGCTGATCTTCGAGAATCTGAGCAGAGGCTTCGGCCGGCAGGGCAAGAGAACCTTTAT
GGCCGAGAGGCAGTACACCAGAATGGAAGATTGGCTCACAGCTAAACTGGCCTACGAGGGACTGCCCAGCAAGACCT
ACCTGTCCAAAACACTGGCCCAGTATACCTCCCGCACCTGCAGCAATTGCGGCTTCACCATCACCAGCGCCGACTAC
GACAGAGTGCTGGAAAAGCTCAAGAAAACCGCCACCGGCTGGATGACCACCATCAACGGCAAAGAGCTGAAGGTTGA
GGGCCAGATCACCTACTACAACAGGTACAAGAGGCAGAACGTCGTGAAGGATCTGAGCGTGGAACTGGACAGACTGA
GCGAAGAGAGCGTGAACAACGACATCAGCAGCTGGACAAAGGGCAGATCAGGCGAGGCTCTGAGCCTGCTGAAGAAG
AGGTTTAGCCACAGACCTGTGCAAGAGAAGTTCGTGTGCCTGAACTGCGGCTTCGAGACACACGCCGATGAACAGGC
TGCCCTGAACATTGCCAGAAGCTGGCTGTTCCTGAGAAGCCAAGAGTACAAGAAGTACCAGACCAACAAGACCACCG
GCAACACCGACAAGAGGGCCTTTGTGGAAACCTGGCAGAGCTTCTACAGAAAAAAGCTGAAAGAAGTCTGGAAGCCC
GCCGTGAATGCATTGCCAAAGAAGAAGCGGAAGGTCGGCAGTTACCCATACGATGTGCCTGACTACGCTTATCCATA
CGACGTACCCGATTACGCCTATCCTTACGACGTACCTGACTACGCCTAAGAATTCCTAGAGCTCGCTGATCAGCCTC
GACTGTGCCTTCTAGTTGCCAGCCATCTGTTGTTTGCCCCTCCCCCGTGCCTTCCTTGACCCTGGAAGGTGCCACTC
CCACTGTCCTTTCCTAATAAAATGAGGAAATTGCATCGCATTGTCTGAGTAGGTGTCATTCTATTCTGGGGGGTGGG
GTGGGGCAGGACAGCAAGGGGGAGGATTGGGAAGAGAATAGCAGGCATGCTGGGGAGGTACCCTTTTGCTGGCCTTT
TGCTCACATGTGAGGGCCTATTTCCCATGATTCCTTCATATTTGCATATACGATACAAGGCTGTTAGAGAGATAATT
GGAATTAATTTGACTGTAAACACAAAGATATTAGTACAAAATACGTGACGTAGAAAGTAATAATTTCTTGGGTAGTT
TGCAGTTTTAAAATTATGTTTTAAAATGGACTATCATATGCTTACCGTAACTTGAAAGTATTTCGATTTCTTGGCTT
TATATATCTTGTGGAAAGGACGAAACACCGTACTGGCGCTTTTATCTCATTACTTTGAGAGCCATCACCAGCGACTA
TGTCGTATGGGTAAAGCGCTTATTTATCGGAGAGAAATCCGATAAATAAGAAGCATCAAAGAAAGTCCCACTGGGCG
GCAGCATTTTTTGGTACGGCCGCCTAGTCTGGCCGCGAGGGCCTATTTCCCATGATTCCTTCATATTTGCATATACG
ATACAAGGCTGTTAGAGAGATAATTGGAATTAATTTGACTGTAAACACAAAGATATTAGTACAAAATACGTGACGTA
GAAAGTAATAATTTCTTGGGTAGTTTGCAGTTTTAAAATTATGTTTTAAAATGGACTATCATATGCTTACCGTAACT
TGAAAGTATTTCGATTTCTTGGCTTTATATATCTTGTGGAAAGGACGAAACACCGTACTGGCGCTTTTATCTCATTA
CTTTGAGAGCCATCACCAGCGACTATGTCGTATGGGTAAAGCGCTTATTTATCGGAGAGAAATCCGATAAATAAGAA
GCATCAAAGTGCTCCCCAGTGGATCGGGTTTTTTGCCGTTTGCGGCCGCAGGAACCCCTAGTGATGGAGTTGGCCAC
TCCCTCTCTGCGCGCTCGCTCGCTCACTGAGGCCGGGCGACCAAAGGTCGCCCGACGCCCGGGCTTTGCCCGGGCGG
CCTCAGTGAGCGAGCGAGCGCGCAGCTGCCTGCAGGGGCGCCTGATGCGGTATTTTCTCCTTACGCATCTGTGCGGT
ATTTCACACCGCATACGTCAAAGCAACCATAGTACGCGCCCTGTAGCGGCGCATTAAGCGCGGCGGGTGTGGTGGTT
ACGCGCAGCGTGACCGCTACACTTGCCAGCGCCTTAGCGCCCGCTCCTTTCGCTTTCTTCCCTTCCTTTCTCGCCAC
GTTCGCCGGCTTTCCCCGTCAAGCTCTAAATCGGGGGCTCCCTTTAGGGTTCCGATTTAGTGCTTTACGGCACCTCG
ACCCCAAAAAACTTGATTTGGGTGATGGTTCACGTAGTGGGCCATCGCCCTGATAGACGGTTTTTCGCCCTTTGACG
TTGGAGTCCACGTTCTTTAATAGTGGACTCTTGTTCCAAACTGGAACAACACTCAACTCTATCTCGGGCTATTCTTT
TGATTTATAAGGGATTTTGCCGATTTCGGTCTATTGGTTAAAAAATGAGCTGATTTAACAAAAATTTAACGCGAATT
TTAACAAAATATTAACGTTTACAATTTTATGGTGCACTCTCAGTACAATCTGCTCTGATGCCGCATAGTTAAGCCAG
CCCCGACACCCGCCAACACCCGCTGACGCGCCCTGACGGGCTTGTCTGCTCCCGGCATCCGCTTACAGACAAGCTGT
GACCGTCTCCGGGAGCTGCATGTGTCAGAGGTTTTCACCGTCATCACCGAAACGCGCGAGACGAAAGGGCCTCGTGA
TACGCCTATTTTTATAGGTTAATGTCATGATAATAATGGTTTCTTAGACGTCAGGTGGCACTTTTCGGGGAAATGTG
CGCGGAACCCCTATTTGTTTATTTTTCTAAATACATTCAAATATGTATCCGCTCATGAGACAATAACCCTGATAAAT
GCTTCAATAATATTGAAAAAGGAAGAGTATGAGTATTCAACATTTCCGTGTCGCCCTTATTCCCTTTTTTGCGGCAT
TTTGCCTTCCTGTTTTTGCTCACCCAGAAACGCTGGTGAAAGTAAAAGATGCTGAAGATCAGTTGGGTGCACGAGTG
GGTTACATCGAACTGGATCTCAACAGCGGTAAGATCCTTGAGAGTTTTCGCCCCGAAGAACGTTTTCCAATGATGAG
CACTTTTAAAGTTCTGCTATGTGGCGCGGTATTATCCCGTATTGACGCCGGGCAAGAGCAACTCGGTCGCCGCATAC
ACTATTCTCAGAATGACTTGGTTGAGTACTCACCAGTCACAGAAAAGCATCTTACGGATGGCATGACAGTAAGAGAA
TTATGCAGTGCTGCCATAACCATGAGTGATAACACTGCGGCCAACTTACTTCTGACAACGATCGGAGGACCGAAGGA
GCTAACCGCTTTTTTGCACAACATGGGGGATCATGTAACTCGCCTTGATCGTTGGGAACCGGAGCTGAATGAAGCCA
TACCAAACGACGAGCGTGACACCACGATGCCTGTAGCAATGGCAACAACGTTGCGCAAACTATTAACTGGCGAACTA
CTTACTCTAGCTTCCCGGCAACAATTAATAGACTGGATGGAGGCGGATAAAGTTGCAGGACCACTTCTGCGCTCGGC

CCTTCCGGCTGGCTGGTTTATTGCTGATAAATCTGGAGCCGGTGAGCGTGGAAGCCGCGGTATCATTGCAGCACTGG
GGCCAGATGGTAAGCCCTCCCGTATCGTAGTTATCTACACGACGGGGAGTCAGGCAACTATGGATGAACGAAATAGA
CAGATCGCTGAGATAGGTGCCTCACTGATTAAGCATTGGTAACTGTCAGACCAAGTTTACTCATATATACTTTAGAT
TGATTTAAAACTTCATTTTTAATTTAAAAGGATCTAGGTGAAGATCCTTTTTGATAATCTCATGACCAAAATCCCTT
AACGTGAGTTTTCGTTCCACTGAGCGTCAGACCCCGTAGAAAAGATCAAAGGATCTTC

**>CasX2Max_dual scaffold**

TTGAGATCCTTTTTTTCTGCGCGTAATCTGCTGCTTGCAAACAAAAAAACCACCGCTACCAGCGGTGGTTTGTTTGC
CGGATCAAGAGCTACCAACTCTTTTTCCGAAGGTAACTGGCTTCAGCAGAGCGCAGATACCAAATACTGTTCTTCTA
GTGTAGCCGTAGTTAGGCCACCACTTCAAGAACTCTGTAGCACCGCCTACATACCTCGCTCTGCTAATCCTGTTACC
AGTGGCTGCTGCCAGTGGCGATAAGTCGTGTCTTACCGGGTTGGACTCAAGACGATAGTTACCGGATAAGGCGCAGC
GGTCGGGCTGAACGGGGGGTTCGTGCACACAGCCCAGCTTGGAGCGAACGACCTACACCGAACTGAGATACCTACAG
CGTGAGCTATGAGAAAGCGCCACGCTTCCCGAAGGGAGAAAGGCGGACAGGTATCCGGTAAGCGGCAGGGTCGGAAC
AGGAGAGCGCACGAGGGAGCTTCCAGGGGGAAACGCCTGGTATCTTTATAGTCCTGTCGGGTTTCGCCACCTCTGAC
TTGAGCGTCGATTTTTGTGATGCTCGTCAGGGGGGCGGAGCCTATGGAAAAACGCCAGCAACGCGGCCTTTTTACGG
TTCCTGGCCTTTTGCTGGCCTTTTGCTCACATGTCCTGCAGGCAGCTGCGCGCTCGCTCGCTCACTGAGGCCGCCCG
GGCGTCGGGCGACCTTTGGTCGCCCGGCCTCAGTGAGCGAGCGAGCGCGCAGAGAGGGAGTGGCCAACTCCATCACT
AGGGGTTCCTGCGGCCTCTAGACTCGAGGCGTTGACATTGATTATTGACTAGTTATTAATAGTAATCAATTACGGGG
TCATTAGTTCATAGCCCATATATGGAGTTCCGCGTTACATAACTTACGGTAAATGGCCCGCCTGGCTGACCGCCCAA
CGACCCCCGCCCATTGACGTCAATAATGACGTATGTTCCCATAGTAACGCCAATAGGGACTTTCCATTGACGTCAAT
GGGTGGAGTATTTACGGTAAACTGCCCACTTGGCAGTACATCAAGTGTATCATATGCCAAGTACGCCCCCTATTGAC
GTCAATGACGGTAAATGGCCCGCCTGGCATTATGCCCAGTACATGACCTTATGGGACTTTCCTACTTGGCAGTACAT
CTACGTATTAGTCATCGCTATTACCATGGTGATGCGGTTTTGGCAGTACATCAATGGGCGTGGATAGCGGTTTGACT
CACGGGGATTTCCAAGTCTCCACCCCATTGACGTCAATGGGAGTTTGTTTTGGCACCAAAATCAACGGGACTTTCCA
AAATGTCGTAACAACTCCGCCCCATTGACGCAAATGGGCGGTAGGCGTGTACGGTGGGAGGTCTATATAAGCAGAGC
TCTCTGGCTAACTACCGGTTCTAGAGCGCTGCCACCATGTCCGGATCCCCTGCTGCCAAGAGGGTCAAGTTGGACAT
GCAAGAGATCAAGAGAATCAACAAGATCAGAAGGAGACTGGTCAAGGACAGCAACACAAAGAAGGCCGGCAAGCGCG
GCCCCATGAAAACCCTGCTCGTCAGAGTGATGACCCCTGACCTGAGAGAGCGGCTGGAAAACCTGAGAAAGAAGCCC
GAGAACATCCCTCAGCCTATCAGCAACACCAGCAGGGCCAACCTGAACAAGCTGCTGACCGACTACACCGAGATGAA
GAAAGCCATCCTGCACGTGTACTGGGAAGAGTTCCAGAAAGACCCCGTGGGCCTGATGAGCAGAGTTGCTCAGCCCG
CTCCTAAGAACATCGACCAGAGAAAGCTGATCCCCGTGAAGGACGGCAACGAGAGACTGACCTCTAGCGGCTTTGCC
TGCAGCCAGTGTTGCCAGCCTCTGTACGTGTACAAGCTGGAACAAGTGAACGACAAGGGCAAGCCCCACACCAACTA
CTTCGGCAGATGCAACGTGTCCGAGCACGAGAGGCTGATCCTGCTGTCTCCTCACAAGCCCGAGGCCAACGATGAGC
TGGTCACATACAGCCTGGGCAAGTTCGGACAGAGAGCCCTGGACTTCTACAGCATCCACGTGACCAGGGAGAGCAAT
CACCCTGTGAAGCCCCTGGAACAGATCGGCGGCAATAGCTGTGCCTCTGGACCTGTGGGAAAAGCCCTGAGCGACGC
CTGTATGGGAGCCGTGGCATCCTTCCTGACCAAGTACCAGGACATCATCCTGGAACACCAGAAAGTGATCAAGAAGA
ACGAGAAAAGACTGGCCAACCTCAAGGATATCGCCAGCGCTAACGGCCTGGCCTTTCCTAAGATCACCCTGCCTCCA
CAGCCTCACACCAAAGAGGGCATCGAGGCCTACAACAACGTGGTGGCCCAGATCGTGATTTGGGTCAACCTGAATCT
GTGGCAGAAGCTGAAGATCGGCAGGGACGAAGCCAAGCCACTGCAGAGACTGAAGGGCTTCCCTAGCTTCCCTCTGG
TGGAAAGACAGGCCAATGAAGTGGATTGGTGGGACATGGTCTGCAACGTGAAGAAGCTGATCAACGAGAAGAAAGAG
GATGGCAAGGTTTTCTGGCAGAACCTGGCCGGCTACAAGAGACAAGAAGCCCTGCTGCCTTACCTGAGCAGCGAAGA
GGACCGGAAGAAGGGCAAGAAGTTCGCCAGATACCAGTTCGGCGACCTGCTGCTGCACCTGGAAAAGAAGCACGGCG
AGGACTGGGGCAAAGTGTACGATGAGGCCTGGGAGAGAATCGACAAGAAGGTGGAAGGCCTGAGCAAGCACATTAAG
CTGGAAGAGGAAAGAAGGAGCGAGGACGCCCAATCTAAAGCCGCTCTGACCGATTGGCTGAGAGCCAAGGCCAGCTT
TGTGATCGAGGGCCTGAAAGAGGCCGACAAGGACGAGTTCTGCAGATGCGAGCTGAAGCTGCAGAAGTGGTACGGCG
ATCTGAGAGGCAAGCCCTTCGCCATTGAGGCCGAGAACAGCATCCTGGACATCAGCGGCTTCAGCAAGCAGTACAAC
TGCGCCTTCATTTGGCAGAAAGACGGCGTCAAGAAACTGAACCTGTACCTGATCATCAATTACTTCAAAGGCGGCAA

GCTGCGGTTCAAGAAGATCAAACCCGAGGCCTTCGAGGCTAACAGATTCTACACCGTGATCAACAAAAAGTCCGGCG
AGATCGTGCCCATGGAAGTGAACTTCAACTTCGACGACCCCAACCTGATTATCCTGCCTCTGGCCTTCGGCAAGAGA
CAGGGCAGAGAGTTCATCTGGAACGATCTGCTGAGCCTGGAAACCGGCTCTCTGCGCCTGGCCAATGGCAGAGTGAT
CGAGAAAACCCTGTACAACAGGAGAACCAGACAGGACGAGCCTGCTCTGTTTGTGGCCCTGACCTTCGAGAGAAGAG
AGGTGCTGGACAGCAGCAACATCAAGCCCATGAACCTGATCGGCATCGACCGGGGCGAGAATATCCCTGCTGTGATC
GCCCTGACAGACCCTGAAGGATGCCCACTGAGCAGATTCAAGGACTCCCTGGGCAACCCTACACACATCCTGAGAAT
CGGCGAGAGCTACAAAGAGAAGCAGAGGACAATCCAGGCCGCCAAAGAGGTGGAACAGAGAAGAGCCGGCGGATACT
CTAGGAAGTACGCCAGCAAGGCCAAGAATCTGGCCGACGACATGGTCCGAAACACCGCCAGAGATCTGCTGTACTAC
GCCGTGACACAGGACGCCATGCTGATCTTCGAGAATCTGAGCAGAGGCTTCGGCCGGCAGGGCAAGAGAACCTTTAT
GGCCGAGAGGCAGTACACCAGAATGGAAGATTGGCTCACAGCTAAACTGGCCTACGAGGGACTGCCCAGCAAGACCT
ACCTGTCCAAAACACTGGCCCAGTATACCTCCCGCACCTGCAGCAATTGCGGCTTCACCATCACCAGCGCCGACTAC
GACAGAGTGCTGGAAAAGCTCAAGAAAACCGCCACCGGCTGGATGACCACCATCAACGGCAAAGAGCTGAAGGTTGA
GGGCCAGATCACCTACTACAACAGGTACAAGAGGCAGAACGTCGTGAAGGATCTGAGCGTGGAACTGGACAGACTGA
GCGAAGAGAGCGTGAACAACGACATCAGCAGCTGGACAAAGGGCAGATCAGGCGAGGCTCTGAGCCTGCTGAAGAAG
AGGTTTAGCCACAGACCTGTGCAAGAGAAGTTCGTGTGCCTGAACTGCGGCTTCGAGACACACGCCGATGAACAGGC
TGCCCTGAACATTGCCAGAAGCTGGCTGTTCCTGAGAAGCCAAGAGTACAAGAAGTACCAGACCAACAAGACCACCG
GCAACACCGACAAGAGGGCCTTTGTGGAAACCTGGCAGAGCTTCTACAGAAAAAAGCTGAAAGAAGTCTGGAAGCCC
GCCGTGAATGCATTGCCAAAGAAGAAGCGGAAGGTCGGCAGTTACCCATACGATGTGCCTGACTACGCTTATCCATA
CGACGTACCCGATTACGCCTATCCTTACGACGTACCTGACTACGCCTAAGAATTCCTAGAGCTCGCTGATCAGCCTC
GACTGTGCCTTCTAGTTGCCAGCCATCTGTTGTTTGCCCCTCCCCCGTGCCTTCCTTGACCCTGGAAGGTGCCACTC
CCACTGTCCTTTCCTAATAAAATGAGGAAATTGCATCGCATTGTCTGAGTAGGTGTCATTCTATTCTGGGGGGTGGG
GTGGGGCAGGACAGCAAGGGGGAGGATTGGGAAGAGAATAGCAGGCATGCTGGGGAGGTACCCTTTTGCTGGCCTTT
TGCTCACATGTGAGGGCCTATTTCCCATGATTCCTTCATATTTGCATATACGATACAAGGCTGTTAGAGAGATAATT
GGAATTAATTTGACTGTAAACACAAAGATATTAGTACAAAATACGTGACGTAGAAAGTAATAATTTCTTGGGTAGTT
TGCAGTTTTAAAATTATGTTTTAAAATGGACTATCATATGCTTACCGTAACTTGAAAGTATTTCGATTTCTTGGCTT
TATATATCTTGTGGAAAGGACGAAACACCGTACTGGCGCTTTTATCTCATTACTTTGAGAGCCATCACCAGCGACTA
TGTCGTATGGGTAAAGCGCTTATTTATCGGAGAGAAATCCGATAAATAAGAAGCATCAAGTCTTCGAGAAGACCTTT
TTTGGTACGGCCGCCTAGTCTGGCCGCGAGGGCCTATTTCCCATGATTCCTTCATATTTGCATATACGATACAAGGC
TGTTAGAGAGATAATTGGAATTAATTTGACTGTAAACACAAAGATATTAGTACAAAATACGTGACGTAGAAAGTAAT
AATTTCTTGGGTAGTTTGCAGTTTTAAAATTATGTTTTAAAATGGACTATCATATGCTTACCGTAACTTGAAAGTAT
TTCGATTTCTTGGCTTTATATATCTTGTGGAAAGGACGAAACACCGTACTGGCGCTTTTATCTCATTACTTTGAGAG
CCATCACCAGCGACTATGTCGTATGGGTAAAGCGCTTATTTATCGGAGAGAAATCCGATAAATAAGAAGCATCAAAG
GGTCTTCTCGAAGACATGCCGTTTGCGGCCGCAGGAACCCCTAGTGATGGAGTTGGCCACTCCCTCTCTGCGCGCTC
GCTCGCTCACTGAGGCCGGGCGACCAAAGGTCGCCCGACGCCCGGGCTTTGCCCGGGCGGCCTCAGTGAGCGAGCGA
GCGCGCAGCTGCCTGCAGGGGCGCCTGATGCGGTATTTTCTCCTTACGCATCTGTGCGGTATTTCACACCGCATACG
TCAAAGCAACCATAGTACGCGCCCTGTAGCGGCGCATTAAGCGCGGCGGGTGTGGTGGTTACGCGCAGCGTGACCGC
TACACTTGCCAGCGCCTTAGCGCCCGCTCCTTTCGCTTTCTTCCCTTCCTTTCTCGCCACGTTCGCCGGCTTTCCCC
GTCAAGCTCTAAATCGGGGGCTCCCTTTAGGGTTCCGATTTAGTGCTTTACGGCACCTCGACCCCAAAAAACTTGAT
TTGGGTGATGGTTCACGTAGTGGGCCATCGCCCTGATAGACGGTTTTTCGCCCTTTGACGTTGGAGTCCACGTTCTT
TAATAGTGGACTCTTGTTCCAAACTGGAACAACACTCAACTCTATCTCGGGCTATTCTTTTGATTTATAAGGGATTT
TGCCGATTTCGGTCTATTGGTTAAAAAATGAGCTGATTTAACAAAAATTTAACGCGAATTTTAACAAAATATTAACG
TTTACAATTTTATGGTGCACTCTCAGTACAATCTGCTCTGATGCCGCATAGTTAAGCCAGCCCCGACACCCGCCAAC
ACCCGCTGACGCGCCCTGACGGGCTTGTCTGCTCCCGGCATCCGCTTACAGACAAGCTGTGACCGTCTCCGGGAGCT
GCATGTGTCAGAGGTTTTCACCGTCATCACCGAAACGCGCGAGACGAAAGGGCCTCGTGATACGCCTATTTTTATAG
GTTAATGTCATGATAATAATGGTTTCTTAGACGTCAGGTGGCACTTTTCGGGGAAATGTGCGCGGAACCCCTATTTG
TTTATTTTTCTAAATACATTCAAATATGTATCCGCTCATGAGACAATAACCCTGATAAATGCTTCAATAATATTGAA
AAAGGAAGAGTATGAGTATTCAACATTTCCGTGTCGCCCTTATTCCCTTTTTTGCGGCATTTTGCCTTCCTGTTTTT
GCTCACCCAGAAACGCTGGTGAAAGTAAAAGATGCTGAAGATCAGTTGGGTGCACGAGTGGGTTACATCGAACTGGA
TCTCAACAGCGGTAAGATCCTTGAGAGTTTTCGCCCCGAAGAACGTTTTCCAATGATGAGCACTTTTAAAGTTCTGC

TATGTGGCGCGGTATTATCCCGTATTGACGCCGGGCAAGAGCAACTCGGTCGCCGCATACACTATTCTCAGAATGAC
TTGGTTGAGTACTCACCAGTCACAGAAAAGCATCTTACGGATGGCATGACAGTAAGAGAATTATGCAGTGCTGCCAT
AACCATGAGTGATAACACTGCGGCCAACTTACTTCTGACAACGATCGGAGGACCGAAGGAGCTAACCGCTTTTTTGC
ACAACATGGGGGATCATGTAACTCGCCTTGATCGTTGGGAACCGGAGCTGAATGAAGCCATACCAAACGACGAGCGT
GACACCACGATGCCTGTAGCAATGGCAACAACGTTGCGCAAACTATTAACTGGCGAACTACTTACTCTAGCTTCCCG
GCAACAATTAATAGACTGGATGGAGGCGGATAAAGTTGCAGGACCACTTCTGCGCTCGGCCCTTCCGGCTGGCTGGT
TTATTGCTGATAAATCTGGAGCCGGTGAGCGTGGAAGCCGCGGTATCATTGCAGCACTGGGGCCAGATGGTAAGCCC
TCCCGTATCGTAGTTATCTACACGACGGGGAGTCAGGCAACTATGGATGAACGAAATAGACAGATCGCTGAGATAGG
TGCCTCACTGATTAAGCATTGGTAACTGTCAGACCAAGTTTACTCATATATACTTTAGATTGATTTAAAACTTCATT
TTTAATTTAAAAGGATCTAGGTGAAGATCCTTTTTGATAATCTCATGACCAAAATCCCTTAACGTGAGTTTTCGTTC
CACTGAGCGTCAGACCCCGTAGAAAAGATCAAAGGATCTTC

**>SaCas9_dual scaffold**

TTGAGATCCTTTTTTTCTGCGCGTAATCTGCTGCTTGCAAACAAAAAAACCACCGCTACCAGCGGTGGTTTGTTTGC
CGGATCAAGAGCTACCAACTCTTTTTCCGAAGGTAACTGGCTTCAGCAGAGCGCAGATACCAAATACTGTTCTTCTA
GTGTAGCCGTAGTTAGGCCACCACTTCAAGAACTCTGTAGCACCGCCTACATACCTCGCTCTGCTAATCCTGTTACC
AGTGGCTGCTGCCAGTGGCGATAAGTCGTGTCTTACCGGGTTGGACTCAAGACGATAGTTACCGGATAAGGCGCAGC
GGTCGGGCTGAACGGGGGGTTCGTGCACACAGCCCAGCTTGGAGCGAACGACCTACACCGAACTGAGATACCTACAG
CGTGAGCTATGAGAAAGCGCCACGCTTCCCGAAGGGAGAAAGGCGGACAGGTATCCGGTAAGCGGCAGGGTCGGAAC
AGGAGAGCGCACGAGGGAGCTTCCAGGGGGAAACGCCTGGTATCTTTATAGTCCTGTCGGGTTTCGCCACCTCTGAC
TTGAGCGTCGATTTTTGTGATGCTCGTCAGGGGGGCGGAGCCTATGGAAAAACGCCAGCAACGCGGCCTTTTTACGG
TTCCTGGCCTTTTGCTGGCCTTTTGCTCACATGTCCTGCAGGCAGCTGCGCGCTCGCTCGCTCACTGAGGCCGCCCG
GGCGTCGGGCGACCTTTGGTCGCCCGGCCTCAGTGAGCGAGCGAGCGCGCAGAGAGGGAGTGGCCAACTCCATCACT
AGGGGTTCCTGCGGCCTCTAGACTCGAGGCGTTGACATTGATTATTGACTAGTTATTAATAGTAATCAATTACGGGG
TCATTAGTTCATAGCCCATATATGGAGTTCCGCGTTACATAACTTACGGTAAATGGCCCGCCTGGCTGACCGCCCAA
CGACCCCCGCCCATTGACGTCAATAATGACGTATGTTCCCATAGTAACGCCAATAGGGACTTTCCATTGACGTCAAT
GGGTGGAGTATTTACGGTAAACTGCCCACTTGGCAGTACATCAAGTGTATCATATGCCAAGTACGCCCCCTATTGAC
GTCAATGACGGTAAATGGCCCGCCTGGCATTATGCCCAGTACATGACCTTATGGGACTTTCCTACTTGGCAGTACAT
CTACGTATTAGTCATCGCTATTACCATGGTGATGCGGTTTTGGCAGTACATCAATGGGCGTGGATAGCGGTTTGACT
CACGGGGATTTCCAAGTCTCCACCCCATTGACGTCAATGGGAGTTTGTTTTGGCACCAAAATCAACGGGACTTTCCA
AAATGTCGTAACAACTCCGCCCCATTGACGCAAATGGGCGGTAGGCGTGTACGGTGGGAGGTCTATATAAGCAGAGC
TCTCTGGCTAACTACCGGTGCCACCATGtcCGGAtccgagaccgTGGGTATCCtggtctcCatgcaagagatcaaga
gaatcaacaagatcagaaggagactggtcaaggacagcaacacaaagaaggccggcaagacaggccccatgaaaacc
ctgctcgtcagagtgatgacccctgacctgagagagcggctggaaaacctgagaaagaagcccgagaacatccctca
gcctatcagcaacaccagcagggccaacctgaacaagctgctgaccgactacaccgagatgaagaaagccatcctgc
acgtgtactgggaagagttccagaaagaccccgtgggcctgatgagcagagttgctcagcccgctcctaagaacatc
gaccagagaaagctgatccccgtgaaggacggcaacgagagactgacctctagcggctttgcctgcagccagtgttg
ccagcctctgtacgtgtacaagctggaacaagtgaacgacaagggcaagccccacaccaactacttcggcagatgca
acgtgtccgagcacgagaggctgatcctgctgtctcctcacaagcccgaggccaacgatgagctggtcacatacagc
ctgggcaagttcggacagagagccctggacttctacagcatccacgtgaccagggagagcaatcaccctgtgaagcc
cctggaacagatcggcggcaatagctgtgcctctggacctgtgggaaaagccctgagcgacgcctgtatgggagccg
tggcatccttcctgaccaagtaccaggacatcatcctggaacaccagaaagtgatcaagaagaacgagaaaagactg
gccaacctcaaggatatcgccagcgctaacggcctggcctttcctaagatcaccctgcctccacagcctcacaccaa
agagggcatcgaggcctacaacaacgtggtggcccagatcgtgatttgggtcaacctgaatctgtggcagaagctga
agatcggcagggacgaagccaagccactgcagagactgaagggcttccctagcttccctctggtggaaagacaggcc
aatgaagtggattggtgggacatggtctgcaacgtgaagaagctgatcaacgagaagaaagaggatggcaaggtttt
ctggcagaacctggccggctacaagagacaagaagccctgctgccttacctgagcagcgaagaggaccggaagaagg

gcaagaagttcgccagataccagttcggcgacctgctgctgcacctggaaaagaagcacggcgaggactggggcaaa
gtgtacgatgaggcctgggagagaatcgacaagaaggtggaaggcctgagcaagcacattaagctggaagaggaaag
aaggagcgaggacgcccaatctaaagccgctctgaccgattggctgagagccaaggccagctttgtgatcgagggcc
tgaaagaggccgacaaggacgagttctgcagatgcgagctgaagctgcagaagtggtacggcgatctgagaggcaag
cccttcgccattgaggccgagaacagcatcctggacatcagcggcttcagcaagcagtacaactgcgccttcatttg
gcagaaagacggcgtcaagaaactgaacctgtacctgatcatcaattacttcaaaggcggcaagctgcggttcaaga
agatcaaacccgaggccttcgaggctaacagattctacaccgtgatcaacaaaaagtccggcgagatcgtgcccatg
gaagtgaacttcaacttcgacgaccccaacctgattatcctgcctctggccttcggcaagagacagggcagagagtt
catctggaacgatctgctgagcctggaaaccggctctctgaagctggccaatggcagagtgatcgagaaaaccctgt
acaacaggagaaccagacaggacgagcctgctctgtttgtggccctgaccttcgagagaagagaggtgctggacagc
agcaacatcaagcccatgaacctgatcggcatcgaccggggcgagaatatccctgctgtgatcgccctgacagaccc
tgaaggatgcccactgagcagattcaaggactccctgggcaaccctacacacatcctgagaatcggcgagagctaca
aagagaagcagaggacaatccaggccgccaaagaggtggaacagagaagagccggcggatactctaggaagtacgcc
agcaaggccaagaatctggccgacgacatggtccgaaacaccgccagagatctgctgtactacgccgtgacacagga
cgccatgctgatcttcgagaatctgagcagaggcttcggccggcagggcaagagaacctttatggccgagaggcagt
acaccagaatggaagattggctcacagctaaactggcctacgagggactgcccagcaagacctacctgtccaaaaca
ctggcccagtatacctccaagacctgcagcaattgcggcttcaccatcaccagcgccgactacgacagagtgctgga
aaagctcaagaaaaccgccaccggctggatgaccaccatcaacggcaaagagctgaaggttgagggccagatcacct
actacaacaggtacaagaggcagaacgtcgtgaaggatctgagcgtggaactggacagactgagcgaagagagcgtg
aacaacgacatcagcagctggacaaagggcagatcaggcgaggctctgagcctgctgaagaagaggtttagccacag
acctgtgcaagagaagttcgtgtgcctgaactgcggcttcgagacacacgccgatgaacaggctgccctgaacattg
ccagaagctggctgttcctgagaagccaagagtacaagaagtaccagaccaacaagaccaccggcaacaccgacaag
agggcctttgtggaaacctggcagagcttctacagaaaaaagctgaaagaagtctggaagcccgccgtgaatgcatt
gAAAAGGCCGGCGGCCACGAAAAAGGCCGGCCAGGCAAAAAAGAAAAAGGGATCCTACCCATACGATGTTCCAGATT
ACGCTTACCCATACGATGTTCCAGATTACGCTTACCCATACGATGTTCCAGATTACGCTTAAGAATTCCTAGAGCTC
GCTGATCAGCCTCGACTGTGCCTTCTAGTTGCCAGCCATCTGTTGTTTGCCCCTCCCCCGTGCCTTCCTTGACCCTG
GAAGGTGCCACTCCCACTGTCCTTTCCTAATAAAATGAGGAAATTGCATCGCATTGTCTGAGTAGGTGTCATTCTAT
TCTGGGGGGTGGGGTGGGGCAGGACAGCAAGGGGGAGGATTGGGAAGAGAATAGCAGGCATGCTGGGGAGGTACCCT
TTTGCTGGCCTTTTGCTCACATGTGAGGGCCTATTTCCCATGATTCCTTCATATTTGCATATACGATACAAGGCTGT
TAGAGAGATAATTGGAATTAATTTGACTGTAAACACAAAGATATTAGTACAAAATACGTGACGTAGAAAGTAATAAT
TTCTTGGGTAGTTTGCAGTTTTAAAATTATGTTTTAAAATGGACTATCATATGCTTACCGTAACTTGAAAGTATTTC
GATTTCTTGGCTTTATATATCTTGTGGAAAGGACGAAACACCGTACTGGCGCTTTTATCTCATTACTTTGAGAGCCA
TCACCAGCGACTATGTCGTATGGGTAAAGCGCTTATTTATCGGAGAGAAATCCGATAAATAAGAAGCATCAAGTCTT
CGAGAAGACCTTTTTTGGTACGGCCGCCTAGTCTGGCCGCGAGGGCCTATTTCCCATGATTCCTTCATATTTGCATA
TACGATACAAGGCTGTTAGAGAGATAATTGGAATTAATTTGACTGTAAACACAAAGATATTAGTACAAAATACGTGA
CGTAGAAAGTAATAATTTCTTGGGTAGTTTGCAGTTTTAAAATTATGTTTTAAAATGGACTATCATATGCTTACCGT
AACTTGAAAGTATTTCGATTTCTTGGCTTTATATATCTTGTGGAAAGGACGAAACACCGTACTGGCGCTTTTATCTC
ATTACTTTGAGAGCCATCACCAGCGACTATGTCGTATGGGTAAAGCGCTTATTTATCGGAGAGAAATCCGATAAATA
AGAAGCATCAAAGGGTCTTCTCGAAGACATGCCGTTTGCGGCCGCAGGAACCCCTAGTGATGGAGTTGGCCACTCCC
TCTCTGCGCGCTCGCTCGCTCACTGAGGCCGGGCGACCAAAGGTCGCCCGACGCCCGGGCTTTGCCCGGGCGGCCTC
AGTGAGCGAGCGAGCGCGCAGCTGCCTGCAGGGGCGCCTGATGCGGTATTTTCTCCTTACGCATCTGTGCGGTATTT
CACACCGCATACGTCAAAGCAACCATAGTACGCGCCCTGTAGCGGCGCATTAAGCGCGGCGGGTGTGGTGGTTACGC
GCAGCGTGACCGCTACACTTGCCAGCGCCTTAGCGCCCGCTCCTTTCGCTTTCTTCCCTTCCTTTCTCGCCACGTTC
GCCGGCTTTCCCCGTCAAGCTCTAAATCGGGGGCTCCCTTTAGGGTTCCGATTTAGTGCTTTACGGCACCTCGACCC
CAAAAAACTTGATTTGGGTGATGGTTCACGTAGTGGGCCATCGCCCTGATAGACGGTTTTTCGCCCTTTGACGTTGG
AGTCCACGTTCTTTAATAGTGGACTCTTGTTCCAAACTGGAACAACACTCAACTCTATCTCGGGCTATTCTTTTGAT
TTATAAGGGATTTTGCCGATTTCGGTCTATTGGTTAAAAAATGAGCTGATTTAACAAAAATTTAACGCGAATTTTAA
CAAAATATTAACGTTTACAATTTTATGGTGCACTCTCAGTACAATCTGCTCTGATGCCGCATAGTTAAGCCAGCCCC
GACACCCGCCAACACCCGCTGACGCGCCCTGACGGGCTTGTCTGCTCCCGGCATCCGCTTACAGACAAGCTGTGACC

GTCTCCGGGAGCTGCATGTGTCAGAGGTTTTCACCGTCATCACCGAAACGCGCGAGACGAAAGGGCCTCGTGATACG
CCTATTTTTATAGGTTAATGTCATGATAATAATGGTTTCTTAGACGTCAGGTGGCACTTTTCGGGGAAATGTGCGCG
GAACCCCTATTTGTTTATTTTTCTAAATACATTCAAATATGTATCCGCTCATGAGACAATAACCCTGATAAATGCTT
CAATAATATTGAAAAAGGAAGAGTATGAGTATTCAACATTTCCGTGTCGCCCTTATTCCCTTTTTTGCGGCATTTTG
CCTTCCTGTTTTTGCTCACCCAGAAACGCTGGTGAAAGTAAAAGATGCTGAAGATCAGTTGGGTGCACGAGTGGGTT
ACATCGAACTGGATCTCAACAGCGGTAAGATCCTTGAGAGTTTTCGCCCCGAAGAACGTTTTCCAATGATGAGCACT
TTTAAAGTTCTGCTATGTGGCGCGGTATTATCCCGTATTGACGCCGGGCAAGAGCAACTCGGTCGCCGCATACACTA
TTCTCAGAATGACTTGGTTGAGTACTCACCAGTCACAGAAAAGCATCTTACGGATGGCATGACAGTAAGAGAATTAT
GCAGTGCTGCCATAACCATGAGTGATAACACTGCGGCCAACTTACTTCTGACAACGATCGGAGGACCGAAGGAGCTA
ACCGCTTTTTTGCACAACATGGGGGATCATGTAACTCGCCTTGATCGTTGGGAACCGGAGCTGAATGAAGCCATACC
AAACGACGAGCGTGACACCACGATGCCTGTAGCAATGGCAACAACGTTGCGCAAACTATTAACTGGCGAACTACTTA
CTCTAGCTTCCCGGCAACAATTAATAGACTGGATGGAGGCGGATAAAGTTGCAGGACCACTTCTGCGCTCGGCCCTT
CCGGCTGGCTGGTTTATTGCTGATAAATCTGGAGCCGGTGAGCGTGGAAGCCGCGGTATCATTGCAGCACTGGGGCC
AGATGGTAAGCCCTCCCGTATCGTAGTTATCTACACGACGGGGAGTCAGGCAACTATGGATGAACGAAATAGACAGA
TCGCTGAGATAGGTGCCTCACTGATTAAGCATTGGTAACTGTCAGACCAAGTTTACTCATATATACTTTAGATTGAT
TTAAAACTTCATTTTTAATTTAAAAGGATCTAGGTGAAGATCCTTTTTGATAATCTCATGACCAAAATCCCTTAACG
TGAGTTTTCGTTCCACTGAGCGTCAGACCCCGTAGAAAAGATCAAAGGATCTTC

**>SaCas9_CCR5_Khalili_A_B**

TTGAGATCCTTTTTTTCTGCGCGTAATCTGCTGCTTGCAAACAAAAAAACCACCGCTACCAGCGGTGGTTTGTTTGC
CGGATCAAGAGCTACCAACTCTTTTTCCGAAGGTAACTGGCTTCAGCAGAGCGCAGATACCAAATACTGTTCTTCTA
GTGTAGCCGTAGTTAGGCCACCACTTCAAGAACTCTGTAGCACCGCCTACATACCTCGCTCTGCTAATCCTGTTACC
AGTGGCTGCTGCCAGTGGCGATAAGTCGTGTCTTACCGGGTTGGACTCAAGACGATAGTTACCGGATAAGGCGCAGC
GGTCGGGCTGAACGGGGGGTTCGTGCACACAGCCCAGCTTGGAGCGAACGACCTACACCGAACTGAGATACCTACAG
CGTGAGCTATGAGAAAGCGCCACGCTTCCCGAAGGGAGAAAGGCGGACAGGTATCCGGTAAGCGGCAGGGTCGGAAC
AGGAGAGCGCACGAGGGAGCTTCCAGGGGGAAACGCCTGGTATCTTTATAGTCCTGTCGGGTTTCGCCACCTCTGAC
TTGAGCGTCGATTTTTGTGATGCTCGTCAGGGGGGCGGAGCCTATGGAAAAACGCCAGCAACGCGGCCTTTTTACGG
TTCCTGGCCTTTTGCTGGCCTTTTGCTCACATGTCCTGCAGGCAGCTGCGCGCTCGCTCGCTCACTGAGGCCGCCCG
GGCGTCGGGCGACCTTTGGTCGCCCGGCCTCAGTGAGCGAGCGAGCGCGCAGAGAGGGAGTGGCCAACTCCATCACT
AGGGGTTCCTGCGGCCTCTAGACTCGAGGCGTTGACATTGATTATTGACTAGTTATTAATAGTAATCAATTACGGGG
TCATTAGTTCATAGCCCATATATGGAGTTCCGCGTTACATAACTTACGGTAAATGGCCCGCCTGGCTGACCGCCCAA
CGACCCCCGCCCATTGACGTCAATAATGACGTATGTTCCCATAGTAACGCCAATAGGGACTTTCCATTGACGTCAAT
GGGTGGAGTATTTACGGTAAACTGCCCACTTGGCAGTACATCAAGTGTATCATATGCCAAGTACGCCCCCTATTGAC
GTCAATGACGGTAAATGGCCCGCCTGGCATTATGCCCAGTACATGACCTTATGGGACTTTCCTACTTGGCAGTACAT
CTACGTATTAGTCATCGCTATTACCATGGTGATGCGGTTTTGGCAGTACATCAATGGGCGTGGATAGCGGTTTGACT
CACGGGGATTTCCAAGTCTCCACCCCATTGACGTCAATGGGAGTTTGTTTTGGCACCAAAATCAACGGGACTTTCCA
AAATGTCGTAACAACTCCGCCCCATTGACGCAAATGGGCGGTAGGCGTGTACGGTGGGAGGTCTATATAAGCAGAGC
TCTCTGGCTAACTACCGGTGCCACCATGGCCCCAAAGAAGAAGCGGAAGGTCGGTATCCACGGAGTCCCAGCAGCCA
AGCGGAACTACATCCTGGGCCTGGACATCGGCATCACCAGCGTGGGCTACGGCATCATCGACTACGAGACACGGGAC
GTGATCGATGCCGGCGTGCGGCTGTTCAAAGAGGCCAACGTGGAAAACAACGAGGGCAGGCGGAGCAAGAGAGGCGC
CAGAAGGCTGAAGCGGCGGAGGCGGCATAGAATCCAGAGAGTGAAGAAGCTGCTGTTCGACTACAACCTGCTGACCG
ACCACAGCGAGCTGAGCGGCATCAACCCCTACGAGGCCAGAGTGAAGGGCCTGAGCCAGAAGCTGAGCGAGGAAGAG
TTCTCTGCCGCCCTGCTGCACCTGGCCAAGAGAAGAGGCGTGCACAACGTGAACGAGGTGGAAGAGGACACCGGCAA
CGAGCTGTCCACCAAAGAGCAGATCAGCCGGAACAGCAAGGCCCTGGAAGAGAAATACGTGGCCGAACTGCAGCTGG
AACGGCTGAAGAAAGACGGCGAAGTGCGGGGCAGCATCAACAGATTCAAGACCAGCGACTACGTGAAAGAAGCCAAA
CAGCTGCTGAAGGTGCAGAAGGCCTACCACCAGCTGGACCAGAGCTTCATCGACACCTACATCGACCTGCTGGAAAC
CCGGCGGACCTACTATGAGGGACCTGGCGAGGGCAGCCCCTTCGGCTGGAAGGACATCAAAGAATGGTACGAGATGC

TGATGGGCCACTGCACCTACTTCCCCGAGGAACTGCGGAGCGTGAAGTACGCCTACAACGCCGACCTGTACAACGCC
CTGAACGACCTGAACAATCTCGTGATCACCAGGGACGAGAACGAGAAGCTGGAATATTACGAGAAGTTCCAGATCAT
CGAGAACGTGTTCAAGCAGAAGAAGAAGCCCACCCTGAAGCAGATCGCCAAAGAAATCCTCGTGAACGAAGAGGATA
TTAAGGGCTACAGAGTGACCAGCACCGGCAAGCCCGAGTTCACCAACCTGAAGGTGTACCACGACATCAAGGACATT
ACCGCCCGGAAAGAGATTATTGAGAACGCCGAGCTGCTGGATCAGATTGCCAAGATCCTGACCATCTACCAGAGCAG
CGAGGACATCCAGGAAGAACTGACCAATCTGAACTCCGAGCTGACCCAGGAAGAGATCGAGCAGATCTCTAATCTGA
AGGGCTATACCGGCACCCACAACCTGAGCCTGAAGGCCATCAACCTGATCCTGGACGAGCTGTGGCACACCAACGAC
AACCAGATCGCTATCTTCAACCGGCTGAAGCTGGTGCCCAAGAAGGTGGACCTGTCCCAGCAGAAAGAGATCCCCAC
CACCCTGGTGGACGACTTCATCCTGAGCCCCGTCGTGAAGAGAAGCTTCATCCAGAGCATCAAAGTGATCAACGCCA
TCATCAAGAAGTACGGCCTGCCCAACGACATCATTATCGAGCTGGCCCGCGAGAAGAACTCCAAGGACGCCCAGAAA
ATGATCAACGAGATGCAGAAGCGGAACCGGCAGACCAACGAGCGGATCGAGGAAATCATCCGGACCACCGGCAAAGA
GAACGCCAAGTACCTGATCGAGAAGATCAAGCTGCACGACATGCAGGAAGGCAAGTGCCTGTACAGCCTGGAAGCCA
TCCCTCTGGAAGATCTGCTGAACAACCCCTTCAACTATGAGGTGGACCACATCATCCCCAGAAGCGTGTCCTTCGAC
AACAGCTTCAACAACAAGGTGCTCGTGAAGCAGGAAGAAAACAGCAAGAAGGGCAACCGGACCCCATTCCAGTACCT
GAGCAGCAGCGACAGCAAGATCAGCTACGAAACCTTCAAGAAGCACATCCTGAATCTGGCCAAGGGCAAGGGCAGAA
TCAGCAAGACCAAGAAAGAGTATCTGCTGGAAGAACGGGACATCAACAGGTTCTCCGTGCAGAAAGACTTCATCAAC
CGGAACCTGGTGGATACCAGATACGCCACCAGAGGCCTGATGAACCTGCTGCGGAGCTACTTCAGAGTGAACAACCT
GGACGTGAAAGTGAAGTCCATCAATGGCGGCTTCACCAGCTTTCTGCGGCGGAAGTGGAAGTTTAAGAAAGAGCGGA
ACAAGGGGTACAAGCACCACGCCGAGGACGCCCTGATCATTGCCAACGCCGATTTCATCTTCAAAGAGTGGAAGAAA
CTGGACAAGGCCAAAAAAGTGATGGAAAACCAGATGTTCGAGGAAAAGCAGGCCGAGAGCATGCCCGAGATCGAAAC
CGAGCAGGAGTACAAAGAGATCTTCATCACCCCCCACCAGATCAAGCACATTAAGGACTTCAAGGACTACAAGTACA
GCCACCGGGTGGACAAGAAGCCTAATAGAGAGCTGATTAACGACACCCTGTACTCCACCCGGAAGGACGACAAGGGC
AACACCCTGATCGTGAACAATCTGAACGGCCTGTACGACAAGGACAATGACAAGCTGAAAAAGCTGATCAACAAGAG
CCCCGAAAAGCTGCTGATGTACCACCACGACCCCCAGACCTACCAGAAACTGAAGCTGATTATGGAACAGTACGGCG
ACGAGAAGAATCCCCTGTACAAGTACTACGAGGAAACCGGGAACTACCTGACCAAGTACTCCAAAAAGGACAACGGC
CCCGTGATCAAGAAGATTAAGTATTACGGCAACAAACTGAACGCCCATCTGGACATCACCGACGACTACCCCAACAG
CAGAAACAAGGTCGTGAAGCTGTCCCTGAAGCCCTACAGATTCGACGTGTACCTGGACAATGGCGTGTACAAGTTCG
TGACCGTGAAGAATCTGGATGTGATCAAAAAAGAAAACTACTACGAAGTGAATAGCAAGTGCTATGAGGAAGCTAAG
AAGCTGAAGAAGATCAGCAACCAGGCCGAGTTTATCGCCTCCTTCTACAACAACGATCTGATCAAGATCAACGGCGA
GCTGTATAGAGTGATCGGCGTGAACAACGACCTGCTGAACCGGATCGAAGTGAACATGATCGACATCACCTACCGCG
AGTACCTGGAAAACATGAACGACAAGAGGCCCCCCAGGATCATTAAGACAATCGCCTCCAAGACCCAGAGCATTAAG
AAGTACAGCACAGACATTCTGGGCAACCTGTATGAAGTGAAATCTAAGAAGCACCCTCAGATCATCAAAAAGGGCAA
AAGGCCGGCGGCCACGAAAAAGGCCGGCCAGGCAAAAAAGAAAAAGGGATCCTACCCATACGATGTTCCAGATTACG
CTTACCCATACGATGTTCCAGATTACGCTTACCCATACGATGTTCCAGATTACGCTTAAGAATTCCTAGAGCTCGCT
GATCAGCCTCGACTGTGCCTTCTAGTTGCCAGCCATCTGTTGTTTGCCCCTCCCCCGTGCCTTCCTTGACCCTGGAA
GGTGCCACTCCCACTGTCCTTTCCTAATAAAATGAGGAAATTGCATCGCATTGTCTGAGTAGGTGTCATTCTATTCT
GGGGGGTGGGGTGGGGCAGGACAGCAAGGGGGAGGATTGGGAAGAGAATAGCAGGCATGCTGGGGAGGTACCGAGGG
CCTATTTCCCATGATTCCTTCATATTTGCATATACGATACAAGGCTGTTAGAGAGATAATTGGAATTAATTTGACTG
TAAACACAAAGATATTAGTACAAAATACGTGACGTAGAAAGTAATAATTTCTTGGGTAGTTTGCAGTTTTAAAATTA
TGTTTTAAAATGGACTATCATATGCTTACCGTAACTTGAAAGTATTTCGATTTCTTGGCTTTATATATCTTGTGGAA
AGGACGAAACACCGcggcagcatagtgagcccagGTTTTAGTACTCTGGAAACAGAATCTACTAAAACAAGGCAAAA
TGCCGTGTTTATCTCGTCAACTTGTTGGCGAGATTTTTGCGGCCGTAGTCTGGCCGCGAGGGCCTATTTCCCATGAT
TCCTTCATATTTGCATATACGATACAAGGCTGTTAGAGAGATAATTGGAATTAATTTGACTGTAAACACAAAGATAT
TAGTACAAAATACGTGACGTAGAAAGTAATAATTTCTTGGGTAGTTTGCAGTTTTAAAATTATGTTTTAAAATGGAC
TATCATATGCTTACCGTAACTTGAAAGTATTTCGATTTCTTGGCTTTATATATCTTGTGGAAAGGACGAAACACCGt
cagtttacacccgatccacGTTTTAGTACTCTGGAAACAGAATCTACTAAAACAAGGCAAAATGCCGTGTTTATCTC
GTCAACTTGTTGGCGAGATTTTTGCGGCCGCAGGAACCCCTAGTGATGGAGTTGGCCACTCCCTCTCTGCGCGCTCG
CTCGCTCACTGAGGCCGGGCGACCAAAGGTCGCCCGACGCCCGGGCTTTGCCCGGGCGGCCTCAGTGAGCGAGCGAG
CGCGCAGCTGCCTGCAGGGGCGCCTGATGCGGTATTTTCTCCTTACGCATCTGTGCGGTATTTCACACCGCATACGT

CAAAGCAACCATAGTACGCGCCCTGTAGCGGCGCATTAAGCGCGGCGGGTGTGGTGGTTACGCGCAGCGTGACCGCT
ACACTTGCCAGCGCCTTAGCGCCCGCTCCTTTCGCTTTCTTCCCTTCCTTTCTCGCCACGTTCGCCGGCTTTCCCCG
TCAAGCTCTAAATCGGGGGCTCCCTTTAGGGTTCCGATTTAGTGCTTTACGGCACCTCGACCCCAAAAAACTTGATT
TGGGTGATGGTTCACGTAGTGGGCCATCGCCCTGATAGACGGTTTTTCGCCCTTTGACGTTGGAGTCCACGTTCTTT
AATAGTGGACTCTTGTTCCAAACTGGAACAACACTCAACTCTATCTCGGGCTATTCTTTTGATTTATAAGGGATTTT
GCCGATTTCGGTCTATTGGTTAAAAAATGAGCTGATTTAACAAAAATTTAACGCGAATTTTAACAAAATATTAACGT
TTACAATTTTATGGTGCACTCTCAGTACAATCTGCTCTGATGCCGCATAGTTAAGCCAGCCCCGACACCCGCCAACA
CCCGCTGACGCGCCCTGACGGGCTTGTCTGCTCCCGGCATCCGCTTACAGACAAGCTGTGACCGTCTCCGGGAGCTG
CATGTGTCAGAGGTTTTCACCGTCATCACCGAAACGCGCGAGACGAAAGGGCCTCGTGATACGCCTATTTTTATAGG
TTAATGTCATGATAATAATGGTTTCTTAGACGTCAGGTGGCACTTTTCGGGGAAATGTGCGCGGAACCCCTATTTGT
TTATTTTTCTAAATACATTCAAATATGTATCCGCTCATGAGACAATAACCCTGATAAATGCTTCAATAATATTGAAA
AAGGAAGAGTATGAGTATTCAACATTTCCGTGTCGCCCTTATTCCCTTTTTTGCGGCATTTTGCCTTCCTGTTTTTG
CTCACCCAGAAACGCTGGTGAAAGTAAAAGATGCTGAAGATCAGTTGGGTGCACGAGTGGGTTACATCGAACTGGAT
CTCAACAGCGGTAAGATCCTTGAGAGTTTTCGCCCCGAAGAACGTTTTCCAATGATGAGCACTTTTAAAGTTCTGCT
ATGTGGCGCGGTATTATCCCGTATTGACGCCGGGCAAGAGCAACTCGGTCGCCGCATACACTATTCTCAGAATGACT
TGGTTGAGTACTCACCAGTCACAGAAAAGCATCTTACGGATGGCATGACAGTAAGAGAATTATGCAGTGCTGCCATA
ACCATGAGTGATAACACTGCGGCCAACTTACTTCTGACAACGATCGGAGGACCGAAGGAGCTAACCGCTTTTTTGCA
CAACATGGGGGATCATGTAACTCGCCTTGATCGTTGGGAACCGGAGCTGAATGAAGCCATACCAAACGACGAGCGTG
ACACCACGATGCCTGTAGCAATGGCAACAACGTTGCGCAAACTATTAACTGGCGAACTACTTACTCTAGCTTCCCGG
CAACAATTAATAGACTGGATGGAGGCGGATAAAGTTGCAGGACCACTTCTGCGCTCGGCCCTTCCGGCTGGCTGGTT
TATTGCTGATAAATCTGGAGCCGGTGAGCGTGGAAGCCGCGGTATCATTGCAGCACTGGGGCCAGATGGTAAGCCCT
CCCGTATCGTAGTTATCTACACGACGGGGAGTCAGGCAACTATGGATGAACGAAATAGACAGATCGCTGAGATAGGT
GCCTCACTGATTAAGCATTGGTAACTGTCAGACCAAGTTTACTCATATATACTTTAGATTGATTTAAAACTTCATTT
TTAATTTAAAAGGATCTAGGTGAAGATCCTTTTTGATAATCTCATGACCAAAATCCCTTAACGTGAGTTTTCGTTCC
ACTGAGCGTCAGACCCCGTAGAAAAGATCAAAGGATCTTC

**>SaCas9_CCR5_Khalili_A**

TTGAGATCCTTTTTTTCTGCGCGTAATCTGCTGCTTGCAAACAAAAAAACCACCGCTACCAGCGGTGGTTTGTTTGC
CGGATCAAGAGCTACCAACTCTTTTTCCGAAGGTAACTGGCTTCAGCAGAGCGCAGATACCAAATACTGTTCTTCTA
GTGTAGCCGTAGTTAGGCCACCACTTCAAGAACTCTGTAGCACCGCCTACATACCTCGCTCTGCTAATCCTGTTACC
AGTGGCTGCTGCCAGTGGCGATAAGTCGTGTCTTACCGGGTTGGACTCAAGACGATAGTTACCGGATAAGGCGCAGC
GGTCGGGCTGAACGGGGGGTTCGTGCACACAGCCCAGCTTGGAGCGAACGACCTACACCGAACTGAGATACCTACAG
CGTGAGCTATGAGAAAGCGCCACGCTTCCCGAAGGGAGAAAGGCGGACAGGTATCCGGTAAGCGGCAGGGTCGGAAC
AGGAGAGCGCACGAGGGAGCTTCCAGGGGGAAACGCCTGGTATCTTTATAGTCCTGTCGGGTTTCGCCACCTCTGAC
TTGAGCGTCGATTTTTGTGATGCTCGTCAGGGGGGCGGAGCCTATGGAAAAACGCCAGCAACGCGGCCTTTTTACGG
TTCCTGGCCTTTTGCTGGCCTTTTGCTCACATGTCCTGCAGGCAGCTGCGCGCTCGCTCGCTCACTGAGGCCGCCCG
GGCGTCGGGCGACCTTTGGTCGCCCGGCCTCAGTGAGCGAGCGAGCGCGCAGAGAGGGAGTGGCCAACTCCATCACT
AGGGGTTCCTGCGGCCTCTAGACTCGAGGCGTTGACATTGATTATTGACTAGTTATTAATAGTAATCAATTACGGGG
TCATTAGTTCATAGCCCATATATGGAGTTCCGCGTTACATAACTTACGGTAAATGGCCCGCCTGGCTGACCGCCCAA
CGACCCCCGCCCATTGACGTCAATAATGACGTATGTTCCCATAGTAACGCCAATAGGGACTTTCCATTGACGTCAAT
GGGTGGAGTATTTACGGTAAACTGCCCACTTGGCAGTACATCAAGTGTATCATATGCCAAGTACGCCCCCTATTGAC
GTCAATGACGGTAAATGGCCCGCCTGGCATTATGCCCAGTACATGACCTTATGGGACTTTCCTACTTGGCAGTACAT
CTACGTATTAGTCATCGCTATTACCATGGTGATGCGGTTTTGGCAGTACATCAATGGGCGTGGATAGCGGTTTGACT
CACGGGGATTTCCAAGTCTCCACCCCATTGACGTCAATGGGAGTTTGTTTTGGCACCAAAATCAACGGGACTTTCCA
AAATGTCGTAACAACTCCGCCCCATTGACGCAAATGGGCGGTAGGCGTGTACGGTGGGAGGTCTATATAAGCAGAGC
TCTCTGGCTAACTACCGGTGCCACCATGGCCCCAAAGAAGAAGCGGAAGGTCGGTATCCACGGAGTCCCAGCAGCCA
AGCGGAACTACATCCTGGGCCTGGACATCGGCATCACCAGCGTGGGCTACGGCATCATCGACTACGAGACACGGGAC

GTGATCGATGCCGGCGTGCGGCTGTTCAAAGAGGCCAACGTGGAAAACAACGAGGGCAGGCGGAGCAAGAGAGGCGC
CAGAAGGCTGAAGCGGCGGAGGCGGCATAGAATCCAGAGAGTGAAGAAGCTGCTGTTCGACTACAACCTGCTGACCG
ACCACAGCGAGCTGAGCGGCATCAACCCCTACGAGGCCAGAGTGAAGGGCCTGAGCCAGAAGCTGAGCGAGGAAGAG
TTCTCTGCCGCCCTGCTGCACCTGGCCAAGAGAAGAGGCGTGCACAACGTGAACGAGGTGGAAGAGGACACCGGCAA
CGAGCTGTCCACCAAAGAGCAGATCAGCCGGAACAGCAAGGCCCTGGAAGAGAAATACGTGGCCGAACTGCAGCTGG
AACGGCTGAAGAAAGACGGCGAAGTGCGGGGCAGCATCAACAGATTCAAGACCAGCGACTACGTGAAAGAAGCCAAA
CAGCTGCTGAAGGTGCAGAAGGCCTACCACCAGCTGGACCAGAGCTTCATCGACACCTACATCGACCTGCTGGAAAC
CCGGCGGACCTACTATGAGGGACCTGGCGAGGGCAGCCCCTTCGGCTGGAAGGACATCAAAGAATGGTACGAGATGC
TGATGGGCCACTGCACCTACTTCCCCGAGGAACTGCGGAGCGTGAAGTACGCCTACAACGCCGACCTGTACAACGCC
CTGAACGACCTGAACAATCTCGTGATCACCAGGGACGAGAACGAGAAGCTGGAATATTACGAGAAGTTCCAGATCAT
CGAGAACGTGTTCAAGCAGAAGAAGAAGCCCACCCTGAAGCAGATCGCCAAAGAAATCCTCGTGAACGAAGAGGATA
TTAAGGGCTACAGAGTGACCAGCACCGGCAAGCCCGAGTTCACCAACCTGAAGGTGTACCACGACATCAAGGACATT
ACCGCCCGGAAAGAGATTATTGAGAACGCCGAGCTGCTGGATCAGATTGCCAAGATCCTGACCATCTACCAGAGCAG
CGAGGACATCCAGGAAGAACTGACCAATCTGAACTCCGAGCTGACCCAGGAAGAGATCGAGCAGATCTCTAATCTGA
AGGGCTATACCGGCACCCACAACCTGAGCCTGAAGGCCATCAACCTGATCCTGGACGAGCTGTGGCACACCAACGAC
AACCAGATCGCTATCTTCAACCGGCTGAAGCTGGTGCCCAAGAAGGTGGACCTGTCCCAGCAGAAAGAGATCCCCAC
CACCCTGGTGGACGACTTCATCCTGAGCCCCGTCGTGAAGAGAAGCTTCATCCAGAGCATCAAAGTGATCAACGCCA
TCATCAAGAAGTACGGCCTGCCCAACGACATCATTATCGAGCTGGCCCGCGAGAAGAACTCCAAGGACGCCCAGAAA
ATGATCAACGAGATGCAGAAGCGGAACCGGCAGACCAACGAGCGGATCGAGGAAATCATCCGGACCACCGGCAAAGA
GAACGCCAAGTACCTGATCGAGAAGATCAAGCTGCACGACATGCAGGAAGGCAAGTGCCTGTACAGCCTGGAAGCCA
TCCCTCTGGAAGATCTGCTGAACAACCCCTTCAACTATGAGGTGGACCACATCATCCCCAGAAGCGTGTCCTTCGAC
AACAGCTTCAACAACAAGGTGCTCGTGAAGCAGGAAGAAAACAGCAAGAAGGGCAACCGGACCCCATTCCAGTACCT
GAGCAGCAGCGACAGCAAGATCAGCTACGAAACCTTCAAGAAGCACATCCTGAATCTGGCCAAGGGCAAGGGCAGAA
TCAGCAAGACCAAGAAAGAGTATCTGCTGGAAGAACGGGACATCAACAGGTTCTCCGTGCAGAAAGACTTCATCAAC
CGGAACCTGGTGGATACCAGATACGCCACCAGAGGCCTGATGAACCTGCTGCGGAGCTACTTCAGAGTGAACAACCT
GGACGTGAAAGTGAAGTCCATCAATGGCGGCTTCACCAGCTTTCTGCGGCGGAAGTGGAAGTTTAAGAAAGAGCGGA
ACAAGGGGTACAAGCACCACGCCGAGGACGCCCTGATCATTGCCAACGCCGATTTCATCTTCAAAGAGTGGAAGAAA
CTGGACAAGGCCAAAAAAGTGATGGAAAACCAGATGTTCGAGGAAAAGCAGGCCGAGAGCATGCCCGAGATCGAAAC
CGAGCAGGAGTACAAAGAGATCTTCATCACCCCCCACCAGATCAAGCACATTAAGGACTTCAAGGACTACAAGTACA
GCCACCGGGTGGACAAGAAGCCTAATAGAGAGCTGATTAACGACACCCTGTACTCCACCCGGAAGGACGACAAGGGC
AACACCCTGATCGTGAACAATCTGAACGGCCTGTACGACAAGGACAATGACAAGCTGAAAAAGCTGATCAACAAGAG
CCCCGAAAAGCTGCTGATGTACCACCACGACCCCCAGACCTACCAGAAACTGAAGCTGATTATGGAACAGTACGGCG
ACGAGAAGAATCCCCTGTACAAGTACTACGAGGAAACCGGGAACTACCTGACCAAGTACTCCAAAAAGGACAACGGC
CCCGTGATCAAGAAGATTAAGTATTACGGCAACAAACTGAACGCCCATCTGGACATCACCGACGACTACCCCAACAG
CAGAAACAAGGTCGTGAAGCTGTCCCTGAAGCCCTACAGATTCGACGTGTACCTGGACAATGGCGTGTACAAGTTCG
TGACCGTGAAGAATCTGGATGTGATCAAAAAAGAAAACTACTACGAAGTGAATAGCAAGTGCTATGAGGAAGCTAAG
AAGCTGAAGAAGATCAGCAACCAGGCCGAGTTTATCGCCTCCTTCTACAACAACGATCTGATCAAGATCAACGGCGA
GCTGTATAGAGTGATCGGCGTGAACAACGACCTGCTGAACCGGATCGAAGTGAACATGATCGACATCACCTACCGCG
AGTACCTGGAAAACATGAACGACAAGAGGCCCCCCAGGATCATTAAGACAATCGCCTCCAAGACCCAGAGCATTAAG
AAGTACAGCACAGACATTCTGGGCAACCTGTATGAAGTGAAATCTAAGAAGCACCCTCAGATCATCAAAAAGGGCAA
AAGGCCGGCGGCCACGAAAAAGGCCGGCCAGGCAAAAAAGAAAAAGGGATCCTACCCATACGATGTTCCAGATTACG
CTTACCCATACGATGTTCCAGATTACGCTTACCCATACGATGTTCCAGATTACGCTTAAGAATTCCTAGAGCTCGCT
GATCAGCCTCGACTGTGCCTTCTAGTTGCCAGCCATCTGTTGTTTGCCCCTCCCCCGTGCCTTCCTTGACCCTGGAA
GGTGCCACTCCCACTGTCCTTTCCTAATAAAATGAGGAAATTGCATCGCATTGTCTGAGTAGGTGTCATTCTATTCT
GGGGGGTGGGGTGGGGCAGGACAGCAAGGGGGAGGATTGGGAAGAGAATAGCAGGCATGCTGGGGAGGTACCGAGGG
CCTATTTCCCATGATTCCTTCATATTTGCATATACGATACAAGGCTGTTAGAGAGATAATTGGAATTAATTTGACTG
TAAACACAAAGATATTAGTACAAAATACGTGACGTAGAAAGTAATAATTTCTTGGGTAGTTTGCAGTTTTAAAATTA
TGTTTTAAAATGGACTATCATATGCTTACCGTAACTTGAAAGTATTTCGATTTCTTGGCTTTATATATCTTGTGGAA
AGGACGAAACACCGCGGCAGCATAGTGAGCCCAGGTTTTAGTACTCTGGAAACAGAATCTACTAAAACAAGGCAAAA

TGCCGTGTTTATCTCGTCAACTTGTTGGCGAGATTTTTGCGGCCGCAGGAACCCCTAGTGATGGAGTTGGCCACTCC
CTCTCTGCGCGCTCGCTCGCTCACTGAGGCCGGGCGACCAAAGGTCGCCCGACGCCCGGGCTTTGCCCGGGCGGCCT
CAGTGAGCGAGCGAGCGCGCAGCTGCCTGCAGGGGCGCCTGATGCGGTATTTTCTCCTTACGCATCTGTGCGGTATT
TCACACCGCATACGTCAAAGCAACCATAGTACGCGCCCTGTAGCGGCGCATTAAGCGCGGCGGGTGTGGTGGTTACG
CGCAGCGTGACCGCTACACTTGCCAGCGCCTTAGCGCCCGCTCCTTTCGCTTTCTTCCCTTCCTTTCTCGCCACGTT
CGCCGGCTTTCCCCGTCAAGCTCTAAATCGGGGGCTCCCTTTAGGGTTCCGATTTAGTGCTTTACGGCACCTCGACC
CCAAAAAACTTGATTTGGGTGATGGTTCACGTAGTGGGCCATCGCCCTGATAGACGGTTTTTCGCCCTTTGACGTTG
GAGTCCACGTTCTTTAATAGTGGACTCTTGTTCCAAACTGGAACAACACTCAACTCTATCTCGGGCTATTCTTTTGA
TTTATAAGGGATTTTGCCGATTTCGGTCTATTGGTTAAAAAATGAGCTGATTTAACAAAAATTTAACGCGAATTTTA
ACAAAATATTAACGTTTACAATTTTATGGTGCACTCTCAGTACAATCTGCTCTGATGCCGCATAGTTAAGCCAGCCC
CGACACCCGCCAACACCCGCTGACGCGCCCTGACGGGCTTGTCTGCTCCCGGCATCCGCTTACAGACAAGCTGTGAC
CGTCTCCGGGAGCTGCATGTGTCAGAGGTTTTCACCGTCATCACCGAAACGCGCGAGACGAAAGGGCCTCGTGATAC
GCCTATTTTTATAGGTTAATGTCATGATAATAATGGTTTCTTAGACGTCAGGTGGCACTTTTCGGGGAAATGTGCGC
GGAACCCCTATTTGTTTATTTTTCTAAATACATTCAAATATGTATCCGCTCATGAGACAATAACCCTGATAAATGCT
TCAATAATATTGAAAAAGGAAGAGTATGAGTATTCAACATTTCCGTGTCGCCCTTATTCCCTTTTTTGCGGCATTTT
GCCTTCCTGTTTTTGCTCACCCAGAAACGCTGGTGAAAGTAAAAGATGCTGAAGATCAGTTGGGTGCACGAGTGGGT
TACATCGAACTGGATCTCAACAGCGGTAAGATCCTTGAGAGTTTTCGCCCCGAAGAACGTTTTCCAATGATGAGCAC
TTTTAAAGTTCTGCTATGTGGCGCGGTATTATCCCGTATTGACGCCGGGCAAGAGCAACTCGGTCGCCGCATACACT
ATTCTCAGAATGACTTGGTTGAGTACTCACCAGTCACAGAAAAGCATCTTACGGATGGCATGACAGTAAGAGAATTA
TGCAGTGCTGCCATAACCATGAGTGATAACACTGCGGCCAACTTACTTCTGACAACGATCGGAGGACCGAAGGAGCT
AACCGCTTTTTTGCACAACATGGGGGATCATGTAACTCGCCTTGATCGTTGGGAACCGGAGCTGAATGAAGCCATAC
CAAACGACGAGCGTGACACCACGATGCCTGTAGCAATGGCAACAACGTTGCGCAAACTATTAACTGGCGAACTACTT
ACTCTAGCTTCCCGGCAACAATTAATAGACTGGATGGAGGCGGATAAAGTTGCAGGACCACTTCTGCGCTCGGCCCT
TCCGGCTGGCTGGTTTATTGCTGATAAATCTGGAGCCGGTGAGCGTGGAAGCCGCGGTATCATTGCAGCACTGGGGC
CAGATGGTAAGCCCTCCCGTATCGTAGTTATCTACACGACGGGGAGTCAGGCAACTATGGATGAACGAAATAGACAG
ATCGCTGAGATAGGTGCCTCACTGATTAAGCATTGGTAACTGTCAGACCAAGTTTACTCATATATACTTTAGATTGA
TTTAAAACTTCATTTTTAATTTAAAAGGATCTAGGTGAAGATCCTTTTTGATAATCTCATGACCAAAATCCCTTAAC
GTGAGTTTTCGTTCCACTGAGCGTCAGACCCCGTAGAAAAGATCAAAGGATCTTC

**>SaCas9_CCR5_Khalili_B**

TTGAGATCCTTTTTTTCTGCGCGTAATCTGCTGCTTGCAAACAAAAAAACCACCGCTACCAGCGGTGGTTTGTTTGC
CGGATCAAGAGCTACCAACTCTTTTTCCGAAGGTAACTGGCTTCAGCAGAGCGCAGATACCAAATACTGTTCTTCTA
GTGTAGCCGTAGTTAGGCCACCACTTCAAGAACTCTGTAGCACCGCCTACATACCTCGCTCTGCTAATCCTGTTACC
AGTGGCTGCTGCCAGTGGCGATAAGTCGTGTCTTACCGGGTTGGACTCAAGACGATAGTTACCGGATAAGGCGCAGC
GGTCGGGCTGAACGGGGGGTTCGTGCACACAGCCCAGCTTGGAGCGAACGACCTACACCGAACTGAGATACCTACAG
CGTGAGCTATGAGAAAGCGCCACGCTTCCCGAAGGGAGAAAGGCGGACAGGTATCCGGTAAGCGGCAGGGTCGGAAC
AGGAGAGCGCACGAGGGAGCTTCCAGGGGGAAACGCCTGGTATCTTTATAGTCCTGTCGGGTTTCGCCACCTCTGAC
TTGAGCGTCGATTTTTGTGATGCTCGTCAGGGGGGCGGAGCCTATGGAAAAACGCCAGCAACGCGGCCTTTTTACGG
TTCCTGGCCTTTTGCTGGCCTTTTGCTCACATGTCCTGCAGGCAGCTGCGCGCTCGCTCGCTCACTGAGGCCGCCCG
GGCGTCGGGCGACCTTTGGTCGCCCGGCCTCAGTGAGCGAGCGAGCGCGCAGAGAGGGAGTGGCCAACTCCATCACT
AGGGGTTCCTGCGGCCTCTAGACTCGAGGCGTTGACATTGATTATTGACTAGTTATTAATAGTAATCAATTACGGGG
TCATTAGTTCATAGCCCATATATGGAGTTCCGCGTTACATAACTTACGGTAAATGGCCCGCCTGGCTGACCGCCCAA
CGACCCCCGCCCATTGACGTCAATAATGACGTATGTTCCCATAGTAACGCCAATAGGGACTTTCCATTGACGTCAAT
GGGTGGAGTATTTACGGTAAACTGCCCACTTGGCAGTACATCAAGTGTATCATATGCCAAGTACGCCCCCTATTGAC
GTCAATGACGGTAAATGGCCCGCCTGGCATTATGCCCAGTACATGACCTTATGGGACTTTCCTACTTGGCAGTACAT
CTACGTATTAGTCATCGCTATTACCATGGTGATGCGGTTTTGGCAGTACATCAATGGGCGTGGATAGCGGTTTGACT
CACGGGGATTTCCAAGTCTCCACCCCATTGACGTCAATGGGAGTTTGTTTTGGCACCAAAATCAACGGGACTTTCCA

AAATGTCGTAACAACTCCGCCCCATTGACGCAAATGGGCGGTAGGCGTGTACGGTGGGAGGTCTATATAAGCAGAGC
TCTCTGGCTAACTACCGGTGCCACCATGGCCCCAAAGAAGAAGCGGAAGGTCGGTATCCACGGAGTCCCAGCAGCCA
AGCGGAACTACATCCTGGGCCTGGACATCGGCATCACCAGCGTGGGCTACGGCATCATCGACTACGAGACACGGGAC
GTGATCGATGCCGGCGTGCGGCTGTTCAAAGAGGCCAACGTGGAAAACAACGAGGGCAGGCGGAGCAAGAGAGGCGC
CAGAAGGCTGAAGCGGCGGAGGCGGCATAGAATCCAGAGAGTGAAGAAGCTGCTGTTCGACTACAACCTGCTGACCG
ACCACAGCGAGCTGAGCGGCATCAACCCCTACGAGGCCAGAGTGAAGGGCCTGAGCCAGAAGCTGAGCGAGGAAGAG
TTCTCTGCCGCCCTGCTGCACCTGGCCAAGAGAAGAGGCGTGCACAACGTGAACGAGGTGGAAGAGGACACCGGCAA
CGAGCTGTCCACCAAAGAGCAGATCAGCCGGAACAGCAAGGCCCTGGAAGAGAAATACGTGGCCGAACTGCAGCTGG
AACGGCTGAAGAAAGACGGCGAAGTGCGGGGCAGCATCAACAGATTCAAGACCAGCGACTACGTGAAAGAAGCCAAA
CAGCTGCTGAAGGTGCAGAAGGCCTACCACCAGCTGGACCAGAGCTTCATCGACACCTACATCGACCTGCTGGAAAC
CCGGCGGACCTACTATGAGGGACCTGGCGAGGGCAGCCCCTTCGGCTGGAAGGACATCAAAGAATGGTACGAGATGC
TGATGGGCCACTGCACCTACTTCCCCGAGGAACTGCGGAGCGTGAAGTACGCCTACAACGCCGACCTGTACAACGCC
CTGAACGACCTGAACAATCTCGTGATCACCAGGGACGAGAACGAGAAGCTGGAATATTACGAGAAGTTCCAGATCAT
CGAGAACGTGTTCAAGCAGAAGAAGAAGCCCACCCTGAAGCAGATCGCCAAAGAAATCCTCGTGAACGAAGAGGATA
TTAAGGGCTACAGAGTGACCAGCACCGGCAAGCCCGAGTTCACCAACCTGAAGGTGTACCACGACATCAAGGACATT
ACCGCCCGGAAAGAGATTATTGAGAACGCCGAGCTGCTGGATCAGATTGCCAAGATCCTGACCATCTACCAGAGCAG
CGAGGACATCCAGGAAGAACTGACCAATCTGAACTCCGAGCTGACCCAGGAAGAGATCGAGCAGATCTCTAATCTGA
AGGGCTATACCGGCACCCACAACCTGAGCCTGAAGGCCATCAACCTGATCCTGGACGAGCTGTGGCACACCAACGAC
AACCAGATCGCTATCTTCAACCGGCTGAAGCTGGTGCCCAAGAAGGTGGACCTGTCCCAGCAGAAAGAGATCCCCAC
CACCCTGGTGGACGACTTCATCCTGAGCCCCGTCGTGAAGAGAAGCTTCATCCAGAGCATCAAAGTGATCAACGCCA
TCATCAAGAAGTACGGCCTGCCCAACGACATCATTATCGAGCTGGCCCGCGAGAAGAACTCCAAGGACGCCCAGAAA
ATGATCAACGAGATGCAGAAGCGGAACCGGCAGACCAACGAGCGGATCGAGGAAATCATCCGGACCACCGGCAAAGA
GAACGCCAAGTACCTGATCGAGAAGATCAAGCTGCACGACATGCAGGAAGGCAAGTGCCTGTACAGCCTGGAAGCCA
TCCCTCTGGAAGATCTGCTGAACAACCCCTTCAACTATGAGGTGGACCACATCATCCCCAGAAGCGTGTCCTTCGAC
AACAGCTTCAACAACAAGGTGCTCGTGAAGCAGGAAGAAAACAGCAAGAAGGGCAACCGGACCCCATTCCAGTACCT
GAGCAGCAGCGACAGCAAGATCAGCTACGAAACCTTCAAGAAGCACATCCTGAATCTGGCCAAGGGCAAGGGCAGAA
TCAGCAAGACCAAGAAAGAGTATCTGCTGGAAGAACGGGACATCAACAGGTTCTCCGTGCAGAAAGACTTCATCAAC
CGGAACCTGGTGGATACCAGATACGCCACCAGAGGCCTGATGAACCTGCTGCGGAGCTACTTCAGAGTGAACAACCT
GGACGTGAAAGTGAAGTCCATCAATGGCGGCTTCACCAGCTTTCTGCGGCGGAAGTGGAAGTTTAAGAAAGAGCGGA
ACAAGGGGTACAAGCACCACGCCGAGGACGCCCTGATCATTGCCAACGCCGATTTCATCTTCAAAGAGTGGAAGAAA
CTGGACAAGGCCAAAAAAGTGATGGAAAACCAGATGTTCGAGGAAAAGCAGGCCGAGAGCATGCCCGAGATCGAAAC
CGAGCAGGAGTACAAAGAGATCTTCATCACCCCCCACCAGATCAAGCACATTAAGGACTTCAAGGACTACAAGTACA
GCCACCGGGTGGACAAGAAGCCTAATAGAGAGCTGATTAACGACACCCTGTACTCCACCCGGAAGGACGACAAGGGC
AACACCCTGATCGTGAACAATCTGAACGGCCTGTACGACAAGGACAATGACAAGCTGAAAAAGCTGATCAACAAGAG
CCCCGAAAAGCTGCTGATGTACCACCACGACCCCCAGACCTACCAGAAACTGAAGCTGATTATGGAACAGTACGGCG
ACGAGAAGAATCCCCTGTACAAGTACTACGAGGAAACCGGGAACTACCTGACCAAGTACTCCAAAAAGGACAACGGC
CCCGTGATCAAGAAGATTAAGTATTACGGCAACAAACTGAACGCCCATCTGGACATCACCGACGACTACCCCAACAG
CAGAAACAAGGTCGTGAAGCTGTCCCTGAAGCCCTACAGATTCGACGTGTACCTGGACAATGGCGTGTACAAGTTCG
TGACCGTGAAGAATCTGGATGTGATCAAAAAAGAAAACTACTACGAAGTGAATAGCAAGTGCTATGAGGAAGCTAAG
AAGCTGAAGAAGATCAGCAACCAGGCCGAGTTTATCGCCTCCTTCTACAACAACGATCTGATCAAGATCAACGGCGA
GCTGTATAGAGTGATCGGCGTGAACAACGACCTGCTGAACCGGATCGAAGTGAACATGATCGACATCACCTACCGCG
AGTACCTGGAAAACATGAACGACAAGAGGCCCCCCAGGATCATTAAGACAATCGCCTCCAAGACCCAGAGCATTAAG
AAGTACAGCACAGACATTCTGGGCAACCTGTATGAAGTGAAATCTAAGAAGCACCCTCAGATCATCAAAAAGGGCAA
AAGGCCGGCGGCCACGAAAAAGGCCGGCCAGGCAAAAAAGAAAAAGGGATCCTACCCATACGATGTTCCAGATTACG
CTTACCCATACGATGTTCCAGATTACGCTTACCCATACGATGTTCCAGATTACGCTTAAGAATTCCTAGAGCTCGCT
GATCAGCCTCGACTGTGCCTTCTAGTTGCCAGCCATCTGTTGTTTGCCCCTCCCCCGTGCCTTCCTTGACCCTGGAA
GGTGCCACTCCCACTGTCCTTTCCTAATAAAATGAGGAAATTGCATCGCATTGTCTGAGTAGGTGTCATTCTATTCT
GGGGGGTGGGGTGGGGCAGGACAGCAAGGGGGAGGATTGGGAAGAGAATAGCAGGCATGCTGGGGAGGTACCGAGGG
CCTATTTCCCATGATTCCTTCATATTTGCATATACGATACAAGGCTGTTAGAGAGATAATTGGAATTAATTTGACTG

TAAACACAAAGATATTAGTACAAAATACGTGACGTAGAAAGTAATAATTTCTTGGGTAGTTTGCAGTTTTAAAATTA
TGTTTTAAAATGGACTATCATATGCTTACCGTAACTTGAAAGTATTTCGATTTCTTGGCTTTATATATCTTGTGGAA
AGGACGAAACACCGTCAGTTTACACCCGATCCACGTTTTAGTACTCTGGAAACAGAATCTACTAAAACAAGGCAAAA
TGCCGTGTTTATCTCGTCAACTTGTTGGCGAGATTTTTGCGGCCGCAGGAACCCCTAGTGATGGAGTTGGCCACTCC
CTCTCTGCGCGCTCGCTCGCTCACTGAGGCCGGGCGACCAAAGGTCGCCCGACGCCCGGGCTTTGCCCGGGCGGCCT
CAGTGAGCGAGCGAGCGCGCAGCTGCCTGCAGGGGCGCCTGATGCGGTATTTTCTCCTTACGCATCTGTGCGGTATT
TCACACCGCATACGTCAAAGCAACCATAGTACGCGCCCTGTAGCGGCGCATTAAGCGCGGCGGGTGTGGTGGTTACG
CGCAGCGTGACCGCTACACTTGCCAGCGCCTTAGCGCCCGCTCCTTTCGCTTTCTTCCCTTCCTTTCTCGCCACGTT
CGCCGGCTTTCCCCGTCAAGCTCTAAATCGGGGGCTCCCTTTAGGGTTCCGATTTAGTGCTTTACGGCACCTCGACC
CCAAAAAACTTGATTTGGGTGATGGTTCACGTAGTGGGCCATCGCCCTGATAGACGGTTTTTCGCCCTTTGACGTTG
GAGTCCACGTTCTTTAATAGTGGACTCTTGTTCCAAACTGGAACAACACTCAACTCTATCTCGGGCTATTCTTTTGA
TTTATAAGGGATTTTGCCGATTTCGGTCTATTGGTTAAAAAATGAGCTGATTTAACAAAAATTTAACGCGAATTTTA
ACAAAATATTAACGTTTACAATTTTATGGTGCACTCTCAGTACAATCTGCTCTGATGCCGCATAGTTAAGCCAGCCC
CGACACCCGCCAACACCCGCTGACGCGCCCTGACGGGCTTGTCTGCTCCCGGCATCCGCTTACAGACAAGCTGTGAC
CGTCTCCGGGAGCTGCATGTGTCAGAGGTTTTCACCGTCATCACCGAAACGCGCGAGACGAAAGGGCCTCGTGATAC
GCCTATTTTTATAGGTTAATGTCATGATAATAATGGTTTCTTAGACGTCAGGTGGCACTTTTCGGGGAAATGTGCGC
GGAACCCCTATTTGTTTATTTTTCTAAATACATTCAAATATGTATCCGCTCATGAGACAATAACCCTGATAAATGCT
TCAATAATATTGAAAAAGGAAGAGTATGAGTATTCAACATTTCCGTGTCGCCCTTATTCCCTTTTTTGCGGCATTTT
GCCTTCCTGTTTTTGCTCACCCAGAAACGCTGGTGAAAGTAAAAGATGCTGAAGATCAGTTGGGTGCACGAGTGGGT
TACATCGAACTGGATCTCAACAGCGGTAAGATCCTTGAGAGTTTTCGCCCCGAAGAACGTTTTCCAATGATGAGCAC
TTTTAAAGTTCTGCTATGTGGCGCGGTATTATCCCGTATTGACGCCGGGCAAGAGCAACTCGGTCGCCGCATACACT
ATTCTCAGAATGACTTGGTTGAGTACTCACCAGTCACAGAAAAGCATCTTACGGATGGCATGACAGTAAGAGAATTA
TGCAGTGCTGCCATAACCATGAGTGATAACACTGCGGCCAACTTACTTCTGACAACGATCGGAGGACCGAAGGAGCT
AACCGCTTTTTTGCACAACATGGGGGATCATGTAACTCGCCTTGATCGTTGGGAACCGGAGCTGAATGAAGCCATAC
CAAACGACGAGCGTGACACCACGATGCCTGTAGCAATGGCAACAACGTTGCGCAAACTATTAACTGGCGAACTACTT
ACTCTAGCTTCCCGGCAACAATTAATAGACTGGATGGAGGCGGATAAAGTTGCAGGACCACTTCTGCGCTCGGCCCT
TCCGGCTGGCTGGTTTATTGCTGATAAATCTGGAGCCGGTGAGCGTGGAAGCCGCGGTATCATTGCAGCACTGGGGC
CAGATGGTAAGCCCTCCCGTATCGTAGTTATCTACACGACGGGGAGTCAGGCAACTATGGATGAACGAAATAGACAG
ATCGCTGAGATAGGTGCCTCACTGATTAAGCATTGGTAACTGTCAGACCAAGTTTACTCATATATACTTTAGATTGA
TTTAAAACTTCATTTTTAATTTAAAAGGATCTAGGTGAAGATCCTTTTTGATAATCTCATGACCAAAATCCCTTAAC
GTGAGTTTTCGTTCCACTGAGCGTCAGACCCCGTAGAAAAGATCAAAGGATCTTC

**>pET-CasX2-twst**

TCAGAGGTTTTCACCGTCATCACCGAAACGCGCGAGGCAGCTGCGGTAAAGCTCATCAGCGTGGTCGTGAAGCGATT
CACAGATGTCTGCCTGTTCATCCGCGTCCAGCTCGTTGAGTTTCTCCAGAAGCGTTAATGTCTGGCTTCTGATAAAG
CGGGCCATGTTAAGGGCGGTTTTTTCCTGTTTGGTCACTGATGCCTCCGTGTAAGGGGGATTTCTGTTCATGGGGGT
AATGATACCGATGAAACGAGAGAGGATGCTCACGATACGGGTTACTGATGATGAACATGCCCGGTTACTGGAACGTT
GTGAGGGTAAACAACTGGCGGTATGGATGCGGCGGGACCAGAGAAAAATCACTCAGGGTCAATGCCAGCGCTTCGTT
AATACAGATGTAGGTGTTCCACAGGGTAGCCAGCAGCATCCTGCGATGCAGATCCGGAACATAATGGTGCAGGGCGC
TGACTTCCGCGTTTCCAGACTTTACGAAACACGGAAACCGAAGACCATTCATGTTGTTGCTCAGGTCGCAGACGTTT
TGCAGCAGCAGTCGCTTCACGTTCGCTCGCGTATCGGTGATTCATTCTGCTAACCAGTAAGGCAACCCCGCCAGCCT
AGCCGGGTCCTCAACGACAGGAGCACGATCATGCGCACCCGTGGCCAGGACCCAACGCTGCCCGAGATCTCGATCCC
GCGAAATTAATACGACTCACTATAGGGAGACCACAACGGTTTCCCTCTAGAAATAATTTTGTTTAACTTTAAGAAGG
AGATATACATATGCCGAAGAAAAAGCGTAAGGTTATGCAAGAAATTAAACGTATTAACAAGATCCGCCGTCGTCTGG
TCAAAGACAGCAATACGAAAAAAGCCGGCAAAACCGGTCCGATGAAAACCCTGCTGGTTCGCGTGATGACGCCGGAT
TTGCGTGAGCGCCTGGAAAATCTGCGTAAGAAACCGGAGAATATTCCGCAACCGATCAGCAATACCAGCAGAGCCAA
TCTGAATAAGCTGCTGACGGATTACACCGAGATGAAAAAAGCCATCCTGCATGTGTATTGGGAAGAATTTCAGAAAG

ACCCGGTGGGTCTGATGAGCCGCGTTGCGCAGCCGGCACCGAAGAACATTGACCAGCGTAAGCTGATTCCGGTTAAA
GACGGCAATGAACGTCTGACCAGCAGCGGTTTCGCGTGTTCACAGTGTTGCCAGCCACTGTACGTGTATAAACTGGA
ACAAGTTAATGATAAGGGTAAGCCGCACACGAACTACTTCGGTCGTTGCAACGTGAGCGAGCATGAGCGTTTGATCT
TGCTGTCGCCTCATAAGCCAGAGGCGAACGACGAACTGGTCACCTACAGCCTGGGTAAATTCGGCCAGCGCGCACTG
GATTTCTACAGCATCCACGTGACCCGTGAGAGCAACCATCCGGTCAAGCCGCTGGAGCAAATCGGTGGCAATAGCTG
TGCATCCGGTCCGGTGGGTAAGGCACTGTCCGATGCATGCATGGGTGCCGTGGCTAGCTTCCTGACCAAGTATCAAG
ACATCATTTTAGAGCACCAAAAAGTCATTAAGAAAAATGAAAAGCGCCTGGCGAACCTGAAAGACATCGCGAGCGCA
AATGGTCTGGCGTTCCCGAAGATTACGCTGCCGCCGCAGCCGCACACTAAGGAAGGCATTGAAGCGTACAATAACGT
TGTGGCGCAGATTGTGATCTGGGTTAACCTGAATCTGTGGCAGAAATTGAAGATTGGTCGTGATGAAGCGAAGCCGC
TGCAGCGTCTGAAGGGTTTCCCGAGCTTCCCTCTGGTTGAACGCCAAGCTAATGAAGTCGACTGGTGGGACATGGTG
TGTAACGTTAAAAAGTTGATCAACGAAAAAAAAGAGGATGGTAAGGTTTTCTGGCAAAACTTGGCCGGTTACAAGCG
TCAAGAAGCGCTGCTGCCGTACCTGAGCTCCGAAGAGGATCGTAAAAAGGGTAAGAAATTTGCACGTTATCAATTCG
GCGACCTGTTGCTGCACTTGGAGAAGAAACACGGCGAGGACTGGGGTAAAGTCTATGATGAAGCGTGGGAGCGTATC
GACAAAAAAGTCGAGGGCCTGAGCAAGCATATTAAACTGGAAGAGGAACGCCGTAGCGAAGATGCACAGAGCAAAGC
CGCACTGACGGATTGGCTGCGCGCTAAAGCGAGCTTTGTGATCGAAGGCCTGAAAGAAGCGGATAAAGACGAATTCT
GTCGCTGCGAACTGAAACTGCAAAAATGGTACGGTGATCTGCGCGGTAAGCCGTTTGCGATTGAAGCGGAGAACTCT
ATTCTGGATATCAGCGGCTTTAGCAAACAGTACAATTGTGCGTTCATTTGGCAAAAGGACGGTGTTAAAAAGCTGAA
TCTGTACCTGATCATTAACTATTTCAAGGGTGGCAAGTTGCGTTTTAAGAAAATCAAGCCTGAGGCTTTCGAGGCAA
ATCGTTTCTACACTGTCATTAACAAGAAATCTGGCGAGATCGTGCCGATGGAAGTCAATTTCAACTTTGATGACCCG
AATCTGATCATTCTGCCGTTGGCATTTGGTAAGCGTCAGGGTCGTGAGTTTATCTGGAACGATCTTCTGAGCCTGGA
AACCGGCTCGCTGAAACTGGCCAACGGTCGTGTGATTGAGAAAACGTTATACAACCGCCGTACGCGTCAAGACGAGC
CAGCGCTGTTTGTAGCCCTGACCTTCGAGCGTCGTGAAGTCCTGGATAGCTCGAATATCAAGCCGATGAATCTGATT
GGCATTGATCGTGGTGAGAACATTCCAGCGGTTATCGCGCTGACCGACCCTGAGGGTTGCCCGCTGAGCCGCTTTAA
AGACTCCCTTGGTAATCCGACCCACATCCTGCGCATCGGCGAGTCCTATAAAGAAAAACAACGTACCATCCAGGCCG
CAAAAGAGGTCGAGCAACGTCGTGCGGGCGGCTACAGCCGTAAATATGCAAGCAAAGCCAAGAACCTGGCGGACGAC
ATGGTGAGAAACACTGCCCGCGATCTGCTGTACTATGCTGTCACCCAAGACGCGATGCTGATCTTCGAGAACCTGTC
CCGCGGTTTCGGTCGCCAGGGTAAACGTACCTTTATGGCAGAGCGTCAATATACCCGTATGGAGGACTGGCTGACCG
CGAAATTGGCCTACGAGGGTCTGCCGTCCAAAACCTATCTGAGCAAGACTCTGGCACAGTATACCAGCAAAACGTGC
AGCAATTGCGGTTTTACCATTACGAGCGCGGACTACGATCGCGTCCTGGAGAAATTGAAAAAGACCGCTACGGGTTG
GATGACCACTATTAACGGTAAAGAATTAAAAGTTGAGGGCCAGATCACGTACTATAACCGCTATAAGCGTCAAAACG
TTGTTAAAGACCTGAGCGTTGAGCTGGACCGTCTGAGCGAAGAGAGCGTGAATAACGACATTTCCAGCTGGACCAAG
GGCCGTTCTGGTGAGGCACTTAGCCTCCTCAAAAAGCGTTTTAGCCACCGTCCGGTGCAGGAAAAGTTTGTGTGTCT
GAACTGCGGCTTTGAAACGCACGCCGACGAACAGGCCGCGCTGAACATTGCGCGCAGCTGGCTGTTCCTGCGCAGCC
AAGAGTACAAAAAGTACCAGACGAACAAGACCACGGGTAACACCGATAAGCGCGCTTTCGTGGAAACCTGGCAGAGC
TTTTACCGCAAAAAGTTGAAAGAAGTCTGGAAACCAGCAGTGAATGCACTGAAGCGTCCGGCGGCCACCAAAAAAGC
TGGCCAGGCAAAGAAAAAGAAAGGTTCTTACCCGTATGATGTTCCGGACTACGCCTATCCGTATGACGTTCCGGATT
ACGCGTATCCGTACGACGTCCCGGATTATGCGAGCGAAAATCTTTATTTTCAAGGTAGCGCATGGAGTCATCCTCAA
TTCGAGAAAGGTGGAGGTTCTGGCGGTGGATCGGGAGGTTCAGCGTGGAGCCACCCGCAGTTCGAAAAAGGAAGGGG
ATCCGGCTGCTAACAAAGCCCGAAAGGAAGCTGAGTTGGCTGCTGCCACCGCTGAGCAATAACTAGCATAACCCCTT
GGGGCCTCTAAACGGGTCTTGAGGGGTTTTTTGCTGAAAGGAGGAACTATATCCGGATTGGCGAATGGGACGCGCCC
TGTAGCGGCGCATTAAGCGCGGCGGGTGTGGTGGTTACGCGCAGCGTGACCGCTACACTTGCCAGCGCCCTAGCGCC
CGCTCCTTTCGCTTTCTTCCCTTCCTTTCTCGCCACGTTCGCCGGCTTTCCCCGTCAAGCTCTAAATCGGGGGCTCC
CTTTAGGGTTCCGATTTAGTGCTTTACGGCACCTCGACCCCAAAAAACTTGATTAGGGTGATGGTTCACGTAGTGGG
CCATCGCCCTGATAGACGGTTTTTCGCCCTTTGACGTTGGAGTCCACGTTCTTTAATAGTGGACTCTTGTTCCAAAC
TGGAACAACACTCAACCCTATCTCGGTCTATTCTTTTGATTTATAAGGGATTTTGCCGATTTCGGCCTATTGGTTAA
AAAATGAGCTGATTTAACAAAAATTTAACGCGAATTTTAACAAAATATTAACGTTTACAATTTCAGGTGGCACTTTT
CGGGGAAATGTGCGCGGAACCCCTATTTGTTTATTTTTCTAAATACATTCAAATATGTATCCGCTCATGAGACAATA
ACCCTGATAAATGCTTCAATAATATTGAAAAAGGAAGAGTATGAGTATTCAACATTTCCGTGTCGCCCTTATTCCCT
TTTTTGCGGCATTTTGCCTTCCTGTTTTTGCTCACCCAGAAACGCTGGTGAAAGTAAAAGATGCTGAAGATCAGTTG

GGTGCACGAGTGGGTTACATCGAACTGGATCTCAACAGCGGTAAGATCCTTGAGAGTTTTCGCCCCGAAGAACGTTT
TCCAATGATGAGCACTTTTAAAGTTCTGCTATGTGGCGCGGTATTATCCCGTATTGACGCCGGGCAAGAGCAACTCG
GTCGCCGCATACACTATTCTCAGAATGACTTGGTTGAGTACTCACCAGTCACAGAAAAGCATCTTACGGATGGCATG
ACAGTAAGAGAATTATGCAGTGCTGCCATAACCATGAGTGATAACACTGCGGCCAACTTACTTCTGACAACGATCGG
AGGACCGAAGGAGCTAACCGCTTTTTTGCACAACATGGGGGATCATGTAACTCGCCTTGATCGTTGGGAACCGGAGC
TGAATGAAGCCATACCAAACGACGAGCGTGACACCACGATGCCTGCAGCAATGGCAACAACGTTGCGCAAACTATTA
ACTGGCGAACTACTTACTCTAGCTTCCCGGCAACAATTAATAGACTGGATGGAGGCGGATAAAGTTGCAGGACCACT
TCTGCGCTCGGCCCTTCCGGCTGGCTGGTTTATTGCTGATAAATCTGGAGCCGGTGAGCGTGGGTCTCGCGGTATCA
TTGCAGCACTGGGGCCAGATGGTAAGCCCTCCCGTATCGTAGTTATCTACACGACGGGGAGTCAGGCAACTATGGAT
GAACGAAATAGACAGATCGCTGAGATAGGTGCCTCACTGATTAAGCATTGGTAACTGTCAGACCAAGTTTACTCATA
TATACTTTAGATTGATTTAAAACTTCATTTTTAATTTAAAAGGATCTAGGTGAAGATCCTTTTTGATAATCTCATGA
CCAAAATCCCTTAACGTGAGTTTTCGTTCCACTGAGCGTCAGACCCCGTAGAAAAGATCAAAGGATCTTCTTGAGAT
CCTTTTTTTCTGCGCGTAATCTGCTGCTTGCAAACAAAAAAACCACCGCTACCAGCGGTGGTTTGTTTGCCGGATCA
AGAGCTACCAACTCTTTTTCCGAAGGTAACTGGCTTCAGCAGAGCGCAGATACCAAATACTGTCCTTCTAGTGTAGC
CGTAGTTAGGCCACCACTTCAAGAACTCTGTAGCACCGCCTACATACCTCGCTCTGCTAATCCTGTTACCAGTGGCT
GCTGCCAGTGGCGATAAGTCGTGTCTTACCGGGTTGGACTCAAGACGATAGTTACCGGATAAGGCGCAGCGGTCGGG
CTGAACGGGGGGTTCGTGCACACAGCCCAGCTTGGAGCGAACGACCTACACCGAACTGAGATACCTACAGCGTGAGC
TATGAGAAAGCGCCACGCTTCCCGAAGGGAGAAAGGCGGACAGGTATCCGGTAAGCGGCAGGGTCGGAACAGGAGAG
CGCACGAGGGAGCTTCCAGGGGGAAACGCCTGGTATCTTTATAGTCCTGTCGGGTTTCGCCACCTCTGACTTGAGCG
TCGATTTTTGTGATGCTCGTCAGGGGGGCGGAGCCTATGGAAAAACGCCAGCAACGCGGCCTTTTTACGGTTCCTGG
CCTTTTGCTGGCCTTTTGCTCACATGTTCTTTCCTGCGTTATCCCCTGATTCTGTGGATAACCGTATTACCGCCTTT
GAGTGAGCTGATACCGCTCGCCGCAGCCGAACGACCGAGCGCAGCGAGTCAGTGAGCGAGGAAGCGGAAGAGCGCCT
GATGCGGTATTTTCTCCTTACGCATCTGTGCGGTATTTCACACCGCATATATGGTGCACTCTCAGTACAATCTGCTC
TGATGCCGCATAGTTAAGCCAGTATACACTCCGCTATCGCTACGTGACTGGGTCATGGCTGCGCCCCGACACCCGCC
AACACCCGCTGACGCGCCCTGACGGGCTTGTCTGCTCCCGGCATCCGCTTACAGACAAGCTGTGACCGTCTCCGGGA
GCTGCATGTG

**>pET-CasX2Max-twst**

TCAGAGGTTTTCACCGTCATCACCGAAACGCGCGAGGCAGCTGCGGTAAAGCTCATCAGCGTGGTCGTGAAGCGATT
CACAGATGTCTGCCTGTTCATCCGCGTCCAGCTCGTTGAGTTTCTCCAGAAGCGTTAATGTCTGGCTTCTGATAAAG
CGGGCCATGTTAAGGGCGGTTTTTTCCTGTTTGGTCACTGATGCCTCCGTGTAAGGGGGATTTCTGTTCATGGGGGT
AATGATACCGATGAAACGAGAGAGGATGCTCACGATACGGGTTACTGATGATGAACATGCCCGGTTACTGGAACGTT
GTGAGGGTAAACAACTGGCGGTATGGATGCGGCGGGACCAGAGAAAAATCACTCAGGGTCAATGCCAGCGCTTCGTT
AATACAGATGTAGGTGTTCCACAGGGTAGCCAGCAGCATCCTGCGATGCAGATCCGGAACATAATGGTGCAGGGCGC
TGACTTCCGCGTTTCCAGACTTTACGAAACACGGAAACCGAAGACCATTCATGTTGTTGCTCAGGTCGCAGACGTTT
TGCAGCAGCAGTCGCTTCACGTTCGCTCGCGTATCGGTGATTCATTCTGCTAACCAGTAAGGCAACCCCGCCAGCCT
AGCCGGGTCCTCAACGACAGGAGCACGATCATGCGCACCCGTGGCCAGGACCCAACGCTGCCCGAGATCTCGATCCC
GCGAAATTAATACGACTCACTATAGGGAGACCACAACGGTTTCCCTCTAGAAATAATTTTGTTTAACTTTAAGAAGG
AGATATACATATGCCGAAGAAAAAGCGTAAGGTTATGCAAGAAATTAAACGTATTAACAAGATCCGCCGTCGTCTGG
TCAAAGACAGCAATACGAAAAAAGCCGGCAAACGCGGTCCGATGAAAACCCTGCTGGTTCGCGTGATGACGCCGGAT
TTGCGTGAGCGCCTGGAAAATCTGCGTAAGAAACCGGAGAATATTCCGCAACCGATCAGCAATACCAGCAGAGCCAA
TCTGAATAAGCTGCTGACGGATTACACCGAGATGAAAAAAGCCATCCTGCATGTGTATTGGGAAGAATTTCAGAAAG
ACCCGGTGGGTCTGATGAGCCGCGTTGCGCAGCCGGCACCGAAGAACATTGACCAGCGTAAGCTGATTCCGGTTAAA
GACGGCAATGAACGTCTGACCAGCAGCGGTTTCGCGTGTTCACAGTGTTGCCAGCCACTGTACGTGTATAAACTGGA
ACAAGTTAATGATAAGGGTAAGCCGCACACGAACTACTTCGGTCGTTGCAACGTGAGCGAGCATGAGCGTTTGATCT
TGCTGTCGCCTCATAAGCCAGAGGCGAACGACGAACTGGTCACCTACAGCCTGGGTAAATTCGGCCAGCGCGCACTG
GATTTCTACAGCATCCACGTGACCCGTGAGAGCAACCATCCGGTCAAGCCGCTGGAGCAAATCGGTGGCAATAGCTG

TGCATCCGGTCCGGTGGGTAAGGCACTGTCCGATGCATGCATGGGTGCCGTGGCTAGCTTCCTGACCAAGTATCAAG
ACATCATTTTAGAGCACCAAAAAGTCATTAAGAAAAATGAAAAGCGCCTGGCGAACCTGAAAGACATCGCGAGCGCA
AATGGTCTGGCGTTCCCGAAGATTACGCTGCCGCCGCAGCCGCACACTAAGGAAGGCATTGAAGCGTACAATAACGT
TGTGGCGCAGATTGTGATCTGGGTTAACCTGAATCTGTGGCAGAAATTGAAGATTGGTCGTGATGAAGCGAAGCCGC
TGCAGCGTCTGAAGGGTTTCCCGAGCTTCCCTCTGGTTGAACGCCAAGCTAATGAAGTCGACTGGTGGGACATGGTG
TGTAACGTTAAAAAGTTGATCAACGAAAAAAAAGAGGATGGTAAGGTTTTCTGGCAAAACTTGGCCGGTTACAAGCG
TCAAGAAGCGCTGCTGCCGTACCTGAGCTCCGAAGAGGATCGTAAAAAGGGTAAGAAATTTGCACGTTATCAATTCG
GCGACCTGTTGCTGCACTTGGAGAAGAAACACGGCGAGGACTGGGGTAAAGTCTATGATGAAGCGTGGGAGCGTATC
GACAAAAAAGTCGAGGGCCTGAGCAAGCATATTAAACTGGAAGAGGAACGCCGTAGCGAAGATGCACAGAGCAAAGC
CGCACTGACGGATTGGCTGCGCGCTAAAGCGAGCTTTGTGATCGAAGGCCTGAAAGAAGCGGATAAAGACGAATTCT
GTCGCTGCGAACTGAAACTGCAAAAATGGTACGGTGATCTGCGCGGTAAGCCGTTTGCGATTGAAGCGGAGAACTCT
ATTCTGGATATCAGCGGCTTTAGCAAACAGTACAATTGTGCGTTCATTTGGCAAAAGGACGGTGTTAAAAAGCTGAA
TCTGTACCTGATCATTAACTATTTCAAGGGTGGCAAGTTGCGTTTTAAGAAAATCAAGCCTGAGGCTTTCGAGGCAA
ATCGTTTCTACACTGTCATTAACAAGAAATCTGGCGAGATCGTGCCGATGGAAGTCAATTTCAACTTTGATGACCCG
AATCTGATCATTCTGCCGTTGGCATTTGGTAAGCGTCAGGGTCGTGAGTTTATCTGGAACGATCTTCTGAGCCTGGA
AACCGGCTCGCTGCGCCTGGCCAACGGTCGTGTGATTGAGAAAACGTTATACAACCGCCGTACGCGTCAAGACGAGC
CAGCGCTGTTTGTAGCCCTGACCTTCGAGCGTCGTGAAGTCCTGGATAGCTCGAATATCAAGCCGATGAATCTGATT
GGCATTGATCGTGGTGAGAACATTCCAGCGGTTATCGCGCTGACCGACCCTGAGGGTTGCCCGCTGAGCCGCTTTAA
AGACTCCCTTGGTAATCCGACCCACATCCTGCGCATCGGCGAGTCCTATAAAGAAAAACAACGTACCATCCAGGCCG
CAAAAGAGGTCGAGCAACGTCGTGCGGGCGGCTACAGCCGTAAATATGCAAGCAAAGCCAAGAACCTGGCGGACGAC
ATGGTGAGAAACACTGCCCGCGATCTGCTGTACTATGCTGTCACCCAAGACGCGATGCTGATCTTCGAGAACCTGTC
CCGCGGTTTCGGTCGCCAGGGTAAACGTACCTTTATGGCAGAGCGTCAATATACCCGTATGGAGGACTGGCTGACCG
CGAAATTGGCCTACGAGGGTCTGCCGTCCAAAACCTATCTGAGCAAGACTCTGGCACAGTATACCAGCCGCACGTGC
AGCAATTGCGGTTTTACCATTACGAGCGCGGACTACGATCGCGTCCTGGAGAAATTGAAAAAGACCGCTACGGGTTG
GATGACCACTATTAACGGTAAAGAATTAAAAGTTGAGGGCCAGATCACGTACTATAACCGCTATAAGCGTCAAAACG
TTGTTAAAGACCTGAGCGTTGAGCTGGACCGTCTGAGCGAAGAGAGCGTGAATAACGACATTTCCAGCTGGACCAAG
GGCCGTTCTGGTGAGGCACTTAGCCTCCTCAAAAAGCGTTTTAGCCACCGTCCGGTGCAGGAAAAGTTTGTGTGTCT
GAACTGCGGCTTTGAAACGCACGCCGACGAACAGGCCGCGCTGAACATTGCGCGCAGCTGGCTGTTCCTGCGCAGCC
AAGAGTACAAAAAGTACCAGACGAACAAGACCACGGGTAACACCGATAAGCGCGCTTTCGTGGAAACCTGGCAGAGC
TTTTACCGCAAAAAGTTGAAAGAAGTCTGGAAACCAGCAGTGAATGCACTGAAGCGTCCGGCGGCCACCAAAAAAGC
TGGCCAGGCAAAGAAAAAGAAAGGTTCTTACCCGTATGATGTTCCGGACTACGCCTATCCGTATGACGTTCCGGATT
ACGCGTATCCGTACGACGTCCCGGATTATGCGAGCGAAAATCTTTATTTTCAAGGTAGCGCATGGAGTCATCCTCAA
TTCGAGAAAGGTGGAGGTTCTGGCGGTGGATCGGGAGGTTCAGCGTGGAGCCACCCGCAGTTCGAAAAAGGAAGGGG
ATCCGGCTGCTAACAAAGCCCGAAAGGAAGCTGAGTTGGCTGCTGCCACCGCTGAGCAATAACTAGCATAACCCCTT
GGGGCCTCTAAACGGGTCTTGAGGGGTTTTTTGCTGAAAGGAGGAACTATATCCGGATTGGCGAATGGGACGCGCCC
TGTAGCGGCGCATTAAGCGCGGCGGGTGTGGTGGTTACGCGCAGCGTGACCGCTACACTTGCCAGCGCCCTAGCGCC
CGCTCCTTTCGCTTTCTTCCCTTCCTTTCTCGCCACGTTCGCCGGCTTTCCCCGTCAAGCTCTAAATCGGGGGCTCC
CTTTAGGGTTCCGATTTAGTGCTTTACGGCACCTCGACCCCAAAAAACTTGATTAGGGTGATGGTTCACGTAGTGGG
CCATCGCCCTGATAGACGGTTTTTCGCCCTTTGACGTTGGAGTCCACGTTCTTTAATAGTGGACTCTTGTTCCAAAC
TGGAACAACACTCAACCCTATCTCGGTCTATTCTTTTGATTTATAAGGGATTTTGCCGATTTCGGCCTATTGGTTAA
AAAATGAGCTGATTTAACAAAAATTTAACGCGAATTTTAACAAAATATTAACGTTTACAATTTCAGGTGGCACTTTT
CGGGGAAATGTGCGCGGAACCCCTATTTGTTTATTTTTCTAAATACATTCAAATATGTATCCGCTCATGAGACAATA
ACCCTGATAAATGCTTCAATAATATTGAAAAAGGAAGAGTATGAGTATTCAACATTTCCGTGTCGCCCTTATTCCCT
TTTTTGCGGCATTTTGCCTTCCTGTTTTTGCTCACCCAGAAACGCTGGTGAAAGTAAAAGATGCTGAAGATCAGTTG
GGTGCACGAGTGGGTTACATCGAACTGGATCTCAACAGCGGTAAGATCCTTGAGAGTTTTCGCCCCGAAGAACGTTT
TCCAATGATGAGCACTTTTAAAGTTCTGCTATGTGGCGCGGTATTATCCCGTATTGACGCCGGGCAAGAGCAACTCG
GTCGCCGCATACACTATTCTCAGAATGACTTGGTTGAGTACTCACCAGTCACAGAAAAGCATCTTACGGATGGCATG
ACAGTAAGAGAATTATGCAGTGCTGCCATAACCATGAGTGATAACACTGCGGCCAACTTACTTCTGACAACGATCGG
AGGACCGAAGGAGCTAACCGCTTTTTTGCACAACATGGGGGATCATGTAACTCGCCTTGATCGTTGGGAACCGGAGC

TGAATGAAGCCATACCAAACGACGAGCGTGACACCACGATGCCTGCAGCAATGGCAACAACGTTGCGCAAACTATTA
ACTGGCGAACTACTTACTCTAGCTTCCCGGCAACAATTAATAGACTGGATGGAGGCGGATAAAGTTGCAGGACCACT
TCTGCGCTCGGCCCTTCCGGCTGGCTGGTTTATTGCTGATAAATCTGGAGCCGGTGAGCGTGGGTCTCGCGGTATCA
TTGCAGCACTGGGGCCAGATGGTAAGCCCTCCCGTATCGTAGTTATCTACACGACGGGGAGTCAGGCAACTATGGAT
GAACGAAATAGACAGATCGCTGAGATAGGTGCCTCACTGATTAAGCATTGGTAACTGTCAGACCAAGTTTACTCATA
TATACTTTAGATTGATTTAAAACTTCATTTTTAATTTAAAAGGATCTAGGTGAAGATCCTTTTTGATAATCTCATGA
CCAAAATCCCTTAACGTGAGTTTTCGTTCCACTGAGCGTCAGACCCCGTAGAAAAGATCAAAGGATCTTCTTGAGAT
CCTTTTTTTCTGCGCGTAATCTGCTGCTTGCAAACAAAAAAACCACCGCTACCAGCGGTGGTTTGTTTGCCGGATCA
AGAGCTACCAACTCTTTTTCCGAAGGTAACTGGCTTCAGCAGAGCGCAGATACCAAATACTGTCCTTCTAGTGTAGC
CGTAGTTAGGCCACCACTTCAAGAACTCTGTAGCACCGCCTACATACCTCGCTCTGCTAATCCTGTTACCAGTGGCT
GCTGCCAGTGGCGATAAGTCGTGTCTTACCGGGTTGGACTCAAGACGATAGTTACCGGATAAGGCGCAGCGGTCGGG
CTGAACGGGGGGTTCGTGCACACAGCCCAGCTTGGAGCGAACGACCTACACCGAACTGAGATACCTACAGCGTGAGC
TATGAGAAAGCGCCACGCTTCCCGAAGGGAGAAAGGCGGACAGGTATCCGGTAAGCGGCAGGGTCGGAACAGGAGAG
CGCACGAGGGAGCTTCCAGGGGGAAACGCCTGGTATCTTTATAGTCCTGTCGGGTTTCGCCACCTCTGACTTGAGCG
TCGATTTTTGTGATGCTCGTCAGGGGGGCGGAGCCTATGGAAAAACGCCAGCAACGCGGCCTTTTTACGGTTCCTGG
CCTTTTGCTGGCCTTTTGCTCACATGTTCTTTCCTGCGTTATCCCCTGATTCTGTGGATAACCGTATTACCGCCTTT
GAGTGAGCTGATACCGCTCGCCGCAGCCGAACGACCGAGCGCAGCGAGTCAGTGAGCGAGGAAGCGGAAGAGCGCCT
GATGCGGTATTTTCTCCTTACGCATCTGTGCGGTATTTCACACCGCATATATGGTGCACTCTCAGTACAATCTGCTC
TGATGCCGCATAGTTAAGCCAGTATACACTCCGCTATCGCTACGTGACTGGGTCATGGCTGCGCCCCGACACCCGCC
AACACCCGCTGACGCGCCCTGACGGGCTTGTCTGCTCCCGGCATCCGCTTACAGACAAGCTGTGACCGTCTCCGGGA
GCTGCATGTG

**>pBLO 62.5 CasX2Max**

TTGAGATCCTTTTTTTCTGCGCGTAATCTGCTGCTTGCAAACAAAAAAACCACCGCTACCAGCGGTGGTTTGTTTGC
CGGATCAAGAGCTACCAACTCTTTTTCCGAAGGTAACTGGCTTCAGCAGAGCGCAGATACCAAATACTGTCCTTCTA
GTGTAGCCGTAGTTAGGCCACCACTTCAAGAACTCTGTAGCACCGCCTACATACCTCGCTCTGCTAATCCTGTTACC
AGTGGCTGCTGCCAGTGGCGATAAGTCGTGTCTTACCGGGTTGGACTCAAGACGATAGTTACCGGATAAGGCGCAGC
GGTCGGGCTGAACGGGGGGTTCGTGCACACAGCCCAGCTTGGAGCGAACGACCTACACCGAACTGAGATACCTACAG
CGTGAGCTATGAGAAAGCGCCACGCTTCCCGAAGGGAGAAAGGCGGACAGGTATCCGGTAAGCGGCAGGGTCGGAAC
AGGAGAGCGCACGAGGGAGCTTCCAGGGGGAAACGCCTGGTATCTTTATAGTCCTGTCGGGTTTCGCCACCTCTGAC
TTGAGCGTCGATTTTTGTGATGCTCGTCAGGGGGGCGGAGCCTATGGAAAAACGCCAGCAACGCGGCCTTTTTACGG
TTCCTGGCCTTTTGCTGGCCTTTTGCTCACATGTGAGGGCCTATTTCCCATGATTCCTTCATATTTGCATATACGAT
ACAAGGCTGTTAGAGAGATAATTGGAATTAATTTGACTGTAAACACAAAGATATTAGTACAAAATACGTGACGTAGA
AAGTAATAATTTCTTGGGTAGTTTGCAGTTTTAAAATTATGTTTTAAAATGGACTATCATATGCTTACCGTAACTTG
AAAGTATTTCGATTTCTTGGCTTTATATATCTTGTGGAAAGGACGAAACACCACATCTGGCGCGTTTATTCCATTAC
TTTGGAGCCAGTCCCAGCGACTATGTCGTATGGACGAAGCGCTTATTTATCGGAGAGAAACCGATAAGTAAAACGCA
TCAAAGGCGTGTCCGGCGAGGGCGAGTTTTTTTTGGTACCCGTTACATAACTTACGGTAAATGGCCCGCCTGGCTGA
CCGCCCAACGACCCCCGCCCATTGACGTCAATAGTAACGCCAATAGGGACTTTCCATTGACGTCAATGGGTGGAGTA
TTTACGGTAAACTGCCCACTTGGCAGTACATCAAGTGTATCATATGCCAAGTACGCCCCCTATTGACGTCAATGACG
GTAAATGGCCCGCCTGGCATTGTGCCCAGTACATGACCTTATGGGACTTTCCTACTTGGCAGTACATCTACGTATTA
GTCATCGCTATTACCATGGTCGAGGTGAGCCCCACGTTCTGCTTCACTCTCCCCATCTCCCCCCCCTCCCCACCCCC
AATTTTGTATTTATTTATTTTTTAATTATTTTGTGCAGCGATGGGGGCGGGGGGGGGGGGGGGGCGCGCGCCAGGCG
GGGCGGGGCGGGGCGAGGGGCGGGGCGGGGCGAGGCGGAGAGGTGCGGCGGCAGCCAATCAGAGCGGCGCGCTCCGA
AAGTTTCCTTTTATGGCGAGGCGGCGGCGGCGGCGGCCCTATAAAAAGCGAAGCGCGCGGCGGGCGGGAGTCGCTGC
GACGCTGCCTTCGCCCCGTGCCCCGCTCCGCCGCCGCCTCGCGCCGCCCGCCCCGGCTCTGACTGACCGCGTTACTC
CCACAGGTGAGCGGGCGGGACGGCCCTTCTCCTCCGGGCTGTAATTAGCTGAGCAAGAGGTAAGGGTTTAAGGGATG
GTTGGTTGGTGGGGTATTAATGTTTAATTACCTGGAGCACCTGCCTGAAATCACTTTTTTTCAGGTTGGACCGGTGC

CACCATGGCCCCAAAGAAGAAGCGGAAGGTCAGCATGCAAGAGATCAAGAGAATCAACAAGATCAGAAGGAGACTGG
TCAAGGACAGCAACACAAAGAAGGCCGGCAAGCGCGGCCCCATGAAAACCCTGCTCGTCAGAGTGATGACCCCTGAC
CTGAGAGAGCGGCTGGAAAACCTGAGAAAGAAGCCCGAGAACATCCCTCAGCCTATCAGCAACACCAGCAGGGCCAA
CCTGAACAAGCTGCTGACCGACTACACCGAGATGAAGAAAGCCATCCTGCACGTGTACTGGGAAGAGTTCCAGAAAG
ACCCCGTGGGCCTGATGAGCAGAGTTGCTCAGCCCGCTCCTAAGAACATCGACCAGAGAAAGCTGATCCCCGTGAAG
GACGGCAACGAGAGACTGACCTCTAGCGGCTTTGCCTGCAGCCAGTGTTGCCAGCCTCTGTACGTGTACAAGCTGGA
ACAAGTGAACGACAAGGGCAAGCCCCACACCAACTACTTCGGCAGATGCAACGTGTCCGAGCACGAGAGGCTGATCC
TGCTGTCTCCTCACAAGCCCGAGGCCAACGATGAGCTGGTCACATACAGCCTGGGCAAGTTCGGACAGAGAGCCCTG
GACTTCTACAGCATCCACGTGACCAGGGAGAGCAATCACCCTGTGAAGCCCCTGGAACAGATCGGCGGCAATAGCTG
TGCCTCTGGACCTGTGGGAAAAGCCCTGAGCGACGCCTGTATGGGAGCCGTGGCATCCTTCCTGACCAAGTACCAGG
ACATCATCCTGGAACACCAGAAAGTGATCAAGAAGAACGAGAAAAGACTGGCCAACCTCAAGGATATCGCCAGCGCT
AACGGCCTGGCCTTTCCTAAGATCACCCTGCCTCCACAGCCTCACACCAAAGAGGGCATCGAGGCCTACAACAACGT
GGTGGCCCAGATCGTGATTTGGGTCAACCTGAATCTGTGGCAGAAGCTGAAGATCGGCAGGGACGAAGCCAAGCCAC
TGCAGAGACTGAAGGGCTTCCCTAGCTTCCCTCTGGTGGAAAGACAGGCCAATGAAGTGGATTGGTGGGACATGGTC
TGCAACGTGAAGAAGCTGATCAACGAGAAGAAAGAGGATGGCAAGGTTTTCTGGCAGAACCTGGCCGGCTACAAGAG
ACAAGAAGCCCTGCTGCCTTACCTGAGCAGCGAAGAGGACCGGAAGAAGGGCAAGAAGTTCGCCAGATACCAGTTCG
GCGACCTGCTGCTGCACCTGGAAAAGAAGCACGGCGAGGACTGGGGCAAAGTGTACGATGAGGCCTGGGAGAGAATC
GACAAGAAGGTGGAAGGCCTGAGCAAGCACATTAAGCTGGAAGAGGAAAGAAGGAGCGAGGACGCCCAATCTAAAGC
CGCTCTGACCGATTGGCTGAGAGCCAAGGCCAGCTTTGTGATCGAGGGCCTGAAAGAGGCCGACAAGGACGAGTTCT
GCAGATGCGAGCTGAAGCTGCAGAAGTGGTACGGCGATCTGAGAGGCAAGCCCTTCGCCATTGAGGCCGAGAACAGC
ATCCTGGACATCAGCGGCTTCAGCAAGCAGTACAACTGCGCCTTCATTTGGCAGAAAGACGGCGTCAAGAAACTGAA
CCTGTACCTGATCATCAATTACTTCAAAGGCGGCAAGCTGCGGTTCAAGAAGATCAAACCCGAGGCCTTCGAGGCTA
ACAGATTCTACACCGTGATCAACAAAAAGTCCGGCGAGATCGTGCCCATGGAAGTGAACTTCAACTTCGACGACCCC
AACCTGATTATCCTGCCTCTGGCCTTCGGCAAGAGACAGGGCAGAGAGTTCATCTGGAACGATCTGCTGAGCCTGGA
AACCGGCTCTCTGCGCCTGGCCAATGGCAGAGTGATCGAGAAAACCCTGTACAACAGGAGAACCAGACAGGACGAGC
CTGCTCTGTTTGTGGCCCTGACCTTCGAGAGAAGAGAGGTGCTGGACAGCAGCAACATCAAGCCCATGAACCTGATC
GGCATCGACCGGGGCGAGAATATCCCTGCTGTGATCGCCCTGACAGACCCTGAAGGATGCCCACTGAGCAGATTCAA
GGACTCCCTGGGCAACCCTACACACATCCTGAGAATCGGCGAGAGCTACAAAGAGAAGCAGAGGACAATCCAGGCCG
CCAAAGAGGTGGAACAGAGAAGAGCCGGCGGATACTCTAGGAAGTACGCCAGCAAGGCCAAGAATCTGGCCGACGAC
ATGGTCCGAAACACCGCCAGAGATCTGCTGTACTACGCCGTGACACAGGACGCCATGCTGATCTTCGAGAATCTGAG
CAGAGGCTTCGGCCGGCAGGGCAAGAGAACCTTTATGGCCGAGAGGCAGTACACCAGAATGGAAGATTGGCTCACAG
CTAAACTGGCCTACGAGGGACTGCCCAGCAAGACCTACCTGTCCAAAACACTGGCCCAGTATACCTCCCGCACCTGC
AGCAATTGCGGCTTCACCATCACCAGCGCCGACTACGACAGAGTGCTGGAAAAGCTCAAGAAAACCGCCACCGGCTG
GATGACCACCATCAACGGCAAAGAGCTGAAGGTTGAGGGCCAGATCACCTACTACAACAGGTACAAGAGGCAGAACG
TCGTGAAGGATCTGAGCGTGGAACTGGACAGACTGAGCGAAGAGAGCGTGAACAACGACATCAGCAGCTGGACAAAG
GGCAGATCAGGCGAGGCTCTGAGCCTGCTGAAGAAGAGGTTTAGCCACAGACCTGTGCAAGAGAAGTTCGTGTGCCT
GAACTGCGGCTTCGAGACACACGCCGATGAACAGGCTGCCCTGAACATTGCCAGAAGCTGGCTGTTCCTGAGAAGCC
AAGAGTACAAGAAGTACCAGACCAACAAGACCACCGGCAACACCGACAAGAGGGCCTTTGTGGAAACCTGGCAGAGC
TTCTACAGAAAAAAGCTGAAAGAAGTCTGGAAGCCCGCCGTGACTAGTCCAAAAAAGAAGAGAAAGGTAGATTACAA
AGATGACGATGACAAAGACTACAAGGATGATGATGATAAGGGATCCGGCGCAACAAACTTCTCTCTGCTGAAACAAG
CCGGAGATGTCGAAGAGAATCCTGGACCGACCGAGTACAAGCCCACGGTGCGCCTCGCCACCCGCGACGACGTCCCC
AGGGCCGTACGCACCCTCGCCGCCGCGTTCGCCGACTACCCCGCCACGCGCCACACCGTCGATCCGGACCGCCACAT
CGAGCGGGTCACCGAGCTGCAAGAACTCTTCCTCACGCGCGTCGGGCTCGACATCGGCAAGGTGTGGGTCGCGGACG
ACGGCGCCGCGGTGGCGGTCTGGACCACGCCGGAGAGCGTCGAAGCGGGGGCGGTGTTCGCCGAGATCGGCCCGCGC
ATGGCCGAGTTGAGCGGTTCCCGGCTGGCCGCGCAGCAACAGATGGAAGGCCTCCTGGCGCCGCACCGGCCCAAGGA
GCCCGCGTGGTTCCTGGCCACCGTCGGAGTCTCGCCCGACCACCAGGGCAAGGGTCTGGGCAGCGCCGTCGTGCTCC
CCGGAGTGGAGGCGGCCGAGCGCGCCGGGGTGCCCGCCTTCCTGGAGACCTCCGCGCCCCGCAACCTCCCCTTCTAC
GAGCGGCTCGGCTTCACCGTCACCGCCGACGTCGAGGTGCCCGAAGGACCGCGCACCTGGTGCATGACCCGCAAGCC
CGGTGCCTGAACGCGTTAAGAATTCCTAGAGCTCGCTGATCAGCCTCGACTGTGCCTTCTAGTTGCCAGCCATCTGT

TGTTTGCCCCTCCCCCGTGCCTTCCTTGACCCTGGAAGGTGCCACTCCCACTGTCCTTTCCTAATAAAATGAGGAAA
TTGCATCGCATTGTCTGAGTAGGTGTCATTCTATTCTGGGGGGTGGGGTGGGGCAGGACAGCAAGGGGGAGGATTGG
GAAGAGAATAGCAGGCATGCTGGGGAGCGGCCGCAGGAACCCCTAGTGATGGAGTTGGCCACTCCCTCTCTGCGCGC
TCGCTCGCTCACTGAGGCCGGGCGACCAAAGGTCGCCCGACGCCCGGGCTTTGCCCGGGCGGCCTCAGTGAGCGAGC
GAGCGCGCAGCTGCCTGCAGGGGCGCCTGATGCGGTATTTTCTCCTTACGCATCTGTGCGGTATTTCACACCGCATA
CGTCAAAGCAACCATAGTACGCGCCCTGTAGCGGCGCATTAAGCGCGGCGGGTGTGGTGGTTACGCGCAGCGTGACC
GCTACACTTGCCAGCGCCCTAGCGCCCGCTCCTTTCGCTTTCTTCCCTTCCTTTCTCGCCACGTTCGCCGGCTTTCC
CCGTCAAGCTCTAAATCGGGGGCTCCCTTTAGGGTTCCGATTTAGTGCTTTACGGCACCTCGACCCCAAAAAACTTG
ATTTGGGTGATGGTTCACGTAGTGGGCCATCGCCCTGATAGACGGTTTTTCGCCCTTTGACGTTGGAGTCCACGTTC
TTTAATAGTGGACTCTTGTTCCAAACTGGAACAACACTCAACCCTATCTCGGGCTATTCTTTTGATTTATAAGGGAT
TTTGCCGATTTCGGCCTATTGGTTAAAAAATGAGCTGATTTAACAAAAATTTAACGCGAATTTTAACAAAATATTAA
CGTTTACAATTTTATGGTGCACTCTCAGTACAATCTGCTCTGATGCCGCATAGTTAAGCCAGCCCCGACACCCGCCA
ACACCCGCTGACGCGCCCTGACGGGCTTGTCTGCTCCCGGCATCCGCTTACAGACAAGCTGTGACCGTCTCCGGGAG
CTGCATGTGTCAGAGGTTTTCACCGTCATCACCGAAACGCGCGAGACGAAAGGGCCTCGTGATACGCCTATTTTTAT
AGGTTAATGTCATGATAATAATGGTTTCTTAGACGTCAGGTGGCACTTTTCGGGGAAATGTGCGCGGAACCCCTATT
TGTTTATTTTTCTAAATACATTCAAATATGTATCCGCTCATGAGACAATAACCCTGATAAATGCTTCAATAATATTG
AAAAAGGAAGAGTATGAGTATTCAACATTTCCGTGTCGCCCTTATTCCCTTTTTTGCGGCATTTTGCCTTCCTGTTT
TTGCTCACCCAGAAACGCTGGTGAAAGTAAAAGATGCTGAAGATCAGTTGGGTGCACGAGTGGGTTACATCGAACTG
GATCTCAACAGCGGTAAGATCCTTGAGAGTTTTCGCCCCGAAGAACGTTTTCCAATGATGAGCACTTTTAAAGTTCT
GCTATGTGGCGCGGTATTATCCCGTATTGACGCCGGGCAAGAGCAACTCGGTCGCCGCATACACTATTCTCAGAATG
ACTTGGTTGAGTACTCACCAGTCACAGAAAAGCATCTTACGGATGGCATGACAGTAAGAGAATTATGCAGTGCTGCC
ATAACCATGAGTGATAACACTGCGGCCAACTTACTTCTGACAACGATCGGAGGACCGAAGGAGCTAACCGCTTTTTT
GCACAACATGGGGGATCATGTAACTCGCCTTGATCGTTGGGAACCGGAGCTGAATGAAGCCATACCAAACGACGAGC
GTGACACCACGATGCCTGTAGCAATGGCAACAACGTTGCGCAAACTATTAACTGGCGAACTACTTACTCTAGCTTCC
CGGCAACAATTAATAGACTGGATGGAGGCGGATAAAGTTGCAGGACCACTTCTGCGCTCGGCCCTTCCGGCTGGCTG
GTTTATTGCTGATAAATCTGGAGCCGGTGAGCGTGGAAGCCGCGGTATCATTGCAGCACTGGGGCCAGATGGTAAGC
CCTCCCGTATCGTAGTTATCTACACGACGGGGAGTCAGGCAACTATGGATGAACGAAATAGACAGATCGCTGAGATA
GGTGCCTCACTGATTAAGCATTGGTAACTGTCAGACCAAGTTTACTCATATATACTTTAGATTGATTTAAAACTTCA
TTTTTAATTTAAAAGGATCTAGGTGAAGATCCTTTTTGATAATCTCATGACCAAAATCCCTTAACGTGAGTTTTCGT
TCCACTGAGCGTCAGACCCCGTAGAAAAGATCAAAGGATCTTC
