## Supplementary material for "ENHANCED CLEAVAGE OF GENOMIC *CCR5* USING CASX2^Max^": Supplmental Table S4

### Supplemental Table S4

#### Guide Cloning Oligonucleotides

| CasX2_sgRNA | forward primer | Reverse primer |
|---|---|---|
| sg5 (17 nt), for plasmids with one guide site | CAAAGAAAGTCCCACTGGGCGGTTTTT | CGGCAAAAACCGCCCAGTGGGACTTTC |
| sg5 (20 nt), for plasmids with one guide site | CAAAGAAAGTCCCACTGGGCGGCAGTTTTT | CGGCAAAAACTGCCGCCCAGTGGGACTTTC |
| sg5 (23 nt), for plasmids with one guide site | CAAAGAAAGTCCCACTGGGCGGCAGCATTTTTT | CGGCAAAAAATGCTGCCGCCCAGTGGGACTTTC |
| sg5 (23 nt), for plasmids with two guide sites (site A) | CATCAAAGAAAGTCCCACTGGGCGGCAGCAT | AAAAATGCTGCCGCCCAGTGGGACTTTCTTT |
| sg10 (17 nt), for plasmids with one guide site | CAAAGTGCTCCCCAGTGGATCGTTTTT | CGGCAAAAACGATCCACTGGGGAGCAC |
| sg10 (20 nt), for plasmids with one guide site | CAAAGTGCTCCCCAGTGGATCGGGTTTTTT | CGGCAAAAAACCCGATCCACTGGGGAGCAC |
| sg10 (23 nt), for plasmids with one guide site | CAAAGTGCTCCCCAGTGGATCGGGTGTATTTTT | CGGCAAAAATACACCCGATCCACTGGGGAGCAC |

| SaCas9_sgRNA | forward primer | Reverse primer |
|---|---|---|
| CCR5-A (20 nt), for plasmids with one guide site | GAAACACCGCGGCAGCATAGTGAGCCCAGGTTT | ACTAAAACCTGGGCTCACTATGCTGCCGCGGTG |
| CCR5-B (20 nt), for plasmids with one guide site | GAAACACCGTCAGTTTACACCCGATCCACGTTT | ACTAAAACGTGGATCGGGTGTAAACTGACGGTG |
| CCR5-A (20 nt), for plasmids with two guide sites (site A) | CACCGCGGCAGCATAGTGAGCCCAG | AAACCTGGGCTCACTATGCTGCCGC |
