## Supplementary material for "ENHANCED CLEAVAGE OF GENOMIC *CCR5* USING CASX2^Max^": Supplmental Table S5

**Supplemental Table S5**

**Primers for PCR**

| Figure | Primer name | Primer Sequence | Amplicon |
|---|---|---|---|
| Figures 1B & 4A/B | CMV-pro_CAH | CAAGTCTCCACCCCATTGAC | Target |
| | pcDNA_CCR5_596Rev | GATAGTCATCTTGGGGCTGG | |
| Figures 1B & 4C/D | pcDNA_CCR5_596 For | CCAGCCCCAAGATGACTATC | Target |
| | CAH_bGH_Rev1 | GGAAAGGACAGTGGGAGTGG | |
| Figures 1C, 3C & 6A | CAH_CCR5_F5 | GGACAGGGAAGCTAGCAGC | Full-length genomic |
| | CAH_CCR5_R5 | CCCCATAGCAAGACAAAGACC | |
| Figure 2A | ND637 | TAATACGACTCACTATAGGG | Target |
| | ND852 | GTCCACTATTAAAGAACGTGG | |
| Figure 3A | T7_pro_for_CAH | ACGACTCACTATAGGGAGAC | Full-length target |
| | pcDNA_bgh_poly_R | GACAATGCGATGCAATTTCC | |
| Figures 4E/G & 5 | CAH_CCR5_F5 | GGACAGGGAAGCTAGCAGC | Genomic |
| | pcDNA_CCR5_596_rev | GATAGTCATCTTGGGGCTGG | |
| Figures 4F/H & 5 | pcDNA_CCR5_596 for | CCAGCCCCAAGATGACTATC | Genomic |
| | CAH_CCR5_R5 | CCCCATAGCAAGACAAAGACC | |
