## Supplementary material for "ENHANCED CLEAVAGE OF GENOMIC *CCR5* USING CASX2^Max^": Supplmental Table S6

### Supplemental Table S6

**PCR Conditions**

Figures 1B, 4 & 5

| STEP | | TEMP | TIME |
|---|---|---|---|
| Initial denaturation | | 98°C | 30 sec |
| Denaturation | | 98°C | 10 sec |
| Annealing | **repeat 34x** | 66°C | 20 sec |
| Extension | | 72°C | 30 sec |
| Final Extension | | 72°C | 2 min |
| Hold | | 12°C | |

Figure 1C, 3C & 6A

| STEP | | TEMP | TIME |
|---|---|---|---|
| Initial denaturation | | 98°C | 1 min |
| Denaturation | | 98°C | 10 sec |
| Annealing | **repeat 34x** | 67°C | 20 sec |
| Extension | | 72°C | 45 sec |
| Final Extension | | 72°C | 5 min |
| Hold | | 12°C | |

Figure 2A

| STEP | | TEMP | TIME |
|---|---|---|---|
| Initial denaturation | | 98°C | 30 sec |
| Denaturation | | 98°C | 10 sec |
| Annealing | **repeat 34x** | 58°C | 20 sec |
| Extension | | 72°C | 45 sec |
| Final Extension | | 72°C | 5 min |
| Hold | | 12°C | |

Figure 3A

| STEP | | TEMP | TIME |
|---|---|---|---|
| Initial denaturation | | 98°C | 1 min |
| Denaturation | | 98°C | 10 sec |
| Annealing | **repeat 34x** | 63°C | 20 sec |
| Extension | | 72°C | 45 sec |
| Final Extension | | 72°C | 5 min |
| Hold | | 12°C | |
